# Evidence-Gated Plasticity and Context-Dependent Consolidation for Multi-Goal Learning in Spiking Neural Networks

**DOI:** 10.64898/2026.06.25.734613

**Authors:** Samuel A Neymotin, Hananel Hazan, Gozde Unal, Christopher Earl, Haroon Anwar, Piotr J Franaszczuk, David Boothe

**Affiliations:** Center for Biomedical Imaging and Neuromodulation, Nathan S. Kline Institute for Psychiatric Research, Orangeburg, NY, USA; Dept. Psychiatry, NYU Grossman School of Medicine, New York, NY, USA; Allen Discovery Center, Tufts University, Medford, MA, USA; Okinawa Institute of Science and Technology, Okinawa, Japan; DEVCOM US Army Research Lab, Aberdeen, MD, USA; Dept. Neurology, Johns Hopkins University, Baltimore, MD, USA

## Abstract

**Background / Introduction:** Biologically inspired spiking neural networks offer a mechanistic framework for adaptive behavior, yet learning multiple goals remains challenging when synaptic updates for different targets interfere. We investigated whether multi-timescale plasticity and context-dependent consolidation improve interleaved multi-goal learning in a spiking navigation system inspired by entorhinal-hippocampal circuitry.

**Methods:** We introduce evidence-gated plasticity (EGP), in which STDP-based reinforcement learning (STDP/RL) generates transient synaptic proposals and need-weighted behavioral evidence modulates their slower consolidation into durable weights. Global EGP pools these proposals and the corresponding evidence across goals. Target-context EGP extends this framework through goal-specific proposal and evidence stores and additional need-dependent scaling of consolidation. We compared the algorithms across paired spiking networks, tested learned weights with plasticity disabled, and examined upstream representations and consolidation components.

**Results:** Global EGP supported multi-goal learning and durable navigation, while target-context EGP improved reward acquisition across replicated network comparisons and increased target-contact success during extended training. Independent evaluation confirmed that both algorithms retained navigation competence when plasticity was disabled. Additional controls suggested that overall consolidation strength contributes to the target-context reward advantage. In matched short-training experiments, conventional STDP/RL acquired shaped reward more rapidly, whereas EGP networks achieved higher target-contact success with learned synaptic weights held fixed.

**Conclusions:** EGP supports durable multi-goal learning in spiking neural networks, and its target-context extension improves acquisition and conditional target selectivity in this interleaved navigation task. The framework offers a basis for testing context-sensitive, evidence-modulated plasticity in other multi-context learning problems.

## Introduction

Animals navigate by combining spatial representations, sensory information, goal signals, and reinforcement feedback[1, 2, 3, 4]. This ability depends on interactions among the entorhinal cortex, hippocampus, basal ganglia, and motor systems. Entorhinal grid cells provide a metric-like representation of space[5], while hippocampal place cells encode specific locations and support memory-guided behavior[6, 7, 8]. Together, these circuits enable animals to learn and execute routes to behaviorally relevant goals[9, 10, 11]. Understanding how such representations can drive adaptive behavior remains an important challenge for both computational neuroscience and biologically inspired artificial intelligence.

A large body of work has examined how grid-cell and place-cell representations emerge from recurrent dynamics, oscillatory mechanisms, attractor networks, sensory inputs, and learning [12, 13, 14, 15, 16, 17, 18]. Other modeling approaches have linked hippocampal spatial representations to reinforcement learning, planning, and navigation decisions, including models incorporating successor representations and hippocampal-striatal interactions [4]. In parallel, STDP-based reinforcement learning has been studied in spiking networks [19, 20] and applied to closed-loop sensorimotor and goal-directed motor tasks [21, 22, 23]. Building on these complementary lines of research, we investigate how spatial and goal representations can support navigation in a spiking network where neural activity, action selection, environmental feedback, and synaptic plasticity continuously interact.

Spiking neural networks (SNNs) provide a useful framework for addressing this problem because they explicitly connect neural activity, synaptic plasticity, and behavior. Spike-timing-dependent plasticity (STDP) offers a biologically plausible mechanism for modifying synapses according to the relative timing of pre- and postsynaptic activity[24, 25, 26]. When modulated by reward signals, STDP can support reinforcement learning in spiking circuits. Studies using STDP-based reinforcement learning (STDP/RL) have shown that spiking networks can learn sensorimotor tasks and goal-directed actions[19, 20, 21, 22, 23]. In spatial navigation models, STDP/RL can modify projections from spatial representations to motor circuits, allowing an agent to learn movements that approach a target.

In previous work, we developed spiking navigation models that coupled biologically inspired spatial representations to reinforcement-modulated plasticity[27]. That distinct value-guided approach illustrates how local learning can support navigation. A related challenge arises when multiple goals must be learned within a shared network: synaptic updates associated with one target can interfere with updates associated with another. When spatial, association, and motor representations support multiple navigation policies, reward-modulated updates can compete across goal contexts and disrupt previously learned behavior. Such interference motivates the present biologically inspired spiking framework[28, 29].

Biological learning systems appear to address similar challenges through plasticity mechanisms operating over multiple timescales[30, 31]. Rapid STDP-like processes can encode recent activity patterns, whereas slower consolidation mechanisms determine which changes are retained[32]. Neuromodulatory signals, replay, synaptic tagging, sleep-dependent consolidation, and context-dependent gating have all been implicated in memory stabilization[33, 34, 35, 36]. Motivated by these observations, we hypothesized that multi-goal learning could benefit from accumulating candidate synaptic changes, evaluating their behavioral consequences, and consolidating them in a context-dependent manner rather than committing every update immediately.

Here we introduce evidence-gated plasticity (EGP), a multi-timescale learning algorithm for a shared spiking network. Throughout the manuscript, EGP without a qualifier denotes the foundational global rule. STDP/RL generates local synaptic changes; global EGP accumulates them as transient proposals and uses need-weighted behavioral evidence to modulate slower durable consolidation. Target-context EGP extends this process by preserving goal-specific proposals and evidence and additionally scaling each context-specific consolidation coefficient by target need. Together, these algorithms link rapid local plasticity to evidence-dependent stabilization of learned navigation behavior, with the target-context variant additionally modulating consolidation according to goal-specific information. We first characterize the spatial and target-coordinate representations, then demonstrate STDP/RL navigation, compare the two EGP variants, evaluate durable weights, and examine consolidation components and a matched STDP/RL benchmark.

Several complementary approaches place this proposal in context. Multi-timescale synaptic models address memory persistence through interacting internal states [37]. Elastic weight consolidation and synaptic intelligence stabilize parameters according to their importance for previous tasks [38, 39], while context-dependent gating modulates the neural resources recruited by a task [29, 40]. Bayesian continual learning and task-agnostic methods have also been developed for spiking networks [41, 42]. EGP complements these approaches by testing whether transient local plasticity can be expressed during behavior and consolidated according to subsequent behavioral evidence. The present study tests this framework during recurring, interleaved multi-goal learning with explicit goal identities; sequential learning without rehearsal remains an important direction for future work.

## Methods

### Simulation environment and closed-loop navigation task

All simulations were implemented in Python using the NEURON simulation environment with NetPyNE for network specification, simulation control, and data output[43, 44]. The spiking neural network was coupled to a two-dimensional navigation environment in which an agent navigated a bounded arena toward one or more target locations. At each behavioral step, the environment provided the model with spatial and target-related inputs, the spiking network generated activity in motor-output populations, and the relative activity of those populations determined the agent’s next movement direction. The resulting state transition and reward were then used to update synaptic plasticity according to the learning rule used in that simulation.

The environment consisted of a two-dimensional square arena of size 100 x 100 pixels with 8 pixel padding to prevent edge artifacts. The agent’s position was represented by x- and y-coordinates. Targets were located at (24,24), (75,75), (24,75), (75,24), and (50,50). Single-target simulations used one fixed target, whereas multi-target simulations sequentially activated different targets across subepisodes. Each episode duration depended on simulation type and learning algorithm (see **Training Protocols**), and performance was quantified using target contacts, cumulative reward, movement trajectories, and dwell time near target locations.

At each simulation step, the agent selected movement independently on the horizontal and vertical axes by comparing opposing motor populations. The resulting action could be one of eight moves or no movement; a tie produced no movement on the corresponding axis. Attempted positions were clipped to x,y∈[8,91]. A target hit was recorded when the agent was positioned directly on the active target. Movement toward or away from the goal supplied dense geometric feedback, with an additional contact bonus in the critic used for plasticity (described further below).

### Event-based integrate-and-fire neuron model

Individual neurons were event-driven integrate-and-fire point-neuron models adapted from prior spiking network models used for sensorimotor reinforcement learning[21, 22, 45]. Each neuron had a membrane voltage state variable, a resting membrane potential, a spike threshold, an absolute refractory period, and additional state variables implementing after-hyperpolarization and relative refractoriness. When synaptic input events arrived, they changed postsynaptic voltage according to the synapse type and synaptic weight. If the membrane voltage crossed threshold and the neuron was not refractory or in depolarization blockade, the neuron emitted a spike and entered an absolute refractory period.

After each spike, a negative after-hyperpolarization contribution was increased in magnitude and then decayed exponentially back toward zero. A relative refractory mechanism transiently increased spike threshold following a spike and decayed back to baseline with its own time constant. A depolarization block threshold prevented firing when the membrane voltage exceeded a high-voltage blockade value. This rule-based event-driven formulation allowed efficient simulation of large spiking networks while retaining several qualitative features of biological neurons, including refractoriness, spike-frequency adaptation, and depolarization block.

Each neuron also contained synaptic voltage variables corresponding to AMPA-like, NMDA-like, and GABAA-like synaptic inputs. Synaptic input events caused stepwise changes in these synaptic variables, which then contributed to the membrane voltage and decayed exponentially with synapse-specific time constants. AMPA-like inputs decayed rapidly, NMDA-like inputs decayed more slowly, and GABAA-like inputs provided inhibitory voltage-state effects with voltage-dependent increments. Synaptic delays were sampled from 1.8-2.2 and 2-4 ms depending on synapse type and projection class.

Neuron and synapse parameters followed prior event-driven sensorimotor models [21, 22] and were adjusted to produce stable baseline firing in the present architecture. Excitatory, fast-spiking inhibitory, and low-threshold inhibitory neurons used separate parameter sets for resting voltage, spike and blockade thresholds, refractoriness, and after-hyperpolarization (**Table 1**).

State variables were updated analytically at incoming and self-generated events; the equations and spike/reset conditions are given in Supplementary Methods **S1**. The resulting point-neuron formulation captures selected firing properties without requiring detailed biophysical reconstructions.

**Table 1.**
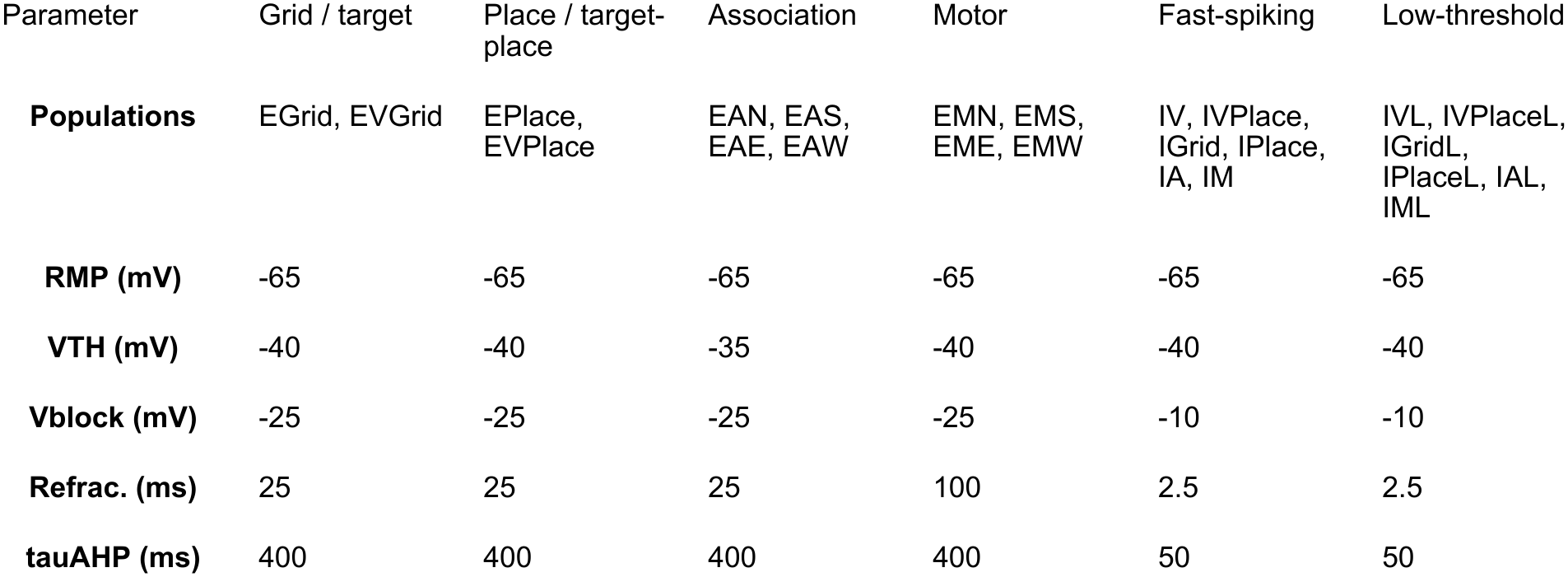
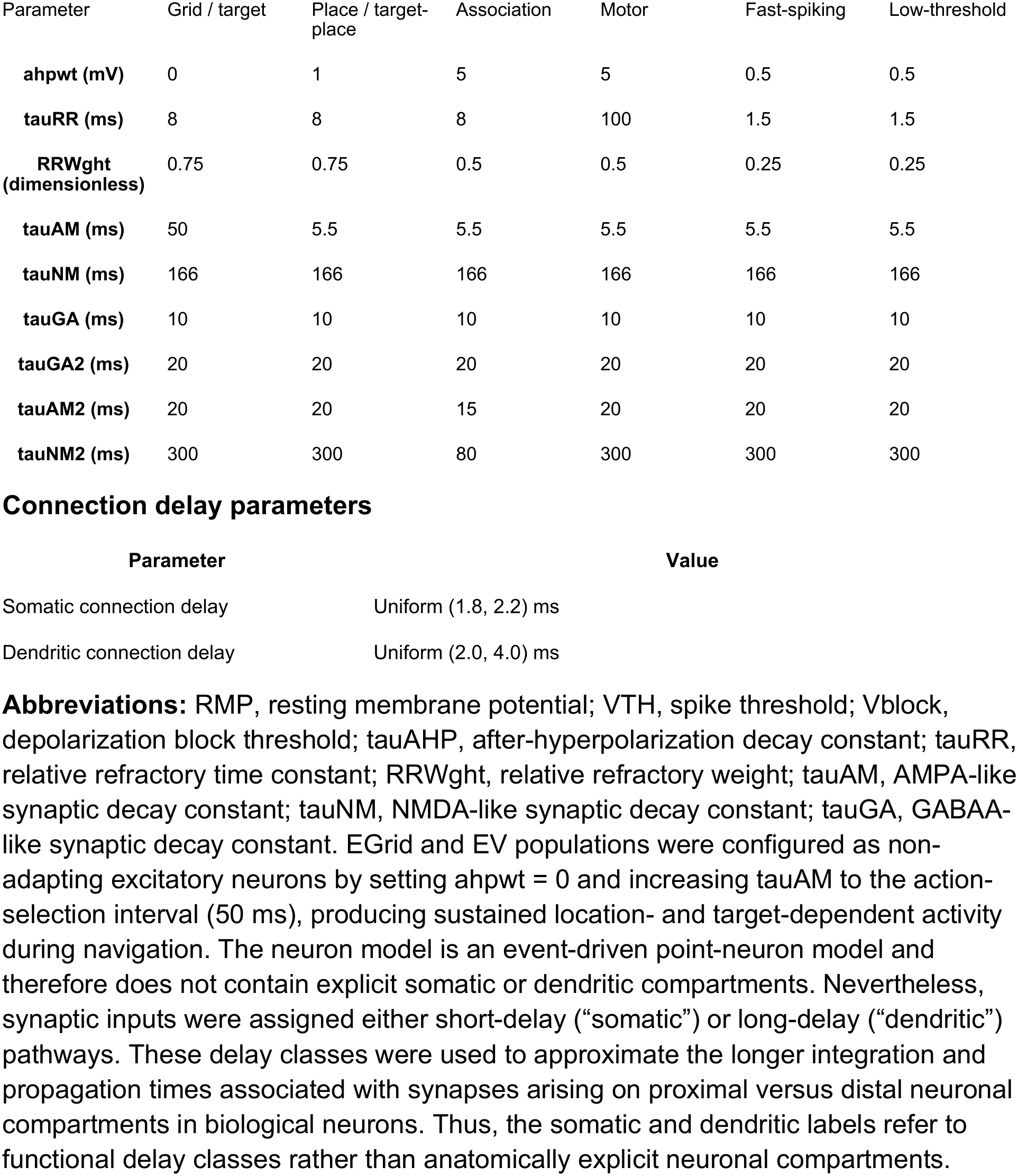
Event-based integrate and fire neuron model parameters. In excitatory association (EA) and motor-output (EM) population names, N, S, E, and W denote north, south, east, and west.

| Parameter | Grid / target | Place / target-place | Association | Motor | Fast-spiking | Low-threshold |
|---|---|---|---|---|---|---|
| **Populations** | EGrid, EVGrid | EPlace, EVPlace | EAN, EAS, EAE, EAW | EMN, EMS, EME, EMW | IV, IVPlace, IGrid, IPlace, IA, IM | IVL, IVPlaceL, IGridL, IPlaceL, IAL, IML |
| **RMP (mV)** | -65 | -65 | -65 | -65 | -65 | -65 |
| **VTH (mV)** | -40 | -40 | -35 | -40 | -40 | -40 |
| **Vblock (mV)** | -25 | -25 | -25 | -25 | -10 | -10 |
| **Refrac. (ms)** | 25 | 25 | 25 | 100 | 2.5 | 2.5 |
| **tauAHP (ms)** | 400 | 400 | 400 | 400 | 50 | 50 |
| Parameter | Grid / target | Place / target-place | Association | Motor | Fast-spiking | Low-threshold |
| **ahpwt (mV)** | 0 | 1 | 5 | 5 | 0.5 | 0.5 |
| **tauRR (ms)** | 8 | 8 | 8 | 100 | 1.5 | 1.5 |
| **RRWght (dimensionless)** | 0.75 | 0.75 | 0.5 | 0.5 | 0.25 | 0.25 |
| **tauAM (ms)** | 50 | 5.5 | 5.5 | 5.5 | 5.5 | 5.5 |
| **tauNM (ms)** | 166 | 166 | 166 | 166 | 166 | 166 |
| **tauGA (ms)** | 10 | 10 | 10 | 10 | 10 | 10 |
| **tauGA2 (ms)** | 20 | 20 | 20 | 20 | 20 | 20 |
| **tauAM2 (ms)** | 20 | 20 | 15 | 20 | 20 | 20 |
| **tauNM2 (ms)** | 300 | 300 | 80 | 300 | 300 | 300 |
| **Connection delay parameters** |  |  |  |  |  |  |
| **Parameter** | **Value** |  |  |  |  |  |
| Somatic connection delay | Uniform (1.8, 2.2) ms |  |  |  |  |  |
| Dendritic connection delay | Uniform (2.0, 4.0) ms |  |  |  |  |  |

**Abbreviations:** RMP, resting membrane potential; VTH, spike threshold; Vblock, depolarization block threshold; tauAHP, after-hyperpolarization decay constant; tauRR, relative refractory time constant; RRWght, relative refractory weight; tauAM, AMPA-like synaptic decay constant; tauNM, NMDA-like synaptic decay constant; tauGA, GABAA-like synaptic decay constant. EGrid and EV populations were configured as non-adapting excitatory neurons by setting ahpwt = 0 and increasing tauAM to the action-selection interval (50 ms), producing sustained location- and target-dependent activity during navigation. The neuron model is an event-driven point-neuron model and therefore does not contain explicit somatic or dendritic compartments. Nevertheless, synaptic inputs were assigned either short-delay (“somatic”) or long-delay (“dendritic”) pathways. These delay classes were used to approximate the longer integration and propagation times associated with synapses arising on proximal versus distal neuronal compartments in biological neurons. Thus, the somatic and dendritic labels refer to functional delay classes rather than anatomically explicit neuronal compartments.

### Spiking network architecture

The model consisted of excitatory spatial representation populations, target/context populations, an association layer, motor-output populations, and inhibitory populations used to regulate network excitability (**Fig. 1**, **Table 2**). The architecture implements a simplified entorhinal-hippocampal-to-motor transformation, using functional rather than physiologically validated population labels. Spatial inputs were represented by grid-like and place-like populations, target information was represented by target-encoding populations, and downstream association populations transformed spatial and target context into direction-specific motor drive.

**Figure 1.**
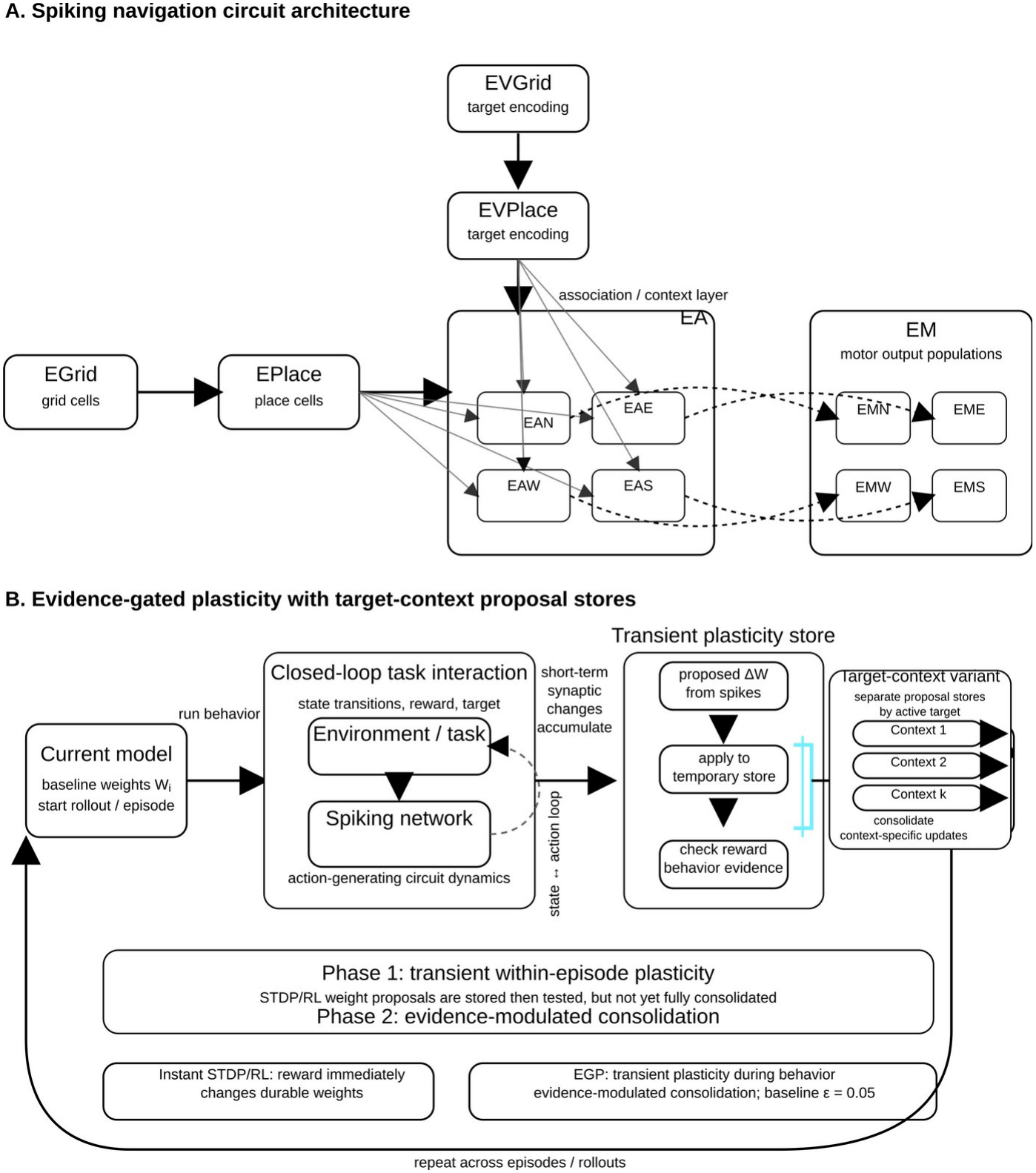
Spiking navigation circuit architecture and evidence-gated plasticity learning framework. **A)** Schematic of the spiking neural network architecture used for closed-loop spatial navigation. The agent’s current position is represented by grid-cell-like and place-cell-like populations (EGrid and EPlace). The currently active target is represented by target-encoding EVGrid (called EV in the implementation and firing-map analyses) and EVPlace populations. Spatial and target-context signals converge onto the excitatory association (EA) layer, with directional subpopulations labeled N (north), E (east), W (west), and S (south). Each EA subpopulation projects to the corresponding excitatory motor-output (EM) population: EAN→EMN, EAE→EME, EAW→EMW, and EAS→EMS. Activity in the EM populations determines the agent’s movement direction, allowing spatial and target-context representations to be transformed into direction-specific motor drive. B) Schematic of evidence-gated plasticity (EGP). During each rollout, the spiking network interacts with the environment in a closed loop: environmental state and target information drive network activity, motor-output activity selects actions, and the environment returns state transitions and reward. In standard online STDP/RL, reward-modulated spike-timing events immediately modify durable synaptic weights. In EGP, these spike- and reward-dependent weight changes are first accumulated as transient synaptic proposals. During TEST, the proposals are temporarily expressed together; the resulting behavioral evidence then modulates their subsequent consolidation. A baseline factor ε=0.05 allows limited consolidation without positive evidence; durable weights are restored before commitment, and TEST-only proposals are discarded. In the target-context variant, transient proposal stores are separated by active target context before consolidation, allowing target-specific evidence and need weights to modulate commitment within one shared network.

The spatial input pathway included an excitatory grid-cell-like population, EGrid, and a downstream place-cell-like population, EPlace. EGrid encoded the agent’s current position using periodic two-dimensional spatial fields. For each EGrid neuron, a grid scale was sampled uniformly between 5 and 10 pixels. The grid spacing was then sampled as an integer between 5 and 7 times that scale, and the spatial phase offset was assigned by randomizing the starting x and y coordinates within the 100 × 100 environment. Grid fields were not restricted to a fixed set of prime scales or predefined modules. Instead, each neuron received an independently sampled periodic field. Grid fields were generated as hexagonally arranged Gaussian-like disks, with adjacent rows shifted by half the grid spacing and separated vertically by 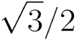 of the spacing.

Target-coordinate input was represented by an analogous EVGrid population (EV in the implementation), which encoded the currently active target location using the same grid-map generation procedure. Thus, EGrid represented the agent’s current location, whereas EVGrid represented the current goal location in a comparable grid-like coordinate system. EGrid projected to EPlace, and EVGrid projected to EVPlace, producing downstream localized spatial and target-related representations. EPlace and EVPlace then projected to directional association populations, where current-location and target-location information could be integrated to guide motor-output activity.

The association layer consisted of excitatory association neurons (EA), subdivided into action-specific context populations: EAN, EAS, EAE, and EAW. These subpopulations were action-pure in the sense that each projected preferentially to the corresponding motor-output subpopulation. EAN projected to EMN, EAS to EMS, EAE to EME, and EAW to EMW. The motor layer consisted of excitatory motor-output populations encoding north, south, east, and west actions. At each behavioral step, spikes in the four EM populations were counted over the action-selection interval, and opposing populations were compared independently on each axis, as detailed below. A tie prevented movement only on that axis.

Inhibitory populations were included to prevent runaway excitation and to regulate firing rates in association and motor layers. Excitatory synapses were mediated by AMPA- and NMDA-like mechanisms, whereas inhibitory synapses were mediated by GABAA-like mechanisms. Plasticity was applied to EA→EM sub-populations. Unless otherwise stated, plasticity was limited to excitatory synapses onto excitatory postsynaptic cells.

**Table 2.** Network populations and configured input convergence.

| Population | Neurons | Inputs/ neuron | Source populations, convergence per postsynaptic neuron |
|---|---|---|---|
| EVGrid | 384 | 29 | IV (16); IVL (13) |
| IV | 88 | 101 | EVGrid (60); IV (22); IVL (19) |
| IVL | 40 | 60 | EVGrid (45); IV (12); IVL (3) |
| EVPlace | 384 | 69 | EVGrid (19); IVPlace (30); IVPlaceL (20) |
| IVPlace | 88 | 102 | EVPlace (60); IVPlace (22); IVPlaceL (20) |
| IVPlaceL | 40 | 67 | EVPlace (45); IVPlace (12); IVPlaceL (10) |
| EGrid | 384 | 50 | IGrid (30); IGridL (20) |
| IGrid | 88 | 102 | EGrid (60); IGrid (22); IGridL (20) |
| IGridL | 40 | 67 | EGrid (45); IGrid (12); IGridL (10) |
| EPlace | 384 | 69 | EGrid (19); IPlace (30); IPlaceL (20) |
| IPlace | 88 | 102 | EPlace (60); IPlace (22); IPlaceL (20) |
| IPlaceL | 40 | 67 | EPlace (45); IPlace (12); IPlaceL (10) |
| EAN | 768 | 220 | EVPlace (10); EPlace (10); IA (120); IAL (80) |
| EAS | 768 | 220 | EVPlace (10); EPlace (10); IA (120); IAL (80) |
| EAE | 768 | 220 | EVPlace (10); EPlace (10); IA (120); IAL (80) |
| EAW | 768 | 220 | EVPlace (10); EPlace (10); IA (120); IAL (80) |
| IA | 704 | 416 | EAN (60); EAS (60); EAW (60); EAE (60); IA (88); IAL (80); EMW (2); EME (2); EMS (2); EMN (2) |
| IAL | 320 | 280 | EAN (46); EAS (46); EAW (46); EAE (46); IA (48); IAL (40); EMW (2); EME (2); EMS (2); EMN (2) |
| EMN | 192 | 88 | EAN (10); IM (57); IML (21) |
| EMS | 192 | 88 | EAS (10); IM (57); IML (21) |
| EME | 192 | 88 | EAE (10); IM (57); IML (21) |
| EMW | 192 | 88 | EAW (10); IM (57); IML (21) |
| IM | 176 | 356 | EMW (61); EME (61); EMS (61); EMN (61); IM (80); IML (32) |
| IML | 80 | 339 | EMW (72); EME (72); EMS (72); EMN (72); IM (44); IML (7) |

**Note:** Only populations with nonzero cell counts are shown. Convergence values indicate the intended number of presynaptic inputs from each source population onto each postsynaptic neuron.

### Background synaptic noise

Selected network populations received independent Poisson-distributed background synaptic input to maintain spontaneous activity and promote behavioral exploration. Noise was implemented using Poisson sources that generated AMPA-like excitatory input at 200 Hz and weaker GABA-like inhibitory input at 100 Hz. These inputs were delivered through fixed-weight synapses and were not subject to learning. Spatial and target-representation populations (EGrid, EVGrid, EPlace, and EVPlace), association populations (EA subpopulations), and most inhibitory populations were excluded from direct noise input. Consequently, background noise primarily influenced downstream action-selection and motor-output populations, providing variability in behavioral choices while leaving the spatial and target representations largely determined by external inputs and network interactions.

### Spatial and target input encoding

At each behavioral step, the agent’s current position determined the subset of active EGrid neurons. Active neurons received position-dependent stimulation with a peak firing rate of approximately 20 Hz. EPlace neurons received convergent input from EGrid and generated spatially selective activity patterns, which we characterize directly using the spike-derived maps described below. An analogous target-representation pathway, consisting of EVGrid and EVPlace populations, encoded the currently active goal location rather than the agent’s position. During multi-target training, the active target changed according to the training schedule, allowing EVGrid and EVPlace activity to provide downstream association populations with goal-specific context information.

This target-context signal contributed to action selection and, in the target-context EGP model, was also used to separate accumulated synaptic proposals according to the currently active goal.

### Action selection

At each environment step, the network was simulated for 50 ms after spatial and target inputs were delivered. Spikes were counted in each motor-output population over this interval. Movement along the horizontal axis was determined by comparing activity in the eastward (EME) and westward (EMW) motor populations, whereas movement along the vertical axis was determined by comparing activity in the northward (EMN) and southward (EMS) populations. If one population in a pair produced more spikes than its opponent, the agent moved one step in the corresponding direction. If both populations in a pair produced identical spike counts, including zero spikes, no movement occurred along that axis. Northward movement decreased y. Thus, both axes could change during the same interval, yielding diagonal as well as cardinal movements. The selected action was sent to the environment, which updated the agent’s position and returned the next state and reward.

### STDP-based reinforcement learning (STDP/RL)

STDP-based reinforcement learning (STDP/RL) directly updated synapses during online learning and generated the transient candidate changes used by EGP [21, 22]. Spike-pair timing establishes an eligibility trace that a subsequent scalar critic signal converts into potentiation or depression. A postsynaptic spike within 100 ms after presynaptic activity tagged the synapse with a 50-ms square-pulse eligibility trace, allowing reinforcement to modify recently active connections.

For each eligible synapse, the reward-modulated weight change was computed as a product of the eligibility trace, a learning-rate parameter, and the critic/reward signal: *Δw_ij_* (*t*)=*η_STDP_ ∗ e_ij_* (*t*)*∗ r* (*t*) where *w_ij_* is the synaptic weight from presynaptic neuron i to postsynaptic neuron j, *e_ij_* (*t*) is the eligibility trace, *r* (*t*) is the critic or reward signal, and *η_STDP_* is the learning-rate parameter. Positive reward increased eligible synaptic weights, whereas negative reward decreased them. Clipping is defined as clip(x,a,b)=min(b,max(a,x)); weights were clipped to the instantiated synaptic bounds. The base AMPA learning increment and bounds in Table 3 are multiplied by the excitatory projection gain before use; in the controlled EGP configurations the gain was 4.4, giving increment 0.198 and bounds 0-52.8. Plasticity was applied every 50 ms during online learning.

The critic signal used for plasticity was derived from, but distinct from, the raw reward reported as a behavioral outcome. In single-target and multi-target STDP/RL simulations, the critic rewarded movements that improved navigation relative to the active target and penalized movements that increased distance from the target.

The movement component of the critic depended on whether the selected action moved the agent toward or away from the active target, with an additional bonus for target contact. The movement reward was defined as: *r_move_* (*t*)=+1 if the action reduced distance to the target, *r_move_* (*t*)=*−* 1 if the action increased distance to the target, *r_move_* (*t*)=0, if the distance to the target did not change.

Because reward scaling by distance was enabled, movement rewards were amplified near the target using a smooth distance-dependent gain: 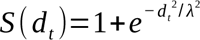, with *d* the distance from the boundary-clipped position to the target, and *λ*=8. The critic signal applied to reward-modulated plasticity was therefore *R* (*t*)=*r_move_* (*t*) *S* (*d_t_*)+*r_hit_* 1*_hit_* (*t*) , where *r_hit_* =2, and 1*_hit_* (*t*)=1 when the agent was on the active target and 0 otherwise.

Plasticity was restricted to action-specific association-to-motor projections: EAN→EMN, EAS→EMS, EAE→EME, and EAW→EMW. For each plastic synapse (*w_ij_*), causal pre-before-post spike timing generated an eligibility trace, implemented as a fixed-duration square pulse. When a presynaptic spike preceded a postsynaptic spike within 100 ms, 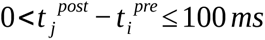, the synapse entered an eligible state for a duration of 50 ms. During this interval, subsequently arriving reward signals could modify the synaptic weight.

The raw environment reward was the decrease in Euclidean distance produced by the attempted movement, divided by the simulator’s fixed distance-normalization scale, D_norm_:

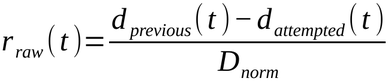

Here W=H=100 pixels are the configured arena dimensions, and D_norm_=√(W²+H²) is a fixed normalization scale, not the maximum reachable distance within the padded arena. The attempted position is evaluated before boundary clipping. Its reward sign defines the movement component of the critic, whereas the distance gain and exact-contact bonus use the clipped position. Consequently, the reported raw reward is neither the signed critic nor a target-hit count. The same dense geometric feedback is available throughout learning; these experiments do not test sparse-reward navigation.

For eligible synapses, the base reward-modulated weight change was *Δwᵢⱼ*= *A ₊R* (*t*), where R(t) is the critic signal and A₊=0.045 before the projection gain. Direction-specific reinforcement factors are applied as described next.

Eligibility tags were reset on reward delivery at the 50-ms action interval, ensuring that reinforcement was assigned only to recent spike-pair events. In the target-specific motor learning rule, synapses contributing to the selected movement direction received the signed critic signal, whereas synapses associated with the opposing movement direction received the opposite sign scaled by 0.25. Candidate updates generated by this rule were either applied immediately in STDP/RL simulations or accumulated as proposals in evidence-gated plasticity simulations described below.

The rule was designed to enable the network to learn associations between spatial/target inputs, association-layer activity, and motor-output populations. In multi-target simulations, however, the same shared plastic synapses were modified across different target contexts, creating the possibility of interference between target-specific policies.

**Table 3.**
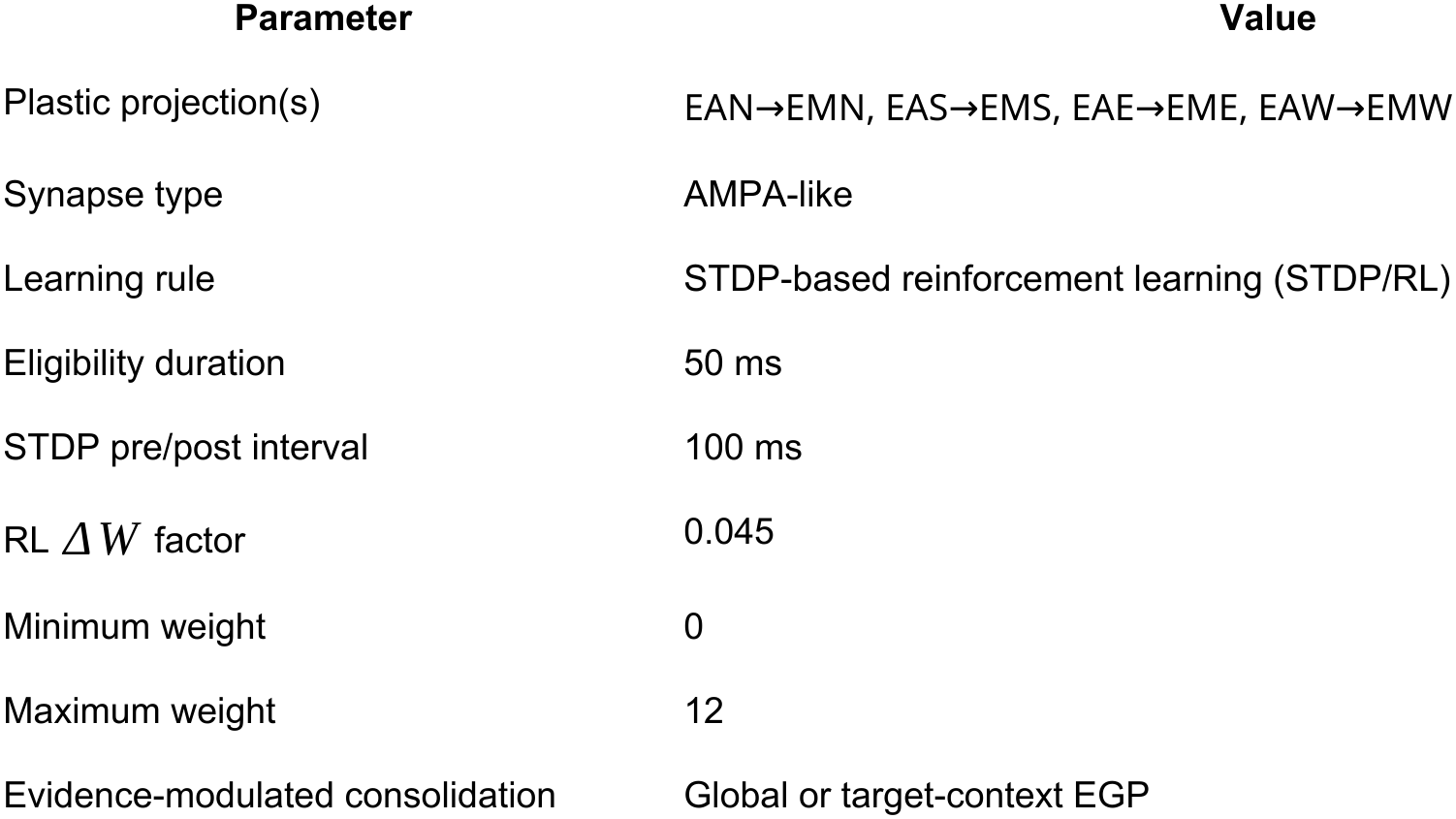
Plastic synapses and base learning parameters. Weight-valued entries precede the projection gain; see text for the instantiated EGP values.

| **Parameter** | **Value** |
|---|---|
| Plastic projection(s) | EAN→EMN, EAS→EMS, EAE→EME, EAW→EMW |
| Synapse type | AMPA-like |
| Learning rule | STDP-based reinforcement learning (STDP/RL) |
| Eligibility duration | 50 ms |
| STDP pre/post interval | 100 ms |
| RL *ΔW* factor | 0.045 |
| Minimum weight | 0 |
| Maximum weight | 12 |
| Evidence-modulated consolidation | Global or target-context EGP |

### Need-weighted evidence and consolidation

Target need was determined from recent performance. For each target k, the estimate qₖ was initialized from its first observed phase mean of raw environment reward, r̄k, and subsequently updated as 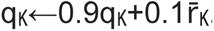. Unseen targets had need nₖ=1. Observed targets received larger need weights when their recent reward was lower:

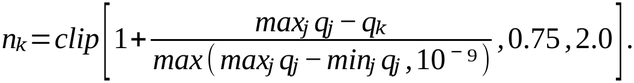

In global EGP, nₖ weights the critic samples entering the single TRAIN/TEST evidence calculation. Target-context EGP uses the same need-weighted samples in each target’s evidence and additionally multiplies that target’s consolidation coefficient by nₖ. Thus the two rules differ in proposal/evidence organization and in the extra coefficient multiplier. For observed targets, the implemented need equation yields values from 1 to 2, so the configured 0.75 lower bound is inactive. The best-performing target has need 1; the worst has need 2 when the performance range is at least 10⁻⁹. Equal estimates give unit need, and smaller positive ranges are limited by the denominator guard.

### Global evidence-gated plasticity

Global evidence-gated plasticity (EGP) separates rapid local plasticity from slower evidence-modulated consolidation. During TRAIN, reward-modulated changes accumulate in a transient proposal store while saved durable weights drive behavior. TEST temporarily expresses the candidate weights, and the resulting need-weighted reward evidence determines consolidation strength. After restoring the durable weights, EGP commits the scaled TRAIN proposal and resets transient state. The baseline factor ε=0.05 permits consolidation even without positive evidence.

For each plastic synapse, EGP maintained a transient proposal variable, *P_ij_*, representing the accumulated candidate weight change since the previous consolidation event. During TRAIN phases, local STDP/RL events did not immediately modify the durable synaptic weight matrix. Instead, candidate changes were accumulated according to

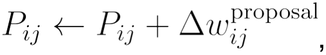

where 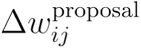 was generated using the same reward-modulated spike-timing mechanism used by standard STDP/RL. Thus, EGP preserved the local credit-assignment structure of STDP/RL while delaying long-term consolidation.

The temporary candidate weight matrix, W_candidate, is the clipped sum of the saved durable weights *W* and the TRAIN proposals, as defined below. Global EGP uses a single proposal *P*; the sum notation also covers target-context EGP, where P=ΣₖPₖ. The entire aggregate candidate remains expressed throughout TEST, including across target switches, rather than testing each target under an isolated proposal. TEST performance is compared with TRAIN to determine subsequent durable consolidation. Here clip applies elementwise at the instantiated weight bounds, 0 and 52.8.

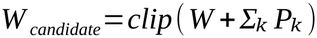

At TEST end, the saved durable W was restored before consolidation. TEST-only proposal events were discarded; only the saved TRAIN proposals were eligible for commitment. This restoration prevents the temporary candidate from being counted again in the durable update.

Reward evidence was quantified using the normalized improvement 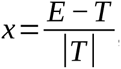, where T and E are the phase totals of the need-weighted critic during TRAIN and TEST, respectively, not the raw reward reported in the Results. If T=0, x=0 when E=0; otherwise x=E/max(|E|,10⁻⁹).

TRAIN and TEST use the same navigation task, but TEST is an internal learning phase that temporarily expresses candidate weights. At TRAIN→TEST, the agent returns to the configured TRAIN start while neural and input-noise state continue. TEST→TRAIN continues from the preceding endpoint; target switches alone do not reset position. Independent fixed-weight evaluation is described separately below.

Only positive reward improvements increased the evidence-dependent component of consolidation. EGP used the rectified sigmoid evidence function

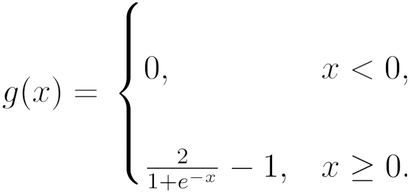

This maps positive reward improvements onto the interval [0, 1), while preventing negative reward evidence from increasing the evidence-dependent component of consolidation.

The effective consolidation coefficient was then

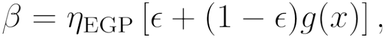

Where *n_EGP_* = 1.0 is the consolidation learning rate and *e* = 0.05 is a minimum consolidation factor (η=1 and ε=0.05). The durable synaptic weight was updated according to

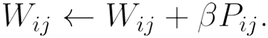

This update was clipped to the instantiated bounds after restoring the saved durable weights. Consequently, all proposals received a small baseline level of consolidation, while proposals associated with larger improvements in TEST relative to TRAIN reward received progressively stronger consolidation. After consolidation, the proposal variables were reset and a new TRAIN-TEST cycle began.

This learning rule implements a functional separation between rapid local plasticity and slower evidence-dependent consolidation. Local STDP/RL generates candidate synaptic modifications on behavioral timescales, whereas EGP evaluates their utility using reward evidence before committing them to the durable synaptic matrix. EGP therefore provides a biologically inspired computational abstraction of the distinction between rapid experience-dependent plasticity and slower stabilization processes, rather than a literal cellular implementation. Conceptually, the framework is related to synaptic tagging and capture models, in which transient activity-dependent traces are stabilized when supported by subsequent consolidation signals[46].

### Target-context evidence-gated plasticity

The target-context EGP variant was developed to retain the target identities of candidate synaptic modifications until consolidation. In global EGP, all accumulated proposals contribute to a single proposal variable (Pᵢⱼ), regardless of which target is active. Consequently, updates that improve navigation toward one target can interfere with updates that improve navigation toward another target.

To address this limitation, target-context EGP maintains separate proposal stores for each target context,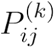 where (k) indexes the currently active target. During TRAIN phases, STDP/RL events generate candidate synaptic modifications using the same local plasticity rule as online STDP/RL, but these updates are accumulated only within the proposal buffer associated with the active target: 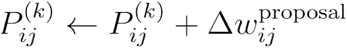.

When the active target changes during a TRAIN phase, the accumulated proposal generated under the previous target context is stored in the corresponding proposal buffer, and accumulation continues within the proposal buffer associated with the newly active target. Thus, candidate weight changes generated while pursuing different navigation goals remain separated throughout the TRAIN phase.

Each target context also maintains separate TRAIN and TEST reward estimates, 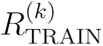 and 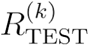These were computed as the need-weighted critic signal accrued while target (k) was active, normalized by the number of action intervals observed for that target. The context-specific reward-improvement measure was

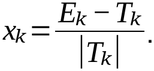

Here Tₖ and Eₖ are the mean need-weighted critic samples for target k in TRAIN and TEST. The same zero-denominator rule applies: if Tₖ=0, set xₖ=0 when Eₖ=0; otherwise use Eₖ/max(|Eₖ|,10⁻⁹).

As in global EGP, a rectified sigmoid transformed this reward-improvement measure into an evidence signal:

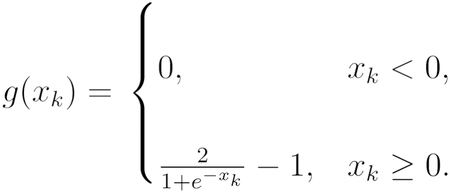

For target *k*, the need weight *nk* follows the shared update above. Target-context EGP additionally uses it to scale the context-specific consolidation coefficient:

The effective consolidation coefficient for target context *k* was therefore 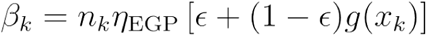, where *n_EGP_* = is the consolidation learning rate and *e* is a minimum consolidation factor. Thus, *Ꞵ*_k_ is the effective evidence- and need-weighted consolidation coefficient applied to proposals generated under target context *k*.

After restoring the saved W, the durable synaptic matrix was updated using the weighted sum of the saved TRAIN proposals, with clipping to the instantiated bounds:

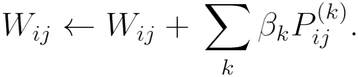

TEST-only proposals were discarded. After consolidation, all context-specific proposal buffers were reset and a new TRAIN-TEST cycle began.

Target-context EGP retains one shared network and one durable synaptic weight matrix. Goal-specific proposal stores and evidence allow proposals from each target to receive a separate consolidation coefficient before they are combined into the durable weight update. The component experiment below examines evidence organization, evidence need weighting, and the extra need multiplier. Because TEST expresses the aggregate candidate, improvement for target k can reflect proposals accumulated under other targets. Target-indexed evidence therefore conditions consolidation but does not isolate the causal behavioral contribution of proposal k.

The full sequence is therefore: accumulate proposals under W; express their clipped aggregate during TEST; restore W; calculate evidence-dependent coefficients; and commit the clipped weighted sum of the saved TRAIN proposals. Supplementary Methods S2 gives pseudocode and the state transition details. For N plastic synapses and K contexts, proposal storage scales as O(KN): with 7,680 plastic synapses, one float64 proposal vector occupies 60 KiB and five occupy 300 KiB, excluding other bookkeeping. Measured within-job wall times are reported under Computational Resources; the storage counts exclude other state.

### Training protocols

#### Single-target STDP/RL

simulations were used to determine whether the network could learn a basic navigation policy. The agent was trained to reach a single central target at (50,50) for 800 episodes, with each episode lasting 187.5 seconds of simulated time. Performance was assessed using target hits, reward, and example trajectories. To assess retention of the learned policy, we evaluated the trained network for 400 post-learning episodes with synaptic plasticity disabled; **Fig. 3B** compares the final 100 learning episodes with the first 100 post-learning episodes. At the beginning of each episode, the agent was initialized to a random starting location in the 100 x 100 pixel environment, to allow the model to learn to reach the target from an arbitrary position.

#### Multi-target STDP/RL

simulations used five fixed target locations: four quadrant targets at (24,24), (75,75), (24,75), and (75,24), and one central target at (50,50). Each episode used a single stationary target, and the active target was advanced sequentially across episodes. Starting location was randomized at the beginning of each episode. Each episode lasted 187.5 s, and the full learning phase contained 4000 episodes, corresponding to 800 episodes per target. The active target was represented by EVGrid and EVPlace input. STDP/RL remained active during the learning phase. After learning, synaptic plasticity was disabled and the network was evaluated for 500 episodes (100 per target) to determine whether target-dependent navigation policies had been retained in the learned synaptic weights. The trajectory and dwell displays use the combined first 100 learning, last 100 learning and last 100 post-learning episodes, giving 20 observations per target in each displayed stage. The target-hit display instead uses 100 observations per target per stage, as specified in its legend.

#### Short multi-seed EGP

simulations compared global EGP with target-context EGP in an initial 10-pair cohort followed by a separate, prospectively specified 30-pair replication. The algorithms differed in how proposals and evidence were organized: global EGP used one pooled proposal store, whereas target-context EGP retained separate proposal buffers and reward evidence for each target. Both EGP modes used need-based reward weighting, with need weights computed from recent target-specific performance and bounded between 0.75 and 2.0. In the target-context implementation, these need weights additionally scaled the context-specific consolidation coefficients.

These simulations used five target locations, (24,24), (75,75), (24,75), (75,24), and (50,50). The active target was updated every 150 action intervals, corresponding to 7.5 s of simulation time, because each action interval lasted 50 ms. Each TRAIN or TEST phase lasted 750 action intervals, or 37.5 s, and therefore contained five 7.5 s target blocks. Each 375 s episode contained repeated alternating TRAIN and TEST phases, and 800 episodes were run for each seed. Connectivity and spatial-input seeds varied across paired realizations; each algorithm pair shared its network and input seeds. The prospective primary endpoint was each seed’s mean TEST raw reward per target block over episodes 0-99. The initial cohort and pooled 40-pair summary provide supporting evidence, not additional independent replications.

To characterize long-timescale learning, global and target-context EGP were compared in eight paired network realizations, each trained for 2000 sequential episodes. Prior to each episode, a new starting location was selected randomly from valid locations within the environment. The same five target locations were used throughout learning. As in the short experiments, target-context EGP maintained separate proposal stores, reward estimates and consolidation coefficients, with need weighting in both evidence and the context-specific coefficient. The paired algorithms shared the architecture, initialization, goal schedule and feedback. The primary endpoint was the seed-level mean TEST raw reward per target block across all 2000 episodes; final-100 and target-specific effects were secondary.

Each episode lasted 1500 s and consisted of alternating TRAIN and TEST phases. TRAIN and TEST phases each lasted 150 s (3000 action intervals of 50 ms). Within both phases, the active target context changed every 30 s (600 action intervals), resulting in five target blocks per TRAIN phase and five target blocks per TEST phase. Consequently, each episode contained five TRAIN phases and five TEST phases, yielding 25 TRAIN target blocks and 25 TEST target blocks per episode. Under target-context EGP, candidate synaptic modifications accumulated by target during TRAIN; in both EGP variants, TEST evidence subsequently modulated consolidation into the durable synaptic weight matrix. These simulations were used to examine long-timescale reward dynamics, context-specific consolidation, trajectory evolution, and target-selective navigation behavior. Position and neural-state continuity followed the phase-transition rules above.

### Independent evaluation of learned synaptic weights

We performed a separate evaluation to distinguish competence stored in durable weights from performance during EGP’s internal TEST phase. For each of the eight paired long-training networks, we evaluated the pretraining weights and the weights after episodes 0, 99, 199, 499, 999, 1499, 1799 and 1999. All learning and proposal mechanisms were disabled, and evaluation began from fresh starts. Each checkpoint used 50 trials per algorithm and seed across the five targets, yielding 7200 trials overall. Each trial lasted at most 187.5 s (3750 actions). The prespecified primary endpoint was the paired difference in final-checkpoint exact-target success probability. Other checkpoints, targets and navigation/dwell measures were secondary and retained their separate multiplicity families. Thus, these trials directly assessed the navigation policies retained in the durable weights, independently of ongoing consolidation.

### Spike-derived spatial firing maps

We characterized the upstream representations in one fixed pretraining network (seed 1), measuring all 384 cells in each of EGrid, EPlace, EV and EVPlace. One balanced sweep varied agent position with the target fixed; a separate sweep varied target coordinates with agent position fixed. EGrid/EPlace therefore describe agent position, and EV/EVPlace target coordinates. Each of 256 eligible 5×5-pixel bins received 50 visits of 50 ms (2.5 s/bin); 144 padding bins remained unobserved.

Rates were recorded spike counts divided by occupancy. Spatial information per spike was calculated from the unsmoothed rates [47] using the equation below. Here pᵢ is the occupancy fraction of eligible bin i, λᵢ its firing rate, and 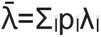 the occupancy-weighted mean rate. Zero-rate terms contribute zero when 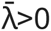; silent cells have undefined information. For the secondary reliability measure, alternating visits formed two sets of 25 visits/bin. Pearson correlation between their independently estimated, smoothed maps measures reproducibility of the spatial pattern. This is distinct from information per spike or gridness. Display/field smoothing and thresholds are specified in Supplementary Methods S2. The first 12 cell identifiers per population were selected before viewing maps; all 48 examples and all 1536 cells remain in the descriptive summaries.

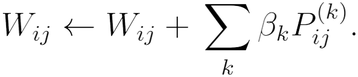

### Consolidation components and matched learning-rule comparisons

To identify the contribution of individual EGP operations, we compared four behavioral conditions across eight paired network seeds in a 200-episode experiment using 7.5-s target blocks. Global EGP (A) had a segmented global shadow (B): an identity/control calculation inside A with separate proposal bookkeeping, A’s common coefficient, and no independent motor behavior. B’s behavioral reward was A’s reward. Context-evidence EGP (C) used context-specific proposals and need-weighted evidence, but no extra need multiplier on the coefficient. Context evidence without need weighting (D) removed need from that evidence. Full target-context EGP (E) added the extra need multiplier to the context-specific coefficient. Four behavioral streams produced 6400 episodes; B was not a fifth simulated stream.

The primary outcome of the component experiment was mean raw TEST reward per target block during episodes 0–99, comparing full target-context with global EGP (E−A). Three component reward contrasts (C−B, C−D and E−C) formed one statistical family. We also examined durable-weight changes, proposal cancellation, coefficient dispersion, clipping, and navigation measures in separately specified families. Inference used paired network comparisons with Holm-Bonferroni correction within each family.

The Supplement provides the full endpoint definitions, uncertainty estimates, adjusted results, and measures unavailable because of insufficient target contacts.

To determine whether the EGP reward differences reflect changes in consolidation strength, and to compare evidence-gated learning with direct online plasticity, we extended this 200-episode framework with three consolidation controls and two STDP/RL baselines.

All eight comparative conditions used the same five recurring goals, eight network seeds, architecture, initial synaptic weights, input seeds, reward and critic signals, eligibility rules, and weight bounds. Each network completed 200 episodes of 7,500 action intervals (1.5 million interactions), with targets alternating every 7.5 s. The global (A), context-evidence (C), and full target-context (E) conditions were the network realizations described above; the remaining conditions followed the same experimental schedule. Only durable synaptic weights carried between episodes, whereas neural state, eligibility, and need estimates were reinitialized. The eight comparative conditions comprised 12,800 network episodes.

For target context k, let bₖ denote the evidence-dependent consolidation coefficient before scaling by need nₖ; full target-context EGP uses βₖ=nₖbₖ. The uniform-gain control (G) applied a common multiplier c, with βₖ=cbₖ and c=Σₖnₖbₖ/Σₖbₖ, whereas the normalized-need control (N) applied βₖ=nₖbₖ/c. These controls separate overall coefficient amplification from relative need weighting within each network’s evolving state. The shuffled-evidence control (S) reassigned the context-specific evidence coefficients among goals while retaining the target-specific need weights. Because shuffling changes the pairing between evidence and need, the final coefficient distribution need not remain identical. Supplementary Methods S10 provides the precise definitions and numerical safeguards.

Online STDP/RL (R) applied eligible synaptic changes directly to durable weights throughout training and reset the agent’s position only at episode starts. Reset-matched STDP/RL (Q) also reproduced EGP’s within-episode TRAIN→TEST position resets, but learning continued throughout the episode. Neither STDP/RL condition used transient EGP proposals, candidate expression, or evidence-gated consolidation. The algorithms received the same number of environmental interactions, although their opportunities for direct durable weight updates and the resulting synaptic changes differed. Net durable-weight changes were measured over common 75-s intervals.

To assess retained navigation, all eight conditions were evaluated after episode 199 using 50 matched fresh-start trials per network (10 per target; 187.5 s per trial), with plasticity and proposal expression disabled. The resulting 64 networks contributed 3,200 evaluation trials. Exact-target success and restricted first-passage time were summarized within each network, with seed-level results in Supplementary Table S18.

Early reward comparisons used each network’s mean raw reward per target block during episodes 0–99. Three separately adjusted families examined (i) E−G, E−N and E−S using internal TEST reward, (ii) E−R, A−R, E−Q and A−Q using reward across all interactions, and (iii) E−C, E−A and N−C using internal TEST reward. Two-sided exact paired sign-flip tests were performed across the eight matched network seeds, with Holm correction within each family. The E−C and E−A comparisons use the component-study networks; late learning curves, synaptic diagnostics, and independent frozen-weight navigation were analyzed descriptively (Supplementary Methods S10 and Table S17).

#### Behavioral and statistical analyses

Performance was quantified using complementary reward, contact, trajectory and dwell measures. Reported reward is cumulative raw environment reward normalized per target block, not the critic signal used for plasticity. For the final 100 episodes (1900– 1999) of the 2,000-episode EGP experiment, the analysis included 2,500 TEST target blocks per network and condition (40,000 in total). Network seed was the unit of inference in paired EGP comparisons; episodes, target blocks and neurons were not treated as independent network replicates. The prospectively specified primaries used two-sided exact paired sign-flip tests, with all 2³⁰ assignments for the 30-pair replication and all 256 assignments for eight-pair experiments. Percentile 95% confidence intervals used 100,000 paired seed-level bootstrap samples. Intervals are unadjusted estimation summaries. Secondary families used Holm-Bonferroni adjustment as specified for each experiment, with α=0.05. Curve smoothing was for display only: 25-episode windows for replicated EGP curves and a 100-episode window for the single-target STDP/RL illustration. Shading denotes the between-seed standard error of the mean (SEM) for replicated curves, not confidence intervals or variability over independent trials.

Target-specific performance was assessed by computing late-stage reward for each target over the final 100 episodes. Weakest-target reward was defined as the minimum target-specific reward for a given seed. The fraction of targets with positive late-stage reward was used as a reward-based summary, not as a count of targets successfully reached. Exact target contact was assessed separately.

To quantify target selectivity, we computed dwell-time confusion matrices. For each active target, we measured the fraction of time the agent spent within radius r (5 pixels) of each possible target location. Matrix rows corresponded to the active target, and columns corresponded to the target location near which the agent was located. Diagonal entries quantify dwell near the correct target, whereas off-diagonal entries quantify occupancy near inactive targets. Summary metrics included mean diagonal dwell, weakest-target diagonal dwell, maximum off-diagonal dwell, and the ratio of mean diagonal to mean off-diagonal dwell. Conditional wrong-target dwell is wrong-target dwell divided by total dwell near any target; raw wrong-target dwell instead uses all observed time as its denominator. These quantities can move differently, so both are reported.

Trajectory plots illustrate behavior during early learning, late learning, and post-learning evaluation. STDP/RL plots use the early, late, and post-learning windows defined above. For EGP, seed 1 and episode 1999 were selected in advance for visualization, with all five targets and all five TRAIN/TEST segments retained for both algorithms. For each trajectory, the active target was indicated separately from inactive targets, and the agent’s position over time was color coded to show movement through the episode.

Navigation success meant at least one exact target contact within the block or trial. First-passage time was measured to the first contact and censored at the full horizon for failures; restricted mean first-passage time includes those failures. Reached-only time and path inefficiency were reported separately because they condition on success. Zero shortest-path denominators and absent reached trials remained undefined. In the ablation, all eight paired seeds were required for an inferential test; four path comparisons were unavailable because one condition had no early reached trials for a seed. These endpoints were retained in their prespecified multiplicity families.

With eight pairs, the minimum attainable two-sided exact p value is 2/256=0.0078125, which exceeds the first Holm thresholds for families of 12 or 15 tests. Nonrejection in these larger families is therefore interpreted cautiously, with emphasis on effect estimates and intervals. The independent-evaluation primary and secondary families were reported separately.

#### Software and Computational Resources

The model was implemented in Python using NetPyNE and NEURON [43, 44] and run on four Linux high-performance multicore servers (Table 4), each equipped with two Intel Xeon Gold or Platinum processors, 48-64 physical cores, and 503 GB-1 TB RAM. Across the 32,000 completed long-duration training episodes, each simulating 1,500 s with 30 Message Passing Interface (MPI) ranks, the median measured wall time was 357.15 s (5.95 min; interquartile range, 331.81-403.62 s). These simulations therefore comprised approximately 3,416 hours (142.3 host-days) of cumulative within-job wall time. Based on the mean observed job duration for each host, the four-host configuration corresponds to an arithmetic continuously busy capacity of approximately 37.6 episodes/hour, equivalent to about 851 hours (35.5 days) under ideal continuous four-host utilization. Actual campaign duration can be longer because the hosts were shared and execution included pauses, retries, queue delays, and periods when not all four hosts were simultaneously occupied. Supplementary Methods **S9** provides additional computational-resource and runtime details. The source code locations are https://github.com/NathanKlineInstitute/NavNetEGP and ModelDB entry https://modeldb.science/2041698 [48].

**Table 4.** Experiment inventory. Network pairs, episodes and horizons describe the completed protocols; single-network STDP/RL examples are descriptive.

| Experiment | Networks / comparisons | Episodes or trials | Target horizon |
|---|---|---|---|
| Single-target STDP/RL | 1 network | 800 learning + 400 post-learning | 187.5 s |
| Multi-target STDP/RL | 1 network; 5 targets | 4000 learning + 500 post-learning | 187.5 s |
| Short EGP | Initial 10 + prospective 30 pairs | 800/arm/seed | 7.5 s |
| Long EGP | 8 global/context pairs | 2000/arm/seed | 30 s |
| Independent evaluation | 8 pairs; 9 checkpoints | 7200 fresh-start trials | 187.5 s |
| Spatial maps | 1 fixed pretraining network | 2 coordinate sweeps; 1536 cells | 2.5 s/bin |
| EGP components | 8 seeds; A/C/D/E + B shadow | 200/condition/seed; 6400 total | 7.5 s |
| Gain controls and STDP/RL benchmark | 8 seeds; A/C/E/G/N/S/R/Q | 200/condition/seed; 12,800 comparative episodes | 7.5 s |
| Matched frozen-weight evaluation | 64 final networks; 8 conditions | 50 trials/network; 3200 total | 187.5 s |

## Results

### Spatial and target-related populations provide structured coordinate representations

Before analyzing learned navigation, we characterized the spatial and goal-related signals available to the circuit. Spike-derived maps show agent-position coding in EGrid/EPlace and target-coordinate coding in EV/EVPlace (**Fig. 2**; Supplementary **Figs. S3-S8**). Across EGrid, EPlace, EV and EVPlace, median raw information was 3.09, 4.66, 3.09 and 4.68 bits/spike. The higher information per spike in EPlace and EVPlace is consistent with their more localized firing patterns. Individual maps often contained multiple fields; irregular EPlace fields are compatible with experimental observations of multiple hippocampal place fields in larger environments [7, 81].

**Figure 2.**
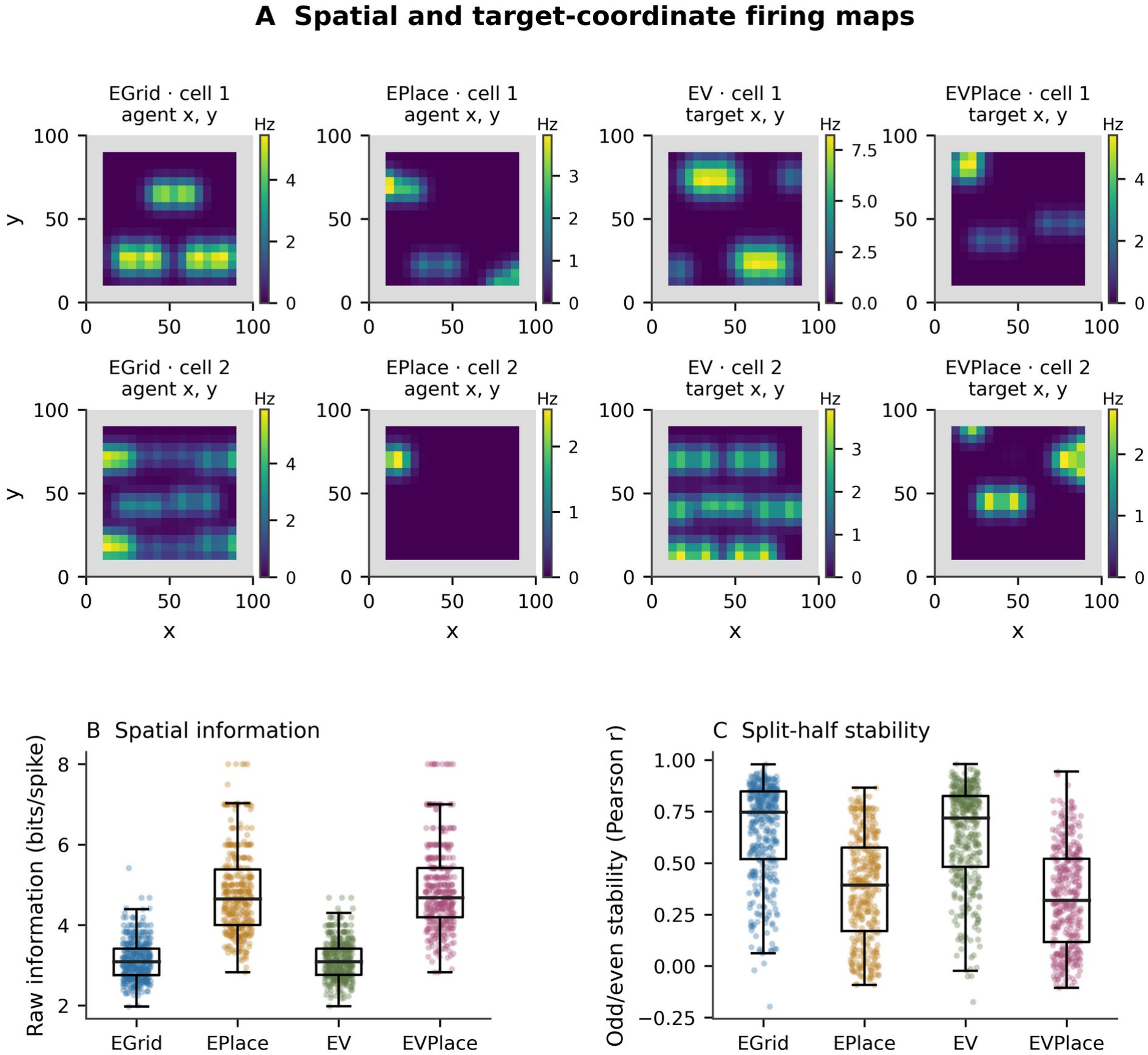
Spatial and target-coordinate firing maps. **A)** The first two of 12 preselected cells per population in one fixed pretraining network. EGrid/EPlace use agent-position coordinates; EV/EVPlace use target coordinates. Each *viridis* color map has its own Hz scale; gray denotes unobserved padding. Display maps use one-bin smoothing. **B)** Raw occupancy-normalized information per spike. **C)** Secondary split-half reliability: Pearson correlation between smoothed maps from alternating sets of 25 visits/bin. Dots show every defined cell; boxes show the median, interquartile range (IQR), and 1.5-IQR whiskers. Information is defined for 384, 368, 384 and 376 cells, and correlations for 384, 362, 384 and 363 cells, respectively. All 384 cells/population, including silent or undefined cases, remain in the inventory. All 48 examples and occupancy coverage are in the Supplement. Neurons are not independent network replicates.

Split-half correlations were lower downstream: median values were 0.75, 0.39, 0.72 and 0.32 for EGrid, EPlace, EV and EVPlace. The contrast between these results shows that information per spike and split-half reproducibility capture different properties of the populations. EPlace contained 16 silent cells and EVPlace eight, whereas EGrid and EV had none. Median threshold-defined field counts were four, three, four and three, respectively. Full distributions and examples, including less structured maps, are retained in the Supplement.

### A spiking navigation circuit learns a single target with STDP/RL

We next tested whether these coordinate representations could support learned navigation. The model learned a single-target navigation policy using STDP-based reinforcement learning (STDP/RL). In this single-network demonstration, the agent navigated in a two-dimensional environment containing a single target, while plasticity modified synaptic projections from direction-specific association populations to their motor-output populations. Across training episodes, the number of target hits increased substantially, indicating that the network learned to transform its spatial and target-related activity into actions that brought the agent toward the goal (**Fig. 3A**).

To determine whether learning reflected durable changes in synaptic weights rather than only transient online adaptation, we next evaluated performance after learning was disabled. The agent maintained similar target-reaching performance when plasticity was turned off, indicating that the learned policy was stored in the synaptic configuration of the network rather than requiring ongoing weight updates during each episode (**Fig. 3B**). Consistent with this interpretation, plastic synaptic weights increased over training (**Fig. S1**). The late-learning example (episode 799) showed trajectories directed from the starting location toward the target, with repeated target interceptions visible in the distance-to-target trace (**Fig. 3C,D**). Raster plots showed activity in spatial representation populations (EGrid, EPlace), target-related populations (EVGrid, EVPlace), association neurons (EA), and motor-output populations (EM) during navigation (**Fig. 3E**).

Together, these results show that STDP/RL can establish a single-target navigation policy in this network that persists without ongoing plasticity.

**Figure 3.**
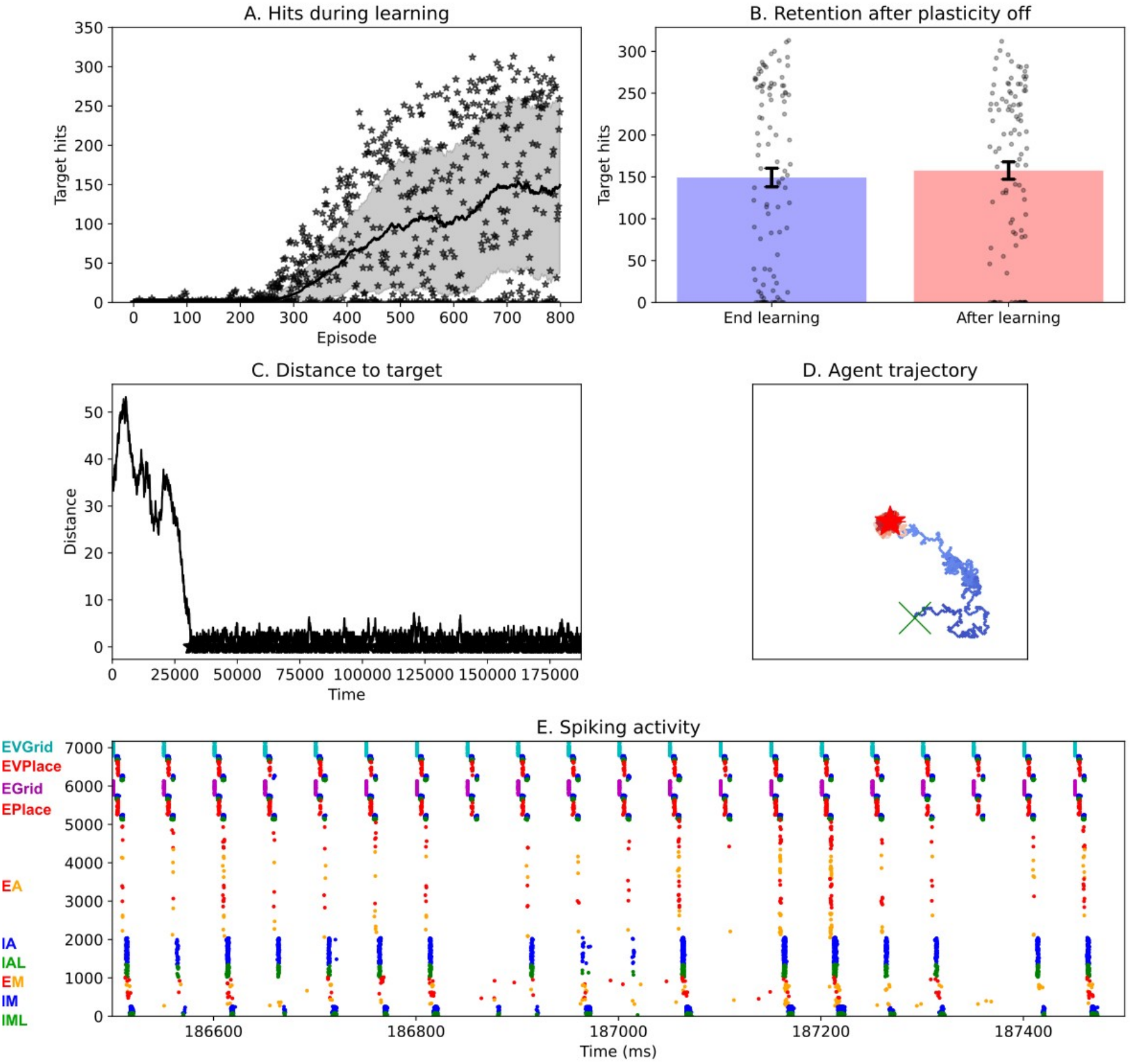
Single-target learning and retention with STDP-based reinforcement learning (STDP/RL) in one network. **A)** Target hits across 800 learning episodes. Dots are episodes; the line and shading show a 100-episode moving mean ± standard deviation (SD). A hit is exact target contact, and repeated contacts can occur in one episode. **B)** Final 100 learning episodes (700-799) and first 100 post-learning episodes with plasticity disabled. Bars show episode means and error bars SEM; these are repeated episodes within one network, not independent network replicates. **C-E)** Distance to target, trajectory and spike raster from late-learning episode 799, while plasticity was still active. **C)** Distance is in pixels and time is in milliseconds. **D)** The green X is the start, the red star the target, and blue-to-red color denotes progression through the episode. **E)** Each point is a spike; time is in milliseconds and the vertical ordering identifies model populations. The raster illustrates population activity during navigation.

### Multi-target learning with STDP/RL leaves uneven target selectivity

Online STDP/RL also produced target-directed behavior across multiple goals, although selectivity remained uneven. In a second single-network demonstration, the agent was trained on five targets distributed across the environment. Early in training, trajectories were broadly exploratory and did not show consistent attraction to the active target. Blate learning, trajectories became more directed toward the current target, indicating that the network acquired target-dependent navigation policies. However, the learned behavior remained uneven across targets and performance declined when plasticity was turned off (**Fig. 4**).

Quantitative analysis of target hits confirmed this pattern (**Fig. 5**). During learning, the number of target hits increased for all five targets, but performance differed substantially across target locations. Disabling plasticity reduced target-reaching performance for several goals, indicating incomplete retention of some target-directed behaviors.

STDP/RL therefore acquired multiple target-directed behaviors, while uneven retention highlighted the challenge of maintaining several policies within shared synaptic projections.

To further characterize this selectivity, we measured the fraction of each episode that the agent spent near each target location (**Fig. 6**). Under ideal target-specific behavior, dwell time should be concentrated along the diagonal of the matrix, corresponding to time spent near the active target. Instead, multi-target STDP/RL produced off-diagonal dwell, especially after learning was turned off, indicating attraction to incorrect target locations. These results illustrate useful target-directed behavior alongside off-target occupancy in one network. The following experiments evaluate the two EGP variants in paired networks under matched training conditions.

**Figure 4.**
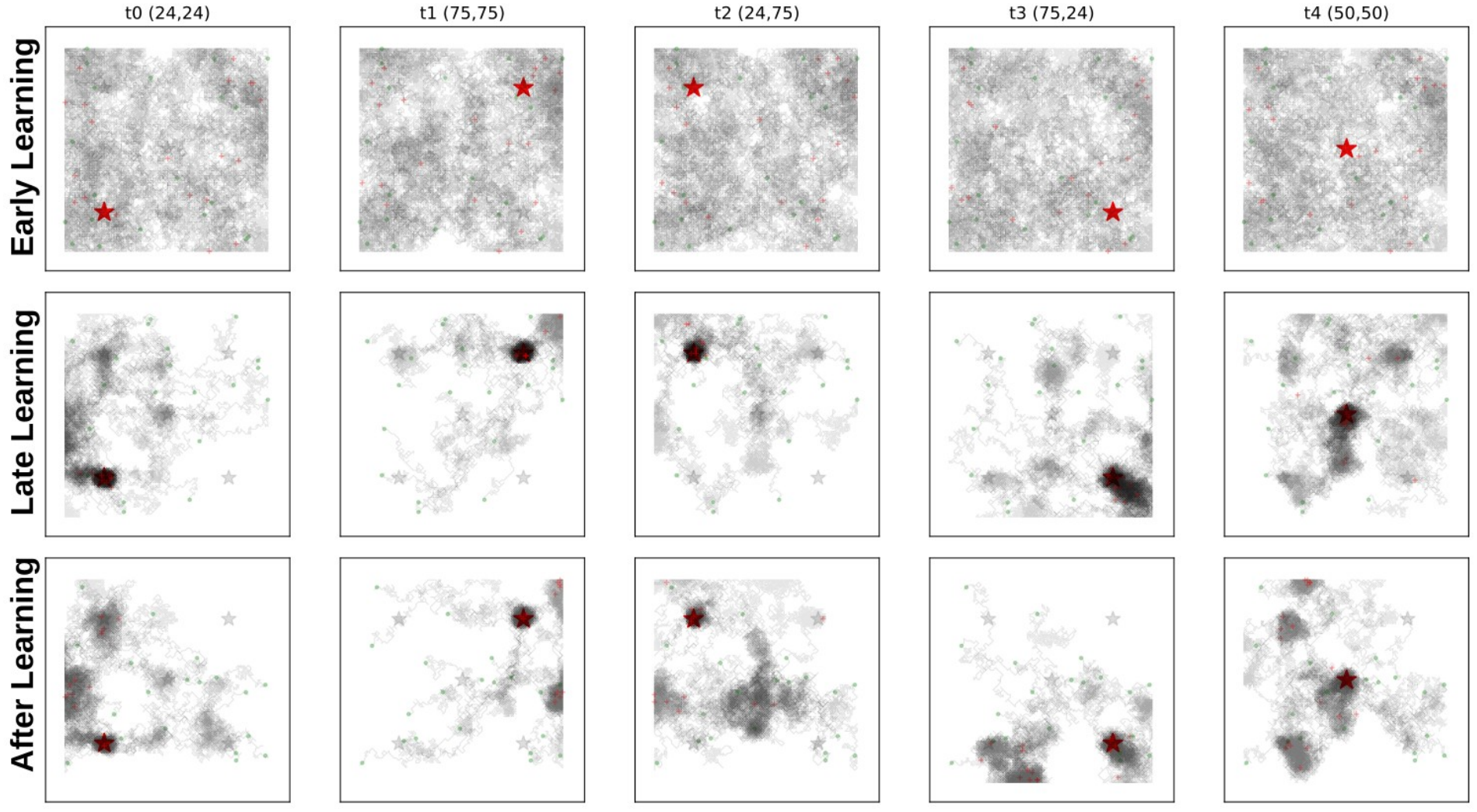
Multi-target STDP/RL trajectories in one network. Columns show the five active goals, and rows show early learning (episodes 0-99), late learning (episodes 3900-3999), and post-learning evaluation (episodes 400-499). Each displayed stage contains 20 episodes per target. Red stars mark the active target, gray stars indicate inactive targets, green circles show starting locations, and red X symbols mark final positions; coordinates are in pixels. Trajectories become more strongly directed toward the active goal during late learning, although off-target occupancy remains visible. Online plasticity is disabled during the post-learning evaluation. These trajectory examples complement the quantitative target-hit and dwell analyses in **Figs. 5-6**.

**Figure 5.**
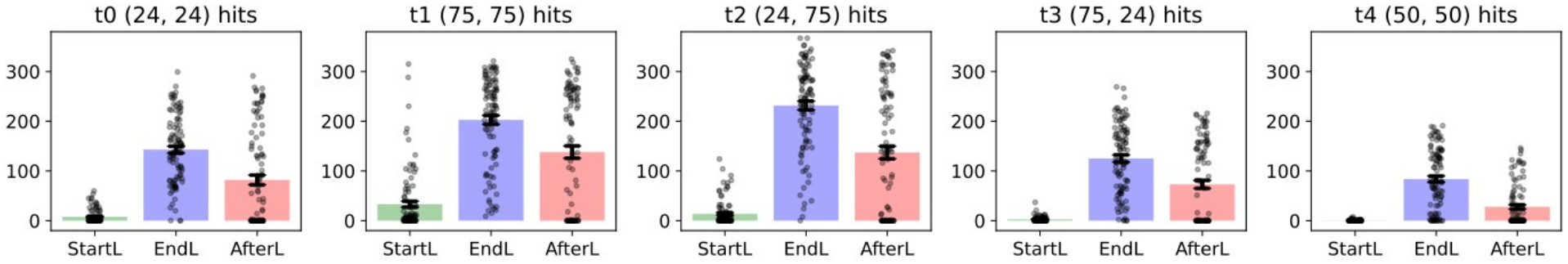
Target-dependent STDP/RL performance. Each target has 100 observations per stage: the first 100 learning cycles (“StartL”), learning cycles 700-799 (“EndL”), and post-learning cycles 0-99 (“AfterL”). A per-target cycle is distinct from the combined episode windows used in Figs. 4 and 6. Points show exact target-contact counts during 187.5-s episodes; repeated contacts can occur within an episode. Bars show means and error bars indicate SEM across episodes within this network. Target contacts increased during learning and decreased after plasticity was disabled, with substantial differences across target locations.

**Figure 6.**
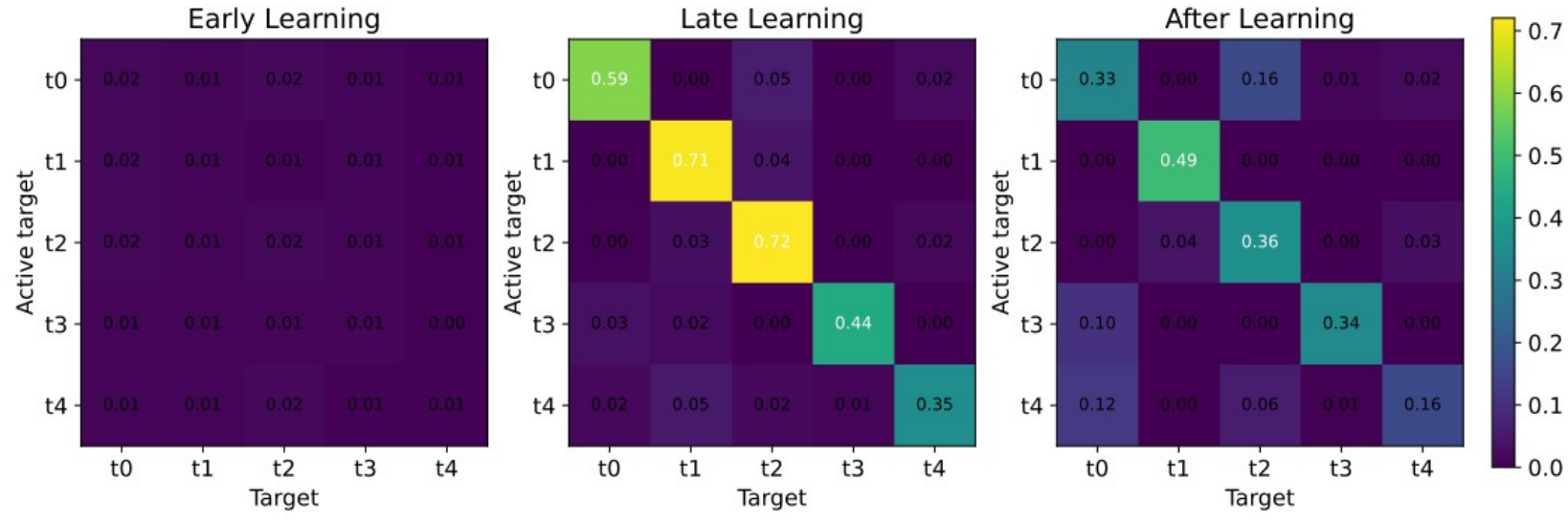
Multi-target STDP/RL dwell matrices in one network. Rows indicate the active target and columns the target region occupied. Entries show the fraction of observed time spent within 5 pixels of each target location, using a shared scale from 0 to 0.72044. Diagonal entries therefore represent dwell near the active target, whereas off-diagonal entries represent dwell near inactive targets. Early and late learning use combined episodes 0-99 and 3900-3999, respectively, and post-learning uses episodes 400-499, with 20 observations per target in each stage. Online plasticity is disabled during the post-learning stage. Diagonal structure increases during learning, while residual off-diagonal occupancy indicates incomplete target selectivity. The corresponding controlled global-versus-target-context EGP dwell analysis is shown in Fig. 10.

### Global EGP separates rapid proposals from durable consolidation

The STDP/RL demonstrations motivated a strategy for managing shared synaptic updates. Global EGP separates rapid reward-modulated proposals from slower durable commitment (**Fig. 1**): proposals accumulate during TRAIN, their aggregate is expressed during TEST, and need-weighted evidence modulates consolidation. Target-context EGP extends this foundational learning rule with goal-specific consolidation.

Global EGP acquired substantial navigation competence that persisted when learning was disabled. Independent fixed-weight success rose from the shared 17.8% pretraining baseline to 56.0% for global EGP and 67.8% for target-context EGP. During long-training internal TEST, target-context EGP also increased contact probability and reduced expected time to reach the goal. The paired comparisons below quantify these benefits and their variability.

### Target-context EGP improves early reward across paired networks

Across 40 paired network realizations, target-context EGP showed a consistent early reward advantage. Mean TEST raw reward per target block over episodes 0-99 was 0.0035 for global and 0.0096 for target-context EGP, a difference of 0.0061 (95% CI 0.0031-0.0093; exact p=0.000190), with 34 of 40 pairs favoring target-context EGP. The pooled analysis combines the initial 10 pairs with the independently specified 30-pair replication (**Supplementary Table S1**).

The effect was independently confirmed in the additional 30-pair replication, whose analysis was specified before those simulations (**Fig. 7**). Early reward was 0.0040 for global and 0.0090 for target-context EGP, a difference of 0.0050 (95% CI 0.0015-0.0088; exact p=0.010), with 24 of 30 pairs favoring target-context EGP. Over the final 100 episodes of the replication, the reward difference widened to 0.0252. These results support the complete target-context rule; the component experiment below separates its operations. Complete cohort-specific statistics and short-run dwell summaries are retained in the Supplement.

**Figure 7.**
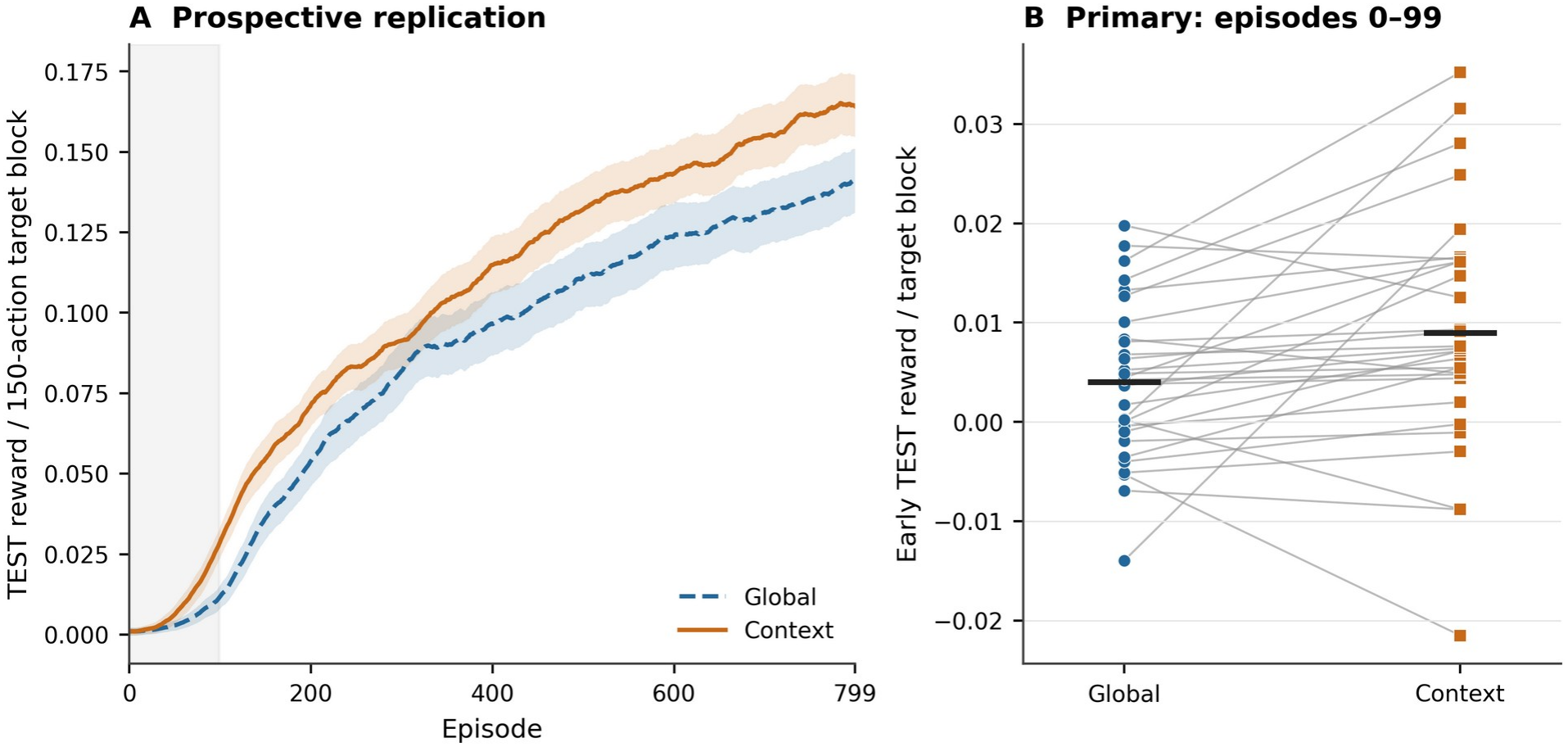
Prospective short-run comparison of global and target-context EGP. **A)** Mean TEST raw reward per 150-action target block across 800 episodes for 30 paired networks. Curves use a centered 25-episode display window; shading is mean ± between-seed SEM, with the mean and SEM smoothed separately. The shaded early window is episodes 0-99. **B)** The prespecified early endpoint for every paired seed; connecting lines identify pairs and black marks show arm means. Target-context EGP had higher early reward in 24 of 30 pairs (exact two-sided paired sign-flip test, p=0.010). The 95% paired-bootstrap interval for the context-minus-global difference was 0.0015-0.0088. Internal TEST temporarily expresses candidate weights; durable fixed-weight performance is evaluated separately in Fig. 11.

### Target-context EGP improves long-duration reward across targets with late-stage variability

**Figure 8.**
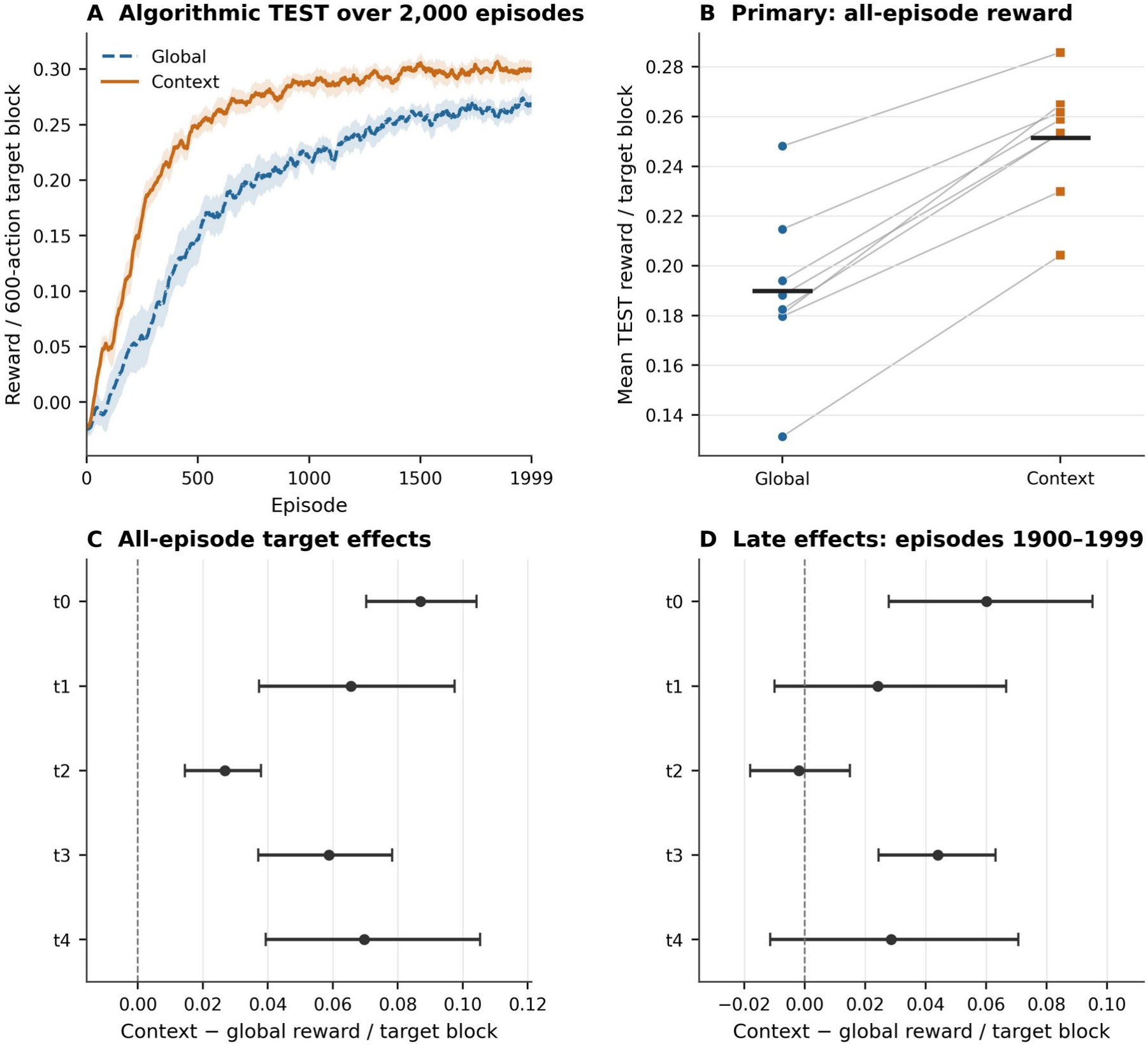
Long-duration reward across eight paired networks. **A)** Internal TEST reward over all 2000 episodes, shown as the mean ± between-seed SEM after a centered 25-episode moving average within each seed. **B)** All paired seed values for the primary mean TEST reward per 600-action target block; black marks denote arm means. Target-context EGP exceeded global EGP in all eight pairs (0.2515 versus 0.1899; paired difference 0.0616; exact two-sided paired sign-flip test, p=0.0078). **C,D)** Context-minus-global effects for every target across all episodes and during episodes 1900-1999. Error bars are unadjusted 95% paired-bootstrap confidence intervals; two-sided exact paired sign-flip p values were Holm-Bonferroni-adjusted within separate five-target families. All five all-episode effects were significant (adjusted p=0.0391 each). Late adjusted p values for t0-t4 were 0.0625, 0.773, 0.828, 0.0391 and 0.773, respectively; only t3 remained significant. Late effects varied across targets, including a small negative t2 estimate. These panels summarize performance during EGP learning; fresh-start fixed-weight performance is evaluated separately in Fig. 11.

The target-context reward advantage persisted over extended learning. Following the short-run replication, we compared both variants across 2000 episodes in eight paired networks (**Fig. 8**). The all-episode primary mean TEST reward was 0.1899 for global and 0.2515 for target-context EGP, a paired difference of 0.0616 (95% CI 0.0515-0.0713; exact p=0.0078), with all eight pairs favoring context. All five target-specific all-episode effects passed Holm-Bonferroni correction within their five-test family (adjusted p=0.0391). The reward advantage remained positive over the final 100 episodes (difference 0.0310; 0.2674 versus 0.2984), although its magnitude was smaller than the all-episode difference. Late target effects were heterogeneous: target t2 had a small negative difference (-0.0019), and only t3 passed correction within the separate five-test late family. These results demonstrate an acquisition-wide reward advantage, with late-stage effects that vary across targets.

The internal dynamics distinguish proposal generation, evidence modulation and the changes retained in durable weights (Supplementary **Fig. S11**). Raw reward differences, transformed evidence, consolidation coefficients, proposal magnitude and cancellation describe distinct stages of this process. These trajectories describe the successive computational stages of learning; the component experiment below examines the reward contribution of specific operations.

### Target-context EGP improves navigation and conditional target selectivity

Target-context EGP increased late-learning goal contact and conditional target selectivity. **Figure 9** shows the preselected seed-1 network in episode 1999, retaining every target and all five pieces in each TRAIN/TEST phase for both algorithms. That network’s final-100 success was 77.2% for global and 85.4% for target-context EGP, above the corresponding ensemble means. These values summarize 100 episodes, whereas the trajectories show a single episode; the example illustrates a relatively high-performing realization rather than typical performance across networks.

**Figure 9.**
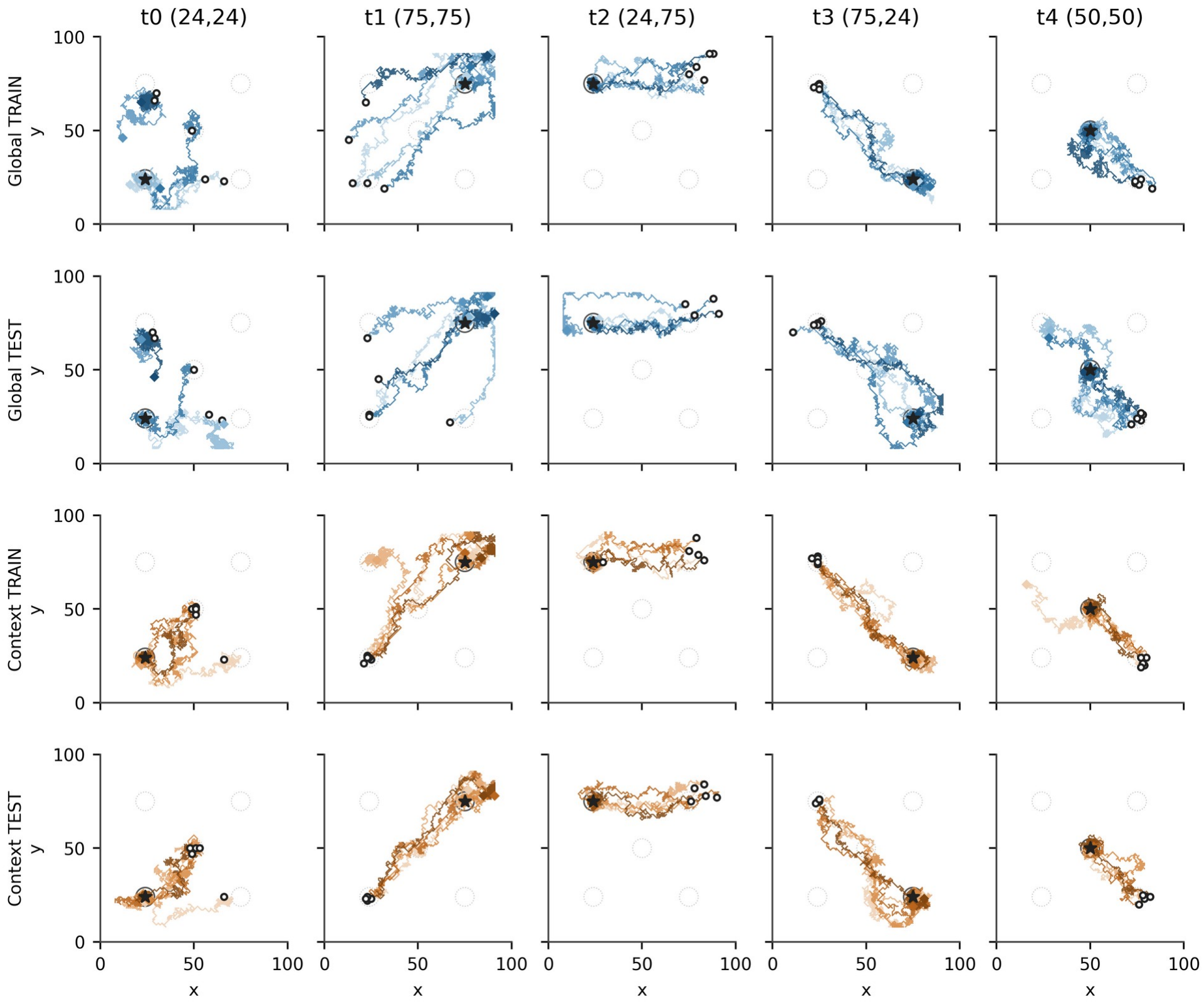
Fixed late-learning EGP trajectories. Seed 1, episode 1999, was selected in advance for visualization. Columns show the five target locations, and rows show global TRAIN, global TEST, target-context TRAIN, and target-context TEST. Each panel contains the five consecutive 600-action trajectory segments for the corresponding target and phase, with darker shades indicating successive segments. Open circles indicate the first recorded post-action positions, diamonds indicate final positions, and black stars mark the active targets. Open circles therefore show recorded trajectory positions rather than reconstructed reset locations. Circles around the goals indicate the 5-pixel reference radius and are not barriers. All 60,000 recorded trajectory points are shown on identical 0-100 pixel axes.

Across eight networks, final-100 exact-target success in 30-s internal-TEST blocks averaged 57.3% for global and 66.6% for target-context EGP, with individual-network values ranging from 27.6-81.0% and 37.0-85.4%, respectively (Supplementary Table S15). Target-context EGP increased the probability of reaching the active goal by 9.3 percentage points (95% CI 5.2-13.5) and reduced restricted mean time to contact from 21.12 to 19.57 s (difference -1.54 s, 95% CI -2.20 to -0.92); both adjusted p values were 0.0313 in the four-endpoint family. Among blocks that reached the target, contact time and path inefficiency did not consistently improve (both adjusted p=1). The main navigation benefit was thus reaching the target in more blocks, rather than uniformly shortening already-successful trajectories. Inference was at the paired-network level.

Target-context EGP improved selectivity primarily by increasing occupancy near the correct target (**Fig. 10**). Correct-target dwell rose from 27.3% to 32.7%, a 5.4-percentage-point increase (approximately 19.8% relative; 95% CI for the absolute difference 3.0-7.9 points; nominal exact p=0.0078). Because each 30-s block includes travel and approach time, the dwell fractions quantify occupancy near the goal rather than the proportion of successful blocks; 100% dwell is therefore not an expected benchmark.

Absolute time near incorrect targets changed little: raw wrong-target dwell was 6.3% for global and 6.5% for target-context EGP (nominal p=0.727). Because correct-target occupancy increased, incorrect targets accounted for a smaller share of all time spent near any goal: conditional wrong-target dwell fell from 20.6% to 18.3% (nominal p=0.0078). Thus, target-context EGP improved selectivity by increasing occupancy near the intended goal, while absolute occupancy near competing goals remained similar.

Together, contact, time and dwell measures show that target-context EGP improves goal acquisition and selectivity across paired networks, while performance remains heterogeneous across targets and networks. Internal TEST temporarily expresses candidate weights; the independent evaluation below tests competence stored in the durable weights with learning and proposal mechanisms disabled.

**Figure 10.**
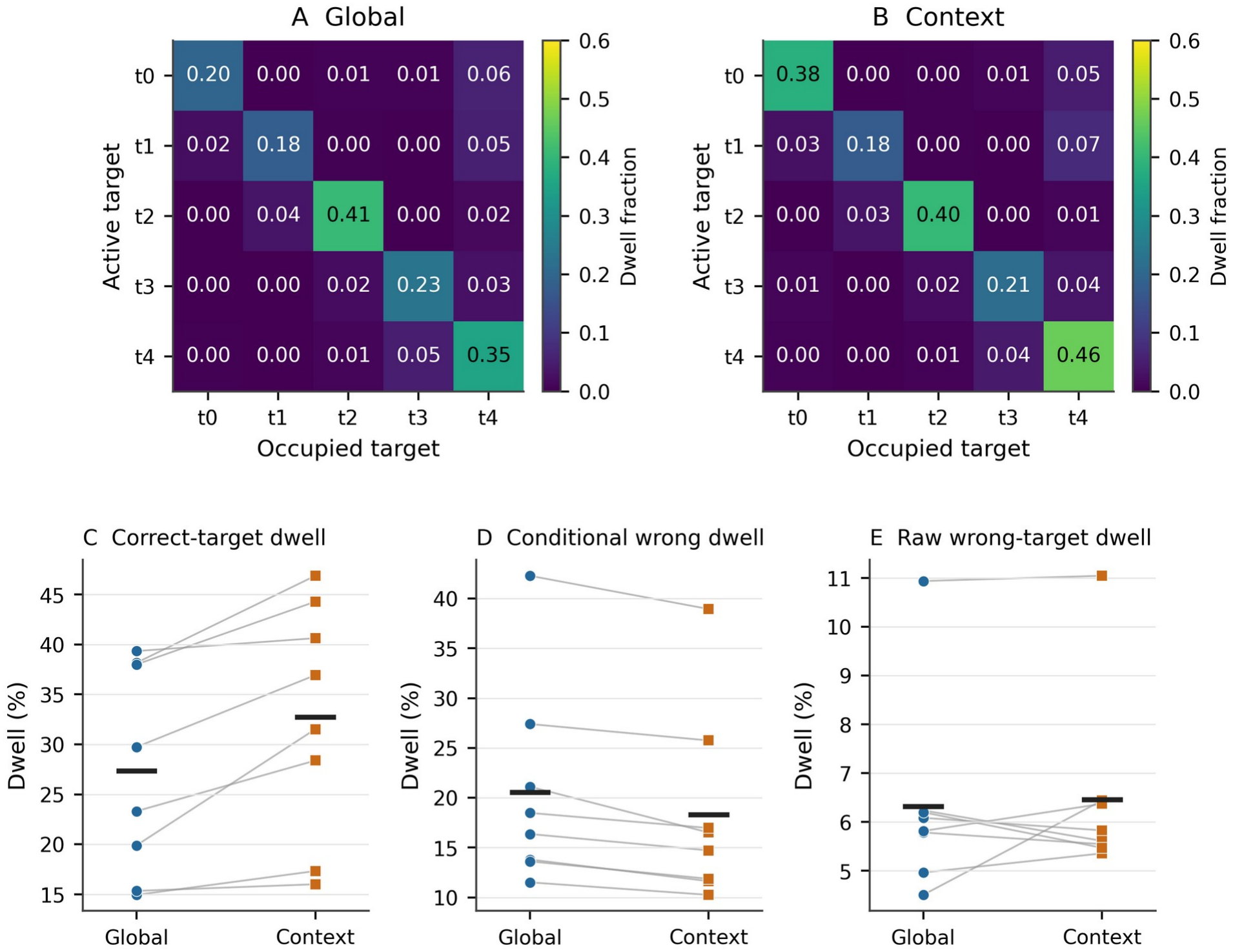
Controlled late-learning target selectivity. **A,B)** Global and target-context EGP dwell matrices, averaged over eight seeds during TEST episodes 1900-1999. Rows denote active goals; columns denote occupied 5-pixel goal regions. Both matrices use the same viridis color scale (0-0.6, fraction of observed time). **C-E)** Every paired seed’s correct-target dwell, conditional wrong-target dwell and raw wrong-target dwell; black marks denote arm means. Correct and raw wrong dwell use all observed time as denominator, while conditional wrong dwell divides wrong dwell by total dwell near any goal. Correct dwell increased and conditional wrong dwell decreased, but raw wrong dwell did not consistently decrease.

### Learned durable weights support navigation after plasticity is disabled

Both algorithms acquired substantial navigational competence that persisted with learning and proposal mechanisms disabled (**Fig. 11**). Final independent success averaged 56.0% for global and 67.8% for target-context EGP, compared with 17.8% before training in both arms. Individual-network values ranged from 18.0-80.0% and 56.0-86.0%, respectively (**Supplementary Table S15**). Seven of eight pairs favored target-context EGP; the estimated advantage was 11.8 percentage points (95% CI 2.3-22.2). The prespecified final exact test did not reach α=0.05 (p=0.063), so the between-algorithm advantage remains directional. The percentile bootstrap interval estimates uncertainty by resampling the observed pairs, whereas the prespecified exact test enumerates sign assignments under its null; these procedures are not inferentially equivalent. The exact test governs the confirmatory conclusion despite the interval excluding zero. Pretraining-to-final gains are descriptive, and pretraining performance is an empirical baseline rather than chance.

Checkpoint curves showed an earlier, larger descriptive target-context advantage and durable gains in both algorithms. Final restricted mean first-passage time was 104.32 s for global and 84.57 s for target-context EGP (adjusted p=0.0938); correct-target dwell was 24.8% and 32.6% (adjusted p=0.0547). These secondary families did not reach their Holm-adjusted thresholds. The overall pattern demonstrates substantial durable learning in both rules, while the final between-rule advantage remains suggestive rather than statistically resolved by the specified tests.

**Figure 11.**
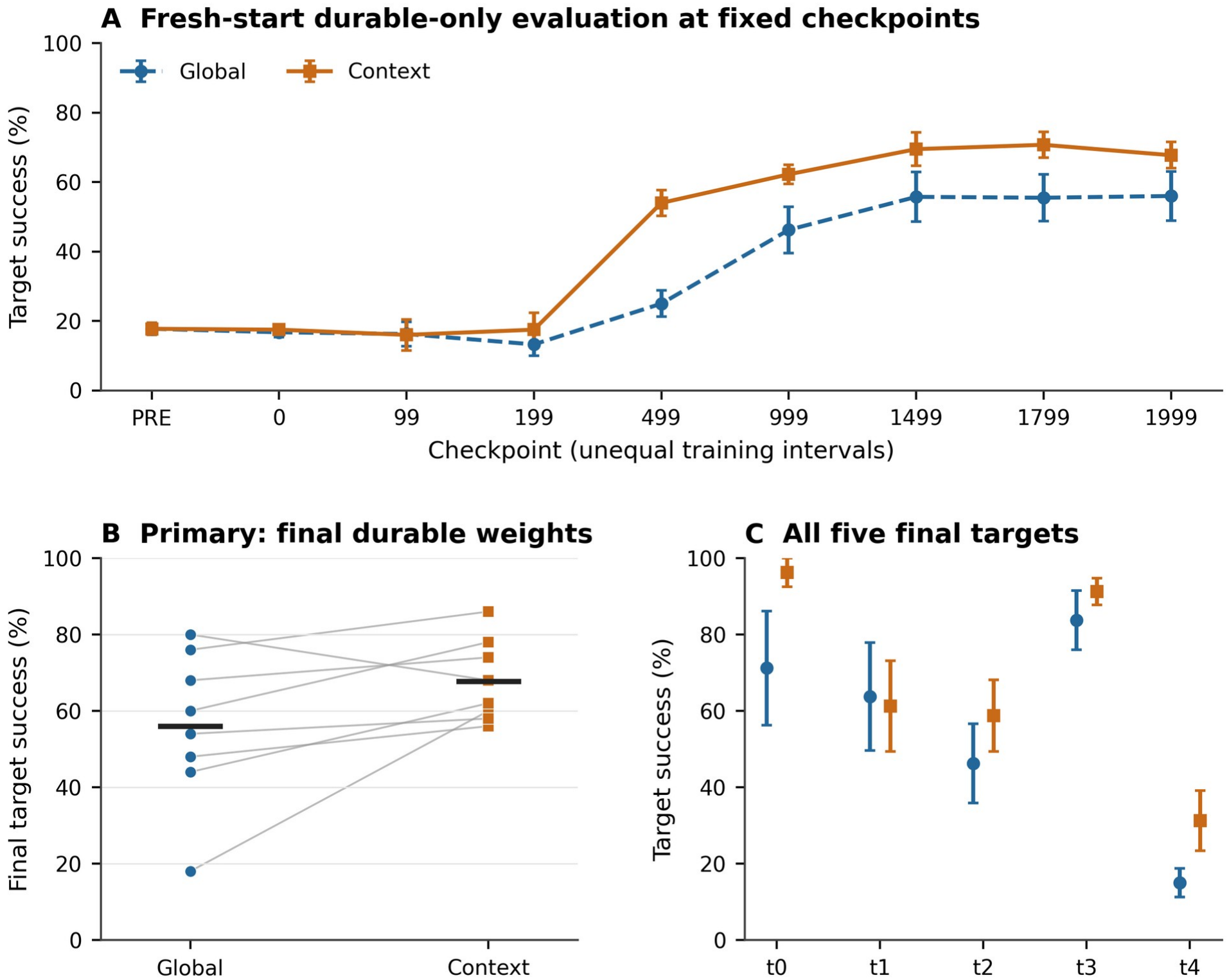
Independent frozen-weight evaluation. All learning and proposal mechanisms were disabled during fresh-start trials. **A)** Pretraining and all eight post-training checkpoints; checkpoint spacing is categorical, not proportional to training time. **B)** All eight final paired success values, with arm means marked in black. Both algorithms retained substantial navigation competence: final success was 56.0% for global and 67.8% for target-context EGP, a difference of 11.8 percentage points, with seven of eight pairs favoring target-context EGP. The prespecified two-sided exact paired sign-flip test did not reach α=0.05 (p=0.063). **C)** All five final targets; none reached significance after Holm-Bonferroni correction within the five-target family. Checkpoint and target error bars are between-seed SEM, not confidence intervals. The experiment contains 7200 trials, with 50 trials per arm/seed/checkpoint and a 187.5-s horizon. It is separate from EGP’s internal TEST phase.

### Need-dependent consolidation scaling contributes to the target-context reward advantage

Full target-context EGP again exceeded global EGP in early reward (**Fig. 12**). Across eight paired networks, mean TEST reward over episodes 0-99 was 0.0023 for global and 0.0099 for full target-context EGP, a difference of 0.0076 (95% CI 0.0043-0.0114; exact p=0.0078), with all eight pairs favoring the full rule. Learning curves show reward acquisition in all four behavioral conditions; the segmented global shadow is identical to global EGP behavior and is not an additional policy.

**Figure 12.**
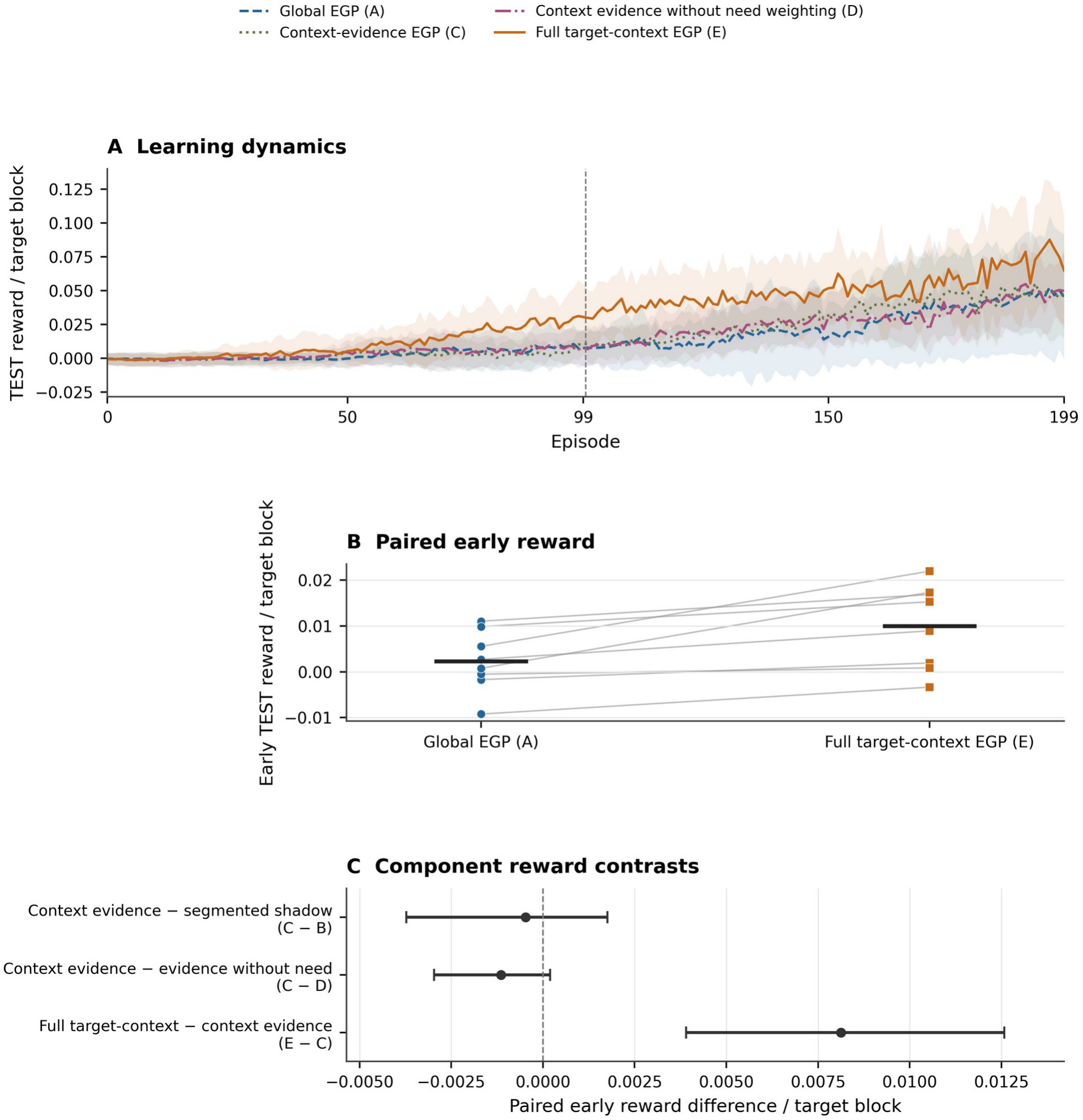
Reward acquisition and components of context-specific consolidation. **A)** All 200 episodes for global EGP (A), context-evidence EGP (C), context evidence without need weighting (D), and full target-context EGP (E). Curves and bands are unsmoothed mean ± between-seed SD over eight networks, not confidence intervals; the divider separates episodes 0-99 from 100-199. The segmented global shadow (B) has A’s behavior and is not a fifth stream. **B)** All eight paired early reward values for full target-context and global EGP, with arm means in black; the paired difference was 0.0076 (95% CI 0.0043-0.0114; exact p=0.0078). **C)** All three component reward effects with unadjusted 95% paired-bootstrap intervals. Holm-adjusted p values are 0.930 for context evidence minus segmented shadow (C−B), 0.781 for context evidence minus evidence without need (C−D), and 0.0234 for full target-context minus context-evidence EGP (E−C). Only the added need multiplier on the consolidation coefficient had adjusted reward support.

Full target-context EGP significantly outperformed the otherwise context-specific condition lacking the additional need multiplier on the consolidation coefficient: the early reward difference was 0.0081 (adjusted p=0.0234). This was the only component contrast with adjusted reward support. Context-evidence EGP versus the segmented global shadow differed by -0.0005 (adjusted p=0.930), and context evidence with versus without need weighting by -0.0011 (adjusted p=0.781). The separate contributions of context-specific evidence and need weighting therefore remain unresolved in these comparisons.

Full target-context EGP also showed descriptive increases in durable-change magnitude, cancellation and coefficient dispersion relative to context-evidence EGP. None of the 12 mechanistic tests or the separate five-test and 15-test behavioral families reached adjusted thresholds, so these secondary measures do not isolate a single intermediate process that accounts for the reward improvement. In particular, the reward benefit was not accompanied by reduced cancellation. Complete estimates and stage-specific clipping remain in the Supplement.

Because the 7.5-s component protocol was designed around shaped-reward acquisition, exact contact was a stringent secondary outcome and did not show an adjusted advantage within that horizon. The early contact proportions were approximately 1-2%, so the supported E-C effect concerns dense-reward acquisition rather than improved successful navigation. Longer internal TEST and independent evaluation provide complementary behavioral evidence, but do not isolate which consolidation component produces it. Target-specific component dynamics remain descriptive (Supplementary **Fig. S10**).

### Gain controls clarify the contribution of consolidation strength

To determine whether the reward advantage of target-context EGP reflects the allocation of consolidation across goals or an overall increase in consolidation strength, we compared the complete algorithm with three additional controls. A uniform gain-matched rule produced similar reward performance, with closely matched mean consolidation coefficients and retained synaptic changes. The difference from full target-context EGP did not reach significance after correction (adjusted p=0.141). In contrast, normalizing the need-dependent coefficients to remove their increase in average consolidation strength reduced reward performance (adjusted p=0.023). Shuffling the correspondence between target contexts and their reward evidence did not produce a statistically supported difference from the full rule. These results suggest that effective consolidation gain contributes importantly to reward acquisition, while the additional contribution of context-specific allocation remains unresolved (**Supplementary Figs.** S12–S14; Table S17).

### Immediate plasticity and evidence-gated consolidation differ in reward learning and retained navigation

The preceding experiments demonstrated that EGP networks can acquire multiple target-directed navigation policies and retain substantial behavioral competence after plasticity is disabled. We next investigated how this evidence-gated approach compares with conventional STDP/RL, in which eligible synaptic changes are committed immediately. Using matched networks and equal environmental interaction budgets, we evaluated reward acquisition during training and subsequent navigation using the learned, fixed synaptic weights.

Conventional STDP/RL acquired shaped reward substantially faster than either EGP variant, with the largest improvements occurring early in training (**Fig. 13A**). EGP learned more gradually and continued improving throughout the 200-episode protocol. However, the relative performance changed when synaptic plasticity was disabled. In independent fresh-start evaluations, both EGP variants achieved higher exact-target success than the continuously learning STDP/RL baseline (**Fig. 13B**). Target-context EGP exceeded conventional STDP/RL in all eight paired networks. Although absolute success rates remained modest after this shorter training protocol, the findings demonstrate that faster reward acquisition during online learning does not necessarily correspond to stronger retained target-directed behavior.

Immediate STDP/RL also produced larger durable synaptic changes than EGP, consistent with their different learning mechanisms: direct reinforcement-driven updates versus evidence-dependent consolidation of transient proposals (**Supplementary Fig. S14**). The experiments matched environmental experience rather than total synaptic-change magnitude.

**Figure 13.**
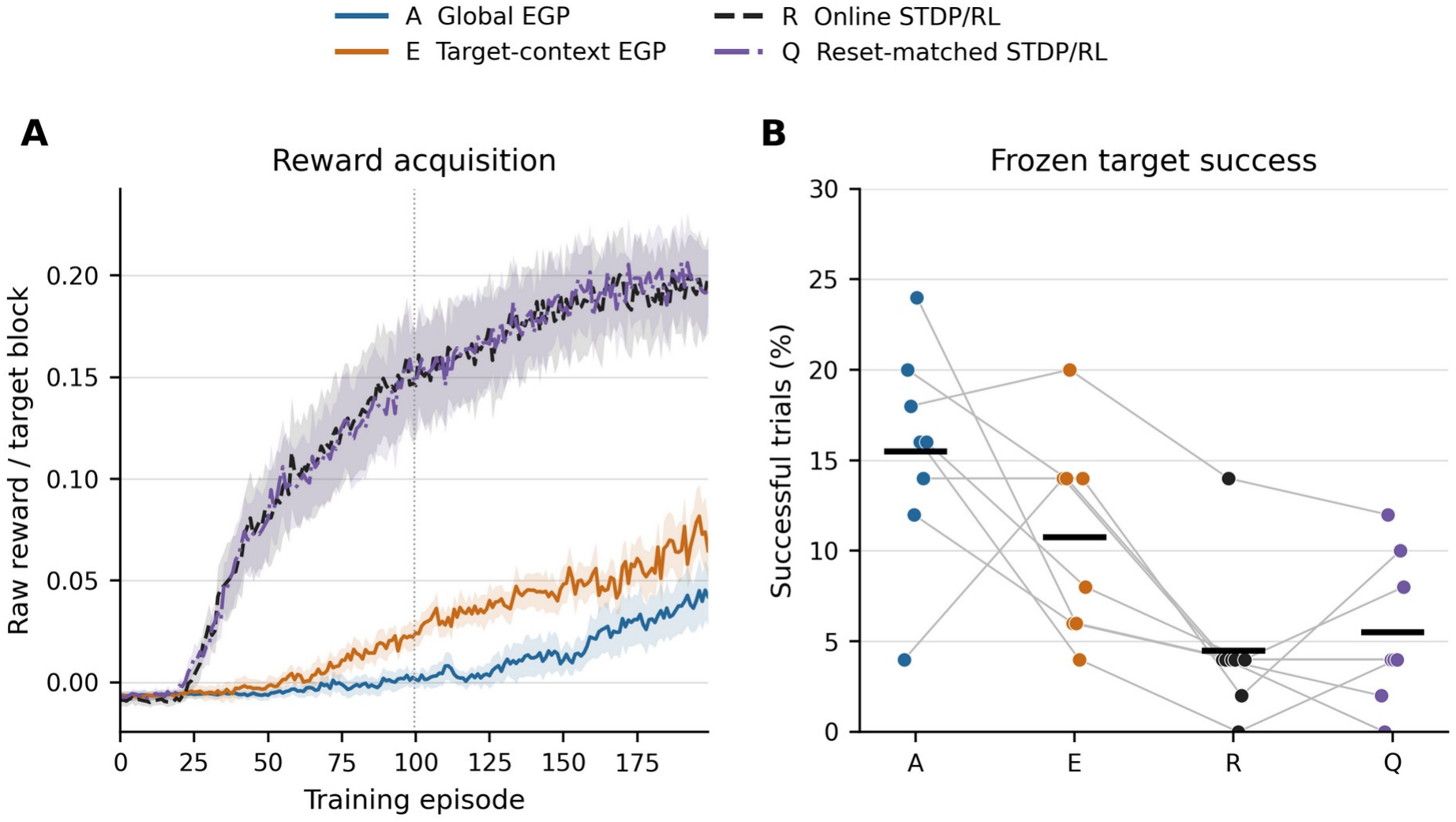
Reward acquisition and retained navigation under matched training experience. **A)** All-interaction raw reward per 7.5-s target block across 200 episodes for global EGP (A), target-context EGP (E), online STDP/RL (R), and reset-matched STDP/RL (Q). Lines are unsmoothed means and shading is between-seed SEM across eight paired networks. The divider marks the early comparison window (episodes 0–99); all four EGP-minus-STDP/RL early contrasts favored STDP/RL (Holm-adjusted p=0.03125; Table S17). Each network received 1.5 million interactions. **B)** Final durable weights evaluated with learning and proposals disabled in 50 matched fresh-start trials per network, each lasting 187.5 s. Points are seed success percentages, gray lines connect paired seeds, and black bars are means: 15.50%, 10.75%, 4.50%, and 5.50% for A, E, R, and Q. These frozen-weight comparisons are descriptive; no inferential p values were calculated. Exact-target contact measures retained navigation competence separately from shaped reward during plasticity. Complete seed-level outcomes are reported in Supplementary Table S18.

## Discussion

These experiments connect structured coordinate representations and local plasticity to durable multi-goal learning. Global EGP separates rapid synaptic proposals from evidence-modulated consolidation, and target-context EGP extends it to improve acquisition and conditional selectivity. Independent evaluation demonstrates retained competence in both algorithms; component ablation supports need-dependent scaling of context-specific consolidation, with the gain controls narrowing its mechanistic interpretation. The matched benchmark distinguishes rapid reward acquisition during immediate plasticity from retained target-reaching behavior after learning is disabled.

Spike-derived maps showed structured agent-position responses in EGrid/EPlace and target-coordinate responses in EV/EVPlace. Their recorded activity extends the description beyond the imposed input fields. EPlace and EVPlace carried higher information per spike but lower split-half reliability than their upstream populations, emphasizing complementary aspects of representation. Multiple irregular EPlace fields are consistent with observations in larger hippocampal recording environments [7, 81]. These findings characterize the implemented coordinate representations in one fixed pretraining network, rather than showing that biologically realistic spatial coding emerged through learning. The modest downstream split-half reproducibility limits claims about the stability of individual spatial patterns. EVPlace target-coordinate responses are not intended to reproduce a specific known hippocampal cell class, but their combination of spatial and goal-related information is broadly consistent with experimental evidence that hippocampal firing can be modulated by goal direction, goal location, and reward [3, 82].

In the STDP/RL demonstrations, the circuit learned and retained a single-target policy and also acquired target-directed behavior across five goals, although multi-target selectivity remained uneven and declined after plasticity was disabled (**Figs. 3-6**).

These observations motivated EGP as a strategy for stabilizing multi-goal learning within the same shared circuit. These STDP/RL demonstrations establish a foundation for the matched learning-rule comparison presented in **Fig. 13**.

Global EGP provides the foundational consolidation algorithm. A shared proposal store and need-weighted evidence supported extended reward learning and durable multi-goal navigation. In independent fixed-weight evaluation, average success increased from 17.8% before training to 56.0% after training, demonstrating that evidence-modulated consolidation can support substantial durable learning without context-specific proposal stores. The 200-episode comparison extends this question to immediate STDP/RL under equal environmental-interaction budgets, without equating that short protocol with the long-duration study.

Target-context EGP improved early reward across 40 paired networks, including the independently specified 30-pair replication, and all-episode reward in eight long-training pairs (**Figs. 7-8**). The controlled comparisons held architecture, initialization, feedback and goal schedules constant. In the longer experiment, target-context EGP reached goals more often and spent more time near the intended target (**Figs. 9-10**), without consistently shortening already-successful routes or reducing absolute wrong-target dwell. Substantial across-network variability complements these ensemble-average benefits. The component experiment helps distinguish how evidence organization and need-dependent coefficient scaling contribute to this advantage.

Independent evaluation distinguishes retained competence from behavior under temporarily expressed candidate weights. Both algorithms developed durable navigation, with final success averaging 56.0% for global and 67.8% for target-context EGP. The corresponding network ranges, 18.0-80.0% and 56.0-86.0%, demonstrate substantial across-network variability and include several high-performing realizations. Seven of eight pairs favored target-context EGP, while the final exact test left the between-rule advantage directional (p=0.063).

The component experiment sharpened the mechanistic interpretation of the target-context advantage. Full target-context EGP outperformed context-evidence EGP after adjustment across the three reward contrasts, identifying the added need multiplier on the context-specific consolidation coefficient as the component with direct reward support. The uniform-gain control achieved closely similar mean coefficients and net durable L1 changes, and E−G did not reach its adjusted threshold. E−N retained support, whereas E−S did not. Increased effective consolidation gain therefore remains a plausible explanation for a substantial portion of the reward advantage; these controls do not establish an isolated context-allocation benefit or equivalence between E and G. Because aggregate TEST mixes the proposals, the mechanism is more precisely described as context-conditioned, need-scaled consolidation than as demonstrated target-specific causal credit assignment.

The matched comparison also highlights an important distinction between rapid behavioral adaptation and the retention of learned navigation policies. Conventional STDP/RL produced faster improvements in shaped reward, whereas EGP networks more frequently reached their targets after synaptic plasticity was disabled. This difference is consistent with the distinction between immediate synaptic modification and the delayed, evidence-modulated consolidation implemented by EGP. Although the present experiments do not establish that the consolidation mechanism alone accounts for the behavioral difference, they demonstrate that reward acquisition and retained navigation competence can follow different trajectories. This distinction may be relevant to biological learning systems, in which the stabilization of useful experience-dependent synaptic changes is not necessarily identical to maximizing immediate reinforcement.

This organization is consistent with the view that biological learning operates over multiple timescales [30, 31]. Rapid STDP-like processes can modify synaptic efficacy during behavior, enabling adaptation to local contingencies, while slower mechanisms determine which changes are retained. Neuromodulatory gating, replay, synaptic tagging, and systems-level consolidation have all been proposed as contributors to memory stabilization [34, 49, 50, 51, 52, 53]. EGP abstracts this principle into a computational framework in which local activity generates candidate modifications and longer-timescale reward evidence modulates how strongly they are consolidated above the baseline contribution. The target-context component is similarly motivated by evidence that hippocampal, entorhinal, prefrontal, and striatal circuits encode contextual information that influences memory retrieval, action selection, and plasticity[4, 54, 55, 56, 57]. In the present model, target identity provides a simplified context signal that allows a shared spiking network to maintain distinct credit traces for different goals while retaining a common long-term synaptic substrate.

An additional biological interpretation of EGP is through the framework of synaptic tagging and capture (STC) and related behavioral tagging mechanisms[46, 58, 59, 60]. In STC, transient synaptic activity establishes local tags that mark recently active synapses, while subsequent plasticity-related signals determine which tagged synapses undergo long-lasting stabilization. Viewed through this lens, the TRAIN phase of EGP resembles synaptic tagging, in which candidate synaptic modifications are generated and stored as transient proposals. The TEST phase provides a form of behavioral evaluation analogous to the availability of plasticity-related signals, providing a behavioral signal that modulates consolidation strength. Consolidation of proposals into the durable synaptic matrix then parallels the stabilization of tagged synapses through late-phase potentiation and memory consolidation[51]. Although EGP is not intended as a mechanistic model of STC, this correspondence suggests that synaptic tagging frameworks may provide a useful biological interpretation of how local plasticity and delayed behavioral outcomes can be linked during ongoing interleaved learning.

Neuromodulatory signals provide a biological motivation for evidence-modulated consolidation. Dopaminergic reward prediction error signals, acetylcholine-dependent state and attention signals, noradrenergic arousal signals, and hippocampal replay events could all influence whether recent synaptic modifications are strengthened, weakened, or stabilized[61, 62, 63, 64]. The TEST phase in EGP abstracts this broader idea: positive performance evidence increases consolidation, while ε=0.05 permits a baseline contribution even without improvement. Temporary candidate expression and restoration are computational devices for implementing this evaluation rather than proposed cellular operations. Biological systems may achieve analogous selection during quiet wakefulness, replay, or sleep, when recent experiences are reactivated and evaluated in relation to reward, novelty, or task relevance[34, 35]. Thus, EGP provides a computational bridge between online synaptic eligibility mechanisms and slower consolidation processes that may operate outside the immediate behavioral episode.

An important feature of the target-context EGP experiments is that learning occurred under an interleaved multi-goal protocol rather than a fixed start-to-goal mapping.

Across 2000 sequential episodes, starting locations were randomized while synaptic weights persisted across episodes. The network therefore had to learn target-selective navigation policies that generalized across many initial conditions rather than memorizing specific trajectories. Within each episode, target contexts were repeatedly alternated during TRAIN and TEST phases, requiring the system to maintain and update multiple navigation policies over extended periods of learning. The independent fixed-weight experiment then tested how much of this multi-goal navigation competence persisted without ongoing plasticity. This setting is particularly relevant biologically, as animals routinely encounter familiar goals from novel locations and must integrate new experiences while preserving previously acquired goal memories. The present design therefore tests sustained learning of multiple recurring goals under changing starting conditions. Interleaving provides continued rehearsal of the recurring goals and does not test the central sequential continual-learning problem of catastrophic forgetting. A complementary protocol withholding earlier goals before later retesting would be required to examine that problem and the stability-adaptability balance [65].

The results also clarify the role of reward shaping in navigation learning. Stepwise rewards based on movement toward or away from the target provided a useful training signal and generally correlated with target-directed behavior. However, reward and target hits were not identical measures of performance. An agent could accumulate positive reward while following inefficient trajectories, or achieve repeated target hits while receiving lower average reward due to oscillatory or circuitous paths. Interpreting target-dependent performance therefore benefits from considering reward, contact, time and dwell together, particularly where individual goals remain difficult.

Several opportunities for extending the model remain. The current architecture deliberately uses simplified spatial and target-related populations [66, 67] and does not include explicit hippocampal recurrence, theta organization, replay or neuromodulator-specific dynamics [68]. Goal identity is supplied externally; future work can test whether comparable context signals emerge endogenously [69, 70, 71]. Residual target heterogeneity and wrong-target occupancy motivate further architectural and representation studies. The eight-pair long-duration and independent-evaluation experiments provide replicated evidence but limit precision, and the fixed-network maps do not establish between-network generality. The obstacle-free arena and dense distance feedback make this a simplified task that could be solved by a direct goal-seeking feedback controller. The controlled setting allows learning-rule behavior to be examined in a spiking circuit; it does not establish general navigation capability. Sparse reward, detours, changing goals and controlled computational-efficiency comparisons remain important tests for future work.

Future work should extend this framework to moving targets and changing goal contingencies, investigate neural mechanisms capable of generating context representations endogenously, and incorporate more biologically detailed consolidation processes such as replay-driven or sleep-like evaluation of candidate synaptic modifications[32]. More broadly, the principles underlying EGP, including rapid local eligibility, slower evidence-modulated consolidation, and context-conditioned consolidation, may prove useful beyond spatial navigation, including continual learning in spiking networks, neuromorphic systems, and biologically grounded reinforcement learning architectures[72, 73, 74, 75, 76, 77, 78, 79, 80].

Global EGP supports durable multi-goal learning by separating rapid synaptic proposals from slower evidence-modulated consolidation. Its target-context extension improves acquisition and conditional target selectivity, with component evidence for additional need-dependent consolidation scaling. Gain controls clarify the role of consolidation strength, while the matched benchmark shows how faster reward acquisition can differ from greater retained target-reaching competence. Together, these findings establish EGP as a testable computational framework for context-sensitive, evidence-modulated plasticity in spiking neural networks and motivate its evaluation in other multi-context learning problems.

## Data Availability Declaration

Code associated with this study is available at https://github.com/NathanKlineInstitute/NavNetEGP, together with the data and analysis materials supporting the reported results. A model package with runnable examples and pretrained weights is deposited in ModelDB (https://modeldb.science/2041698).

## Author Contributions Statement

SN, HH, GU, CE, HA, PF, DB contributed to conceptualization. SN, HH, GU, CE, HA developed the model, performed simulations, and ran data analysis. SN, HH, HA, PF, DB wrote the manuscript. All authors reviewed the manuscript.

## Supplementary material

Supplementary Methods and Results provide neuron equations, EGP pseudocode, complete map examples, cohort and seed-level tables, component diagnostics, and computational resources. Supplementary **Figs. S1-S2** show weight and clipping dynamics; **S3-S8** show coordinate coverage and complete maps; **S9** provides the enlarged learning curves integrated into main **Fig. 12A**; **S10** retains target-specific component results; and **S11** distinguishes raw phase rewards from internal consolidation quantities. Supplementary Methods **S10**, **Figs. S12-S16** and **Tables S17-S18** provide the gain controls, matched STDP/RL benchmark, coefficient and update diagnostics, and seed-level frozen navigation outcomes.

