## supplement for "Evidence-Gated Plasticity and Context-Dependent Consolidation for Multi-Goal Learning in Spiking Neural Networks"

### Supplementary Methods and Results

#### S1. Event-based neuron and synapse dynamics

This supplement accompanies “Evidence-Gated Plasticity and Context-Dependent Consolidation for Multi-Goal Learning in Spiking Neural Networks.” It presents the model equations, algorithmic details, analysis conventions, and supporting results. Main Tables 1-3 report parameter values and population sizes.

For synaptic voltage state S and elapsed time Δt between events, the update is $S(t^{-})=S(t_0)EXP(-\Delta t/\tau)$. The AHP state follows the same decay. EXP denotes a lookup-table approximation with 5000 intervals over [−20,0]. Excitatory and inhibitory states whose magnitude falls below the numerical epsilon are set to zero. Membrane deviation Vdev is the sum of the synaptic states and AHP; absolute voltage is V=clip(RMP+Vdev, −90,0). These state updates are applied analytically at event times.

$$\Delta S_{AMPA}=w_{AMPA}\left(1-Vdev/65\right)$$

$$\Delta S_{NMDA}=w_{NMDA}\,M\left(RMP+Vdev\right)\left(1-Vdev/90\right)$$

$$\Delta S_{GABA}=-w_{GABA}\left(1-Vdev/\left(-15\right)\right)$$

The voltage-dependent magnesium factor is M(v)=maxnmc/[1+exp(−0.062v)mg/mg0]. Reversal-dependent sign conditions and event ordering determine which synaptic increments are applied; main Table 1 gives the time constants of the proximal and distal synaptic states. A spike requires V>VTHC and V≤Vblock while the neuron is not absolutely refractory. Following a spike, AHP decreases by ahpwt. In the nonadditive relative-refractory branch, VTHC=VTH+RRWght(Vblock−VTH), and this value is stored as VTHR. Absolute refractoriness ends after refrac−AHP·AHP2REF; the relative threshold then decays toward VTH with time constant τRR. Spatial input populations retain relative refractoriness even when their AHP increment is zero.

Reward-modulated plasticity is restricted to the action-specific association-to-motor AMPA projections: EAN→EMN, EAS→EMS, EAE→EME and EAW→EMW, with 7680 plastic synapses overall. A causal pre-before-post interval of at most 100 ms establishes a 50-ms eligibility pulse. Reinforcement is evaluated at the 50-ms action interval. The base learning increment 0.045 and bounds 0-12 are multiplied by the excitatory gain 4.4 in the controlled EGP experiments, giving 0.198 and 0-52.8. Synapses associated with the selected motor direction receive the signed critic; those associated with the opposing direction receive the opposite sign multiplied by 0.25. The single- and multi-target STDP/RL demonstrations and the controlled EGP comparisons use the protocols described in the main Methods; the matched 200-episode learning-rule benchmark is detailed in S10.

Motor output is determined by independent east-versus-west and north-versus-south comparisons, allowing eight moves or no movement. North decreases y. The arena clips x,y to [8,91]. Raw reward is the decrease in Euclidean distance from the attempted movement divided by $Dnorm=\sqrt{(W^2+H^2)}$, the simulator’s fixed distance-normalization scale, with W=H=100 pixels. Raw reward uses the attempted position before clipping, whereas the critic’s distance factor and exact-contact bonus use the clipped position. These separate signals are defined in the main Methods. The learning experiments use this fixed critic, without a learned value function or temporal-difference prediction error.

#### S2. EGP state updates, map definitions and statistical conventions

EGP organizes plasticity into candidate generation, temporary expression, and durable consolidation. Global EGP maintains a single proposal store and consolidation coefficient, whereas target-context EGP maintains goal-specific proposals and evidence. During TEST, the combined TRAIN proposals are expressed together, allowing the network to evaluate the aggregate candidate while retaining goal-indexed consolidation weights without isolating each proposal’s causal contribution.

```
For each TRAIN/TEST cycle:
  Save the durable weight matrix W.
  TRAIN: W drives behavior; eligible changes accumulate in P or P[k].
  At target switches, store the old target’s live proposal.
  At TRAIN end, store the last proposal and save all TRAIN proposals.
  TEST: express clip(W + sum_k P[k]) for the whole phase.
  Accumulate TEST evidence; keep TEST-only proposals separate.
  Restore the saved W and discard TEST-only proposals.
  Calculate global or target-specific evidence and coefficients.
  Commit clip(W + sum_k beta[k] P[k]); clear transient proposals.
```

TRAIN→TEST restores the configured TRAIN start, but neural and input-noise state continue. TEST→TRAIN continues from the preceding endpoint, and target switches alone do not reset position. Recent target performance q is tracked as a phase-level exponential mean of raw reward ($\alpha$=0.1), initialized at the first observation; unseen targets have need 1. Need is bounded at 0.75-2. Both full variants use need-weighted critic evidence; global evidence is a phase sum and context evidence a per-target mean. The normalized improvement uses the exact zero-denominator guard in the main Methods. $\eta$=1 and $\varepsilon$=0.05 retain nonzero baseline consolidation; full context EGP adds the need multiplier to $\beta$. The experiments used fixed $\varepsilon$, need-range, and phase-duration settings; their sensitivity can be evaluated in future work. Proposal storage is 60 KiB globally or 300 KiB for five contexts at float64 precision, excluding other state.

Firing rates were estimated by dividing spike counts by occupancy in 5×5-pixel bins. Each of 256 eligible bins has 50 visits of 50 ms; 144 padding bins are NA. Information is computed on raw rates as $I=\Sigma_i p_i(\lambda_i/\bar{\lambda})\log_2(\lambda_i/\bar{\lambda})$, with $\bar{\lambda}=\Sigma_i p_i \lambda_i$. Zero-rate terms contribute zero when $\bar{\lambda}>0$, and silent cells have undefined information. Display and field detection use Gaussian smoothing with $\sigma$=1 bin and normalization over the eligible region. Fields are ≥20% of the cell peak, four-connected and at least four bins; bin area is 25 pixel$^2$. Alternating visits form two sets of 25 visits/bin; Pearson correlation between the smoothed odd/even maps measures pattern reproducibility. This split-half reliability measure complements spatial information and is distinct from gridness. Undefined correlations are recorded as NA rather than zero.

Statistical inference uses paired network seeds as the independent units. The specified endpoints were analyzed using exact two-sided sign-flip tests and 100000-resample percentile intervals. Bootstrap generators were RandomState 20260817 for the replication primary, RandomState 260817 for long training, RandomState 270927 for independent evaluation, and PCG64 20260929 for the component experiment. Confidence intervals are unadjusted estimation summaries, and Holm adjustments use the specified test-family sizes. For n=8, the minimum exact p is 2/256=0.0078125, larger than the first Holm-12 and Holm-15 thresholds, so these larger families have limited exact-test resolution. Effect estimates and confidence intervals complement the adjusted tests, especially in these larger families. Undefined ratios are recorded as NA; comparisons requiring all eight pairs are reported as unavailable if only seven pairs are defined.

#### S3. Supporting short-run and STDP/RL results

**Table S1. Early TEST reward, episodes 0-99.** The independently specified 30-pair replication is reported alongside the initial 10-pair cohort and their pooled 40-pair summary. Differences are context−global, with unadjusted 95% intervals.

| Cohort / paired n | Global / context | Difference [95% CI] | Exact p; context higher |
|---|---|---|---|
| Initial cohort / 10 | 0.0022 / 0.0117 | 0.0095 [0.0047, 0.0155] | 0.00195; 10/10 |
| Prospective replication / 30 | 0.004 / 0.009 | 0.005 [0.0015, 0.0088] | 0.00999; 24/30 |
| Pooled support / 40 | 0.0035 / 0.0096 | 0.0061 [0.0031, 0.0093] | 0.00019; 34/40 |

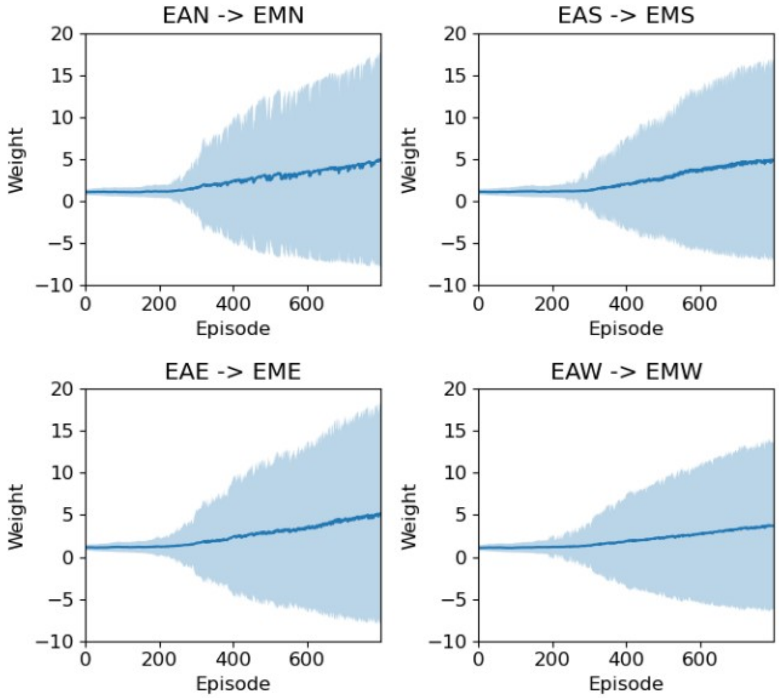


**Figure S1. Synaptic weight dynamics during the 800 single-target learning episodes.** The four panels show the direction-specific projections. Lines show mean weight and bands ±SD across synapses (7680 total). Shading may extend below zero when the SD exceeds the mean, although individual synaptic weights remain nonnegative. The panels illustrate how synaptic weights and their dispersion evolved during single-target learning in this network.

**Table S2. Complete multi-target STDP/RL post-learning evaluation, with plasticity disabled.** One network, 500 episodes, 100 per target, each 187.5 s. These complete-stage results complement the last-100 combined-episode windows shown in main Figs. 4 and 6. The STDP/RL experiment is descriptive; the matched global-versus-target-context EGP comparisons are presented separately.

| Group / n | Successes / fraction | Restricted time (actions) | Correct / raw wrong / conditional wrong dwell |
|---|---|---|---|
| All / 500 | 278 / 0.556 | 2158.98 | 0.3344 / 0.1075 / 0.2434 |
| Target 0 / 100 | 54 / 0.54 | 2320.15 | 0.306 / 0.1557 / 0.3372 |
| Target 1 / 100 | 63 / 0.63 | 1747.62 | 0.4704 / 0.0125 / 0.026 |
| Target 2 / 100 | 65 / 0.65 | 2110.18 | 0.4223 / 0.0836 / 0.1652 |
| Target 3 / 100 | 50 / 0.5 | 2256.89 | 0.317 / 0.0998 / 0.2394 |
| Target 4 / 100 | 46 / 0.46 | 2360.05 | 0.1561 / 0.1862 / 0.5439 |

Among the 500 post-learning STDP/RL episodes, 278 reached the target (55.6%); the reached-only median first-passage time was 498 actions and mean physical path inefficiency was 26.65.

#### S4. Long-duration target, navigation and selectivity results

In the prospective short EGP cohort, the final-100 reward difference was 0.0252. Whole-training correct dwell was 1.5%/2.3%, conditional wrong dwell 74.7%/67.2%, and raw wrong dwell 4.6%/4.8% for global/target-context EGP. These 7.5-s target blocks devote substantial time to approaching a goal, making reward acquisition the most informative short-run outcome. The longer protocol below provides complementary measures of navigation success and target selectivity.

**Table S3. All-episode and final-100 target reward results.** Effects are context−global. All-episode and final-100 target comparisons each use their own Holm-5 family; the late all-target summary is nominal.

| Window / target | Global / context | Difference [95% CI] | Exact p / Holm p |
|---|---|---|---|
| 0-1999 / t0 | 0.0567 / 0.1436 | 0.0869 [0.0703, 0.1043] | 0.00781 / 0.0391 |
| 0-1999 / t1 | 0.2658 / 0.3315 | 0.0656 [0.0372, 0.0975] | 0.00781 / 0.0391 |
| 0-1999 / t2 | 0.2582 / 0.2849 | 0.0268 [0.0144, 0.0379] | 0.0156 / 0.0391 |
| 0-1999 / t3 | 0.2732 / 0.332 | 0.0588 [0.0371, 0.0783] | 0.00781 / 0.0391 |
| 0-1999 / t4 | 0.0956 / 0.1653 | 0.0697 [0.0394, 0.1054] | 0.00781 / 0.0391 |
| 1900-1999 / all | 0.2674 / 0.2984 | 0.031 [0.0208, 0.0422] | 0.00781 / NA |
| 1900-1999 / t0 | 0.1212 / 0.1814 | 0.0602 [0.0279, 0.0952] | 0.0156 / 0.0625 |
| 1900-1999 / t1 | 0.3721 / 0.3962 | 0.0242 [-0.01, 0.0666] | 0.344 / 0.773 |
| 1900-1999 / t2 | 0.3211 / 0.3192 | -0.0019 [-0.018, 0.015] | 0.828 / 0.828 |
| 1900-1999 / t3 | 0.3492 / 0.3932 | 0.044 [0.0244, 0.0632] | 0.00781 / 0.0391 |
| 1900-1999 / t4 | 0.1734 / 0.202 | 0.0286 [-0.0113, 0.0706] | 0.258 / 0.773 |

**Table S4. Final-100 internal-TEST navigation (30-s horizon), with the complete four-endpoint Holm family.** The outcomes summarize repeated 30-s TEST blocks from eight paired networks, with paired-network seeds serving as the units of inference.

| Endpoint | Global / context | Difference [95% CI] | Exact p / Holm p |
|---|---|---|---|
| Contact fraction | 0.5735 / 0.6663 | 0.0929 [0.0517, 0.1347] | 0.00781 / 0.03125 |
| Restricted FPT (s) | 21.12 / 19.57 | -1.54 [-2.2, -0.9157] | 0.00781 / 0.03125 |
| Reached FPT (s) | 14.5 / 14.65 | 0.149 [-0.8056, 1.2] | 0.789 / 1 |
| Reached path ineff. | 5.76 / 6.02 | 0.2578 [-0.4599, 1.06] | 0.586 / 1 |

**Table S5. Late target-selectivity measures.** Intervals are unadjusted and p values are nominal, without multiplicity adjustment. Correct and raw wrong dwell use all observed time, conditional wrong dwell only time near any goal.

| Endpoint | Global / context | Difference [95% CI] | Exact p |
|---|---|---|---|
| Correct dwell | 0.2734 / 0.3274 | 0.0541 [0.0299, 0.0793] | 0.00781 |
| Minimum diagonal | 0.0817 / 0.0971 | 0.0154 [-0.0144, 0.043] | 0.32 |
| Maximum off-diagonal | 0.0903 / 0.0898 | -4.75e-04 [-0.0229, 0.0241] | 0.977 |
| Diagonal/off-diagonal ratio | 18.78 / 21.99 | 3.21 [1.99, 4.39] | 0.00781 |
| Conditional wrong dwell | 0.2057 / 0.1833 | -0.0224 [-0.0305, -0.016] | 0.00781 |
| Raw wrong dwell | 0.0632 / 0.0646 | 0.0014 [-0.0034, 0.0075] | 0.727 |

#### S4 continued. Candidate clipping and retained-weight bounds

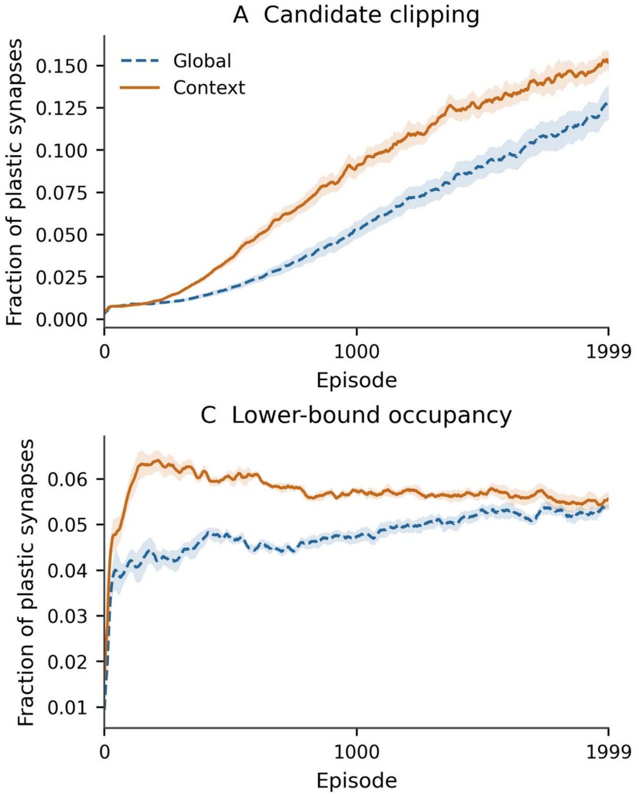


**Figure S2. Candidate and commit clipping and durable-weight bound occupancy during long training.** Curves span all 2000 episodes across eight paired seeds. Five boundary values are averaged per episode, followed by the fixed 25-episode seed-first display smoother; shading is between-seed SEM. Candidate expression and durable commitment are distinct stages. Clipping is measured from synapse-event counts, whereas bound occupancy is measured in the saved durable weights. These panels show how clipping and retained-weight bounds evolve across long-duration learning, complementing the behavioral comparisons.

Supplementary Figure S11 traces the progression from raw phase reward to transformed evidence, consolidation coefficients, synaptic proposals, durable weight changes, and cancellation. Need-weighted critic evidence can differ from raw environmental reward, and normalized improvements can be large when TRAIN evidence approaches zero. The summaries include these large relative changes. Durable weight changes are measured after weight restoration and clipping, which distinguishes committed changes from the proposed update. L1 cancellation describes destructive overlap; the corresponding L2 aggregation loss also reflects vector geometry and is therefore a complementary quantity.

The final-100 navigation analysis included 2500 TEST blocks per seed/arm (40000 total), with failures censored at the 30-s horizon. Physical path inefficiency is (actual path−shortest octile path)/shortest octile path, conditional on contact and a positive optimum. A start already on the active target counts as success but has undefined path inefficiency. Because each arm contributes its own set of successful blocks, reached-only metrics are interpreted alongside all-block contact success and restricted mean first-passage time. Neither reached-only path inefficiency nor reached-only time showed adjusted evidence of an arm difference.

#### S5. Independent frozen-weight evaluation: complete secondary families

Independent fixed-weight evaluation demonstrated durable navigation in both algorithms (main Fig. 11). At the final checkpoint, success was 56.0% versus 67.8%, a difference of 11.8 percentage points (95% CI 2.3-22.2; exact p=0.063), with a directional target-context advantage. All evaluations use fresh starts, durable weights only, and learning/proposals disabled. Tables S6-S8 report the seven-endpoint, five-target and eight-checkpoint families separately. Each comparison has eight network pairs; the 187.5-s horizon is distinct from internal TEST blocks. PRETRAIN success was 17.8% in both arms and serves as the empirical pretraining baseline.

**Table S6. All seven final-checkpoint secondary endpoints.** The seven endpoints comprise one Holm-adjusted family.

| Endpoint | Global / context | Difference [95% CI] | Exact p / Holm p |
|---|---|---|---|
| Restricted FPT (s) | 104.32 / 84.57 | -19.75 [-32.75, -8.86] | 0.0156 / 0.0938 |
| Reached path ineff. | 27.08 / 25.98 | -1.1 [-9.65, 6.95] | 0.805 / 1 |
| Reached action ineff. | 27.58 / 26 | -1.58 [-10.34, 7.06] | 0.734 / 1 |
| Correct dwell | 0.2481 / 0.3256 | 0.0776 [0.0468, 0.1075] | 0.00781 / 0.0547 |
| Conditional wrong dwell | 0.1392 / 0.0804 | -0.0588 [-0.1177, -0.0049] | 0.148 / 0.594 |
| Raw wrong dwell | 0.0356 / 0.0281 | -0.0075 [-0.0282, 0.0092] | 0.547 / 1 |
| Raw reward/action | -2.88e-04 / -1.75e-04 | 1.12e-04 [4.61e-05, 1.82e-04] | 0.0312 / 0.156 |

**Table S7. All five final target-success comparisons.** All five targets form one Holm-adjusted family.

| Endpoint | Global / context | Difference [95% CI] | Exact p / Holm p |
|---|---|---|---|
| target t0 success | 0.7125 / 0.9625 | 0.25 [0.0125, 0.575] | 0.25 / 1 |
| target t1 success | 0.6375 / 0.6125 | -0.025 [-0.175, 0.0875] | 0.938 / 1 |
| target t2 success | 0.4625 / 0.5875 | 0.125 [-0.1, 0.325] | 0.359 / 1 |
| target t3 success | 0.8375 / 0.9125 | 0.075 [-0.0625, 0.225] | 0.5 / 1 |
| target t4 success | 0.15 / 0.3125 | 0.1625 [0.0625, 0.275] | 0.0312 / 0.156 |

**Table S8. All eight trained-checkpoint success comparisons.** The eight trained checkpoints form one Holm-adjusted family; PRETRAIN is the separate baseline.

| Endpoint | Global / context | Difference [95% CI] | Exact p / Holm p |
|---|---|---|---|
| Episode 0 | 0.1675 / 0.175 | 0.0075 [-0.0125, 0.0275] | 0.688 / 1 |
| Episode 99 | 0.1625 / 0.16 | -0.0025 [-0.04, 0.0325] | 1 / 1 |
| Episode 199 | 0.1325 / 0.175 | 0.0425 [-0.025, 0.115] | 0.359 / 1 |
| Episode 499 | 0.25 / 0.54 | 0.29 [0.1899, 0.3876] | 0.00781 / 0.0625 |
| Episode 999 | 0.4625 / 0.6225 | 0.16 [0.0425, 0.2575] | 0.0469 / 0.234 |
| Episode 1499 | 0.5575 / 0.695 | 0.1375 [0.065, 0.215] | 0.0156 / 0.109 |
| Episode 1799 | 0.555 / 0.7075 | 0.1525 [0.0525, 0.285] | 0.0234 / 0.141 |
| Episode 1999 | 0.56 / 0.6775 | 0.1175 [0.0225, 0.2225] | 0.0625 / 0.25 |

#### S6. Spatial sampling and all-cell representation summaries

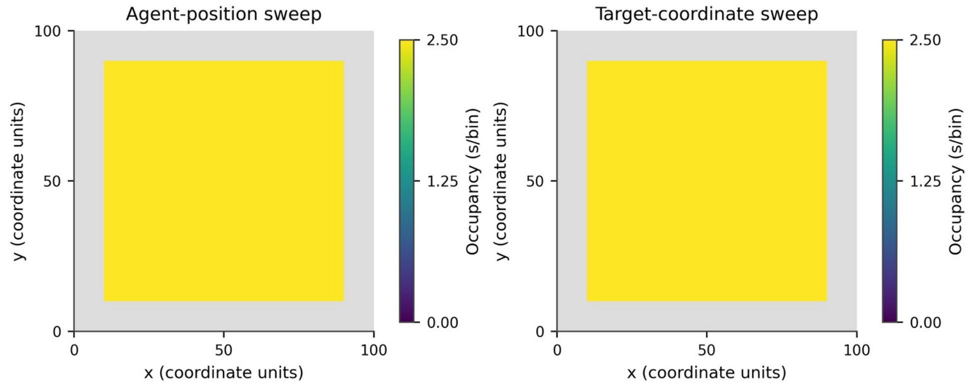


**Figure S3. Occupancy in the separate agent-position and target-coordinate sweeps.** Both panels use viridis with shared limits of 0-2.5 s/bin. Each eligible bin has 2.5 s occupancy (50 visits×50 ms); gray denotes 144 unobserved padding bins, distinct from sampled bins with zero firing. Odd/even halves have 25 visits per eligible bin. All plastic weights are fixed.

**Table S9. All-cell map summaries: median [IQR] and defined n.** Every population retains all 384 cells in the inventory, with silent and undefined cases reported explicitly.

| Population | Rate Hz | Information bits/spike | Odd/even r | Silent / retained |
|---|---|---|---|---|
| EGrid | 1.21 [0.9359,1.56]; n=384 | 3.09 [2.75,3.42]; n=384 | 0.7475 [0.5191,0.8476]; n=384 | 0/384 |
| EPlace | 0.3672 [0.1992,0.598]; n=384 | 4.66 [4,5.38]; n=368 | 0.3934 [0.1696,0.5747]; n=362 | 16/384 |
| EV | 1.19 [0.9359,1.48]; n=384 | 3.09 [2.76,3.42]; n=384 | 0.7188 [0.4813,0.8254]; n=384 | 0/384 |
| EVPlace | 0.3875 [0.1953,0.5469]; n=384 | 4.68 [4.19,5.42]; n=376 | 0.3197 [0.1159,0.5205]; n=363 | 8/384 |

**Table S10. Field summaries across all cells.** Areas are pixel²; field thresholds are specified in S2. These operational fields summarize spatial structure under the stated thresholds and are not used as physiological cell classifications.

| Population | Field count: median [IQR] | Largest area: median [IQR] |
|---|---|---|
| EGrid | 4 [3,4]; n=384 | 650 [550,800]; n=384 |
| EPlace | 3 [2,4]; n=384 | 525 [400,750]; n=384 |
| EV | 4 [3,4]; n=384 | 675 [550,850]; n=384 |
| EVPlace | 3 [2,4]; n=384 | 575 [450,775]; n=384 |

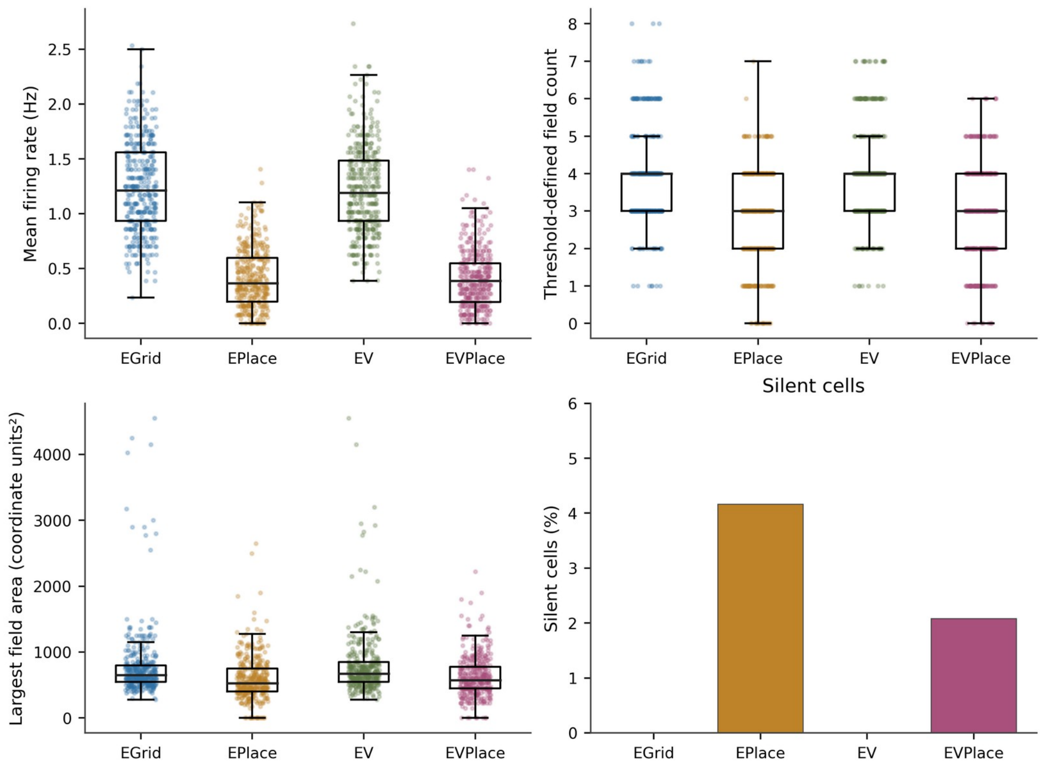


**Figure S4. All-cell distributions of firing rate, threshold-defined field count and largest-field area, plus silent-cell fractions.** Dots represent all 384 cells per population for each displayed metric; boxes show median/IQR with 1.5-IQR whiskers. Silent-cell counts are 0, 16, 0 and 8 for EGrid, EPlace, EV and EVPlace, respectively, each out of 384. These 1536 neurons were measured in one fixed network; the paired behavioral experiments provide the separate network-level replication. EGrid/EPlace use agent position; EV/EVPlace use target coordinates.

#### S6 continued. EGrid examples

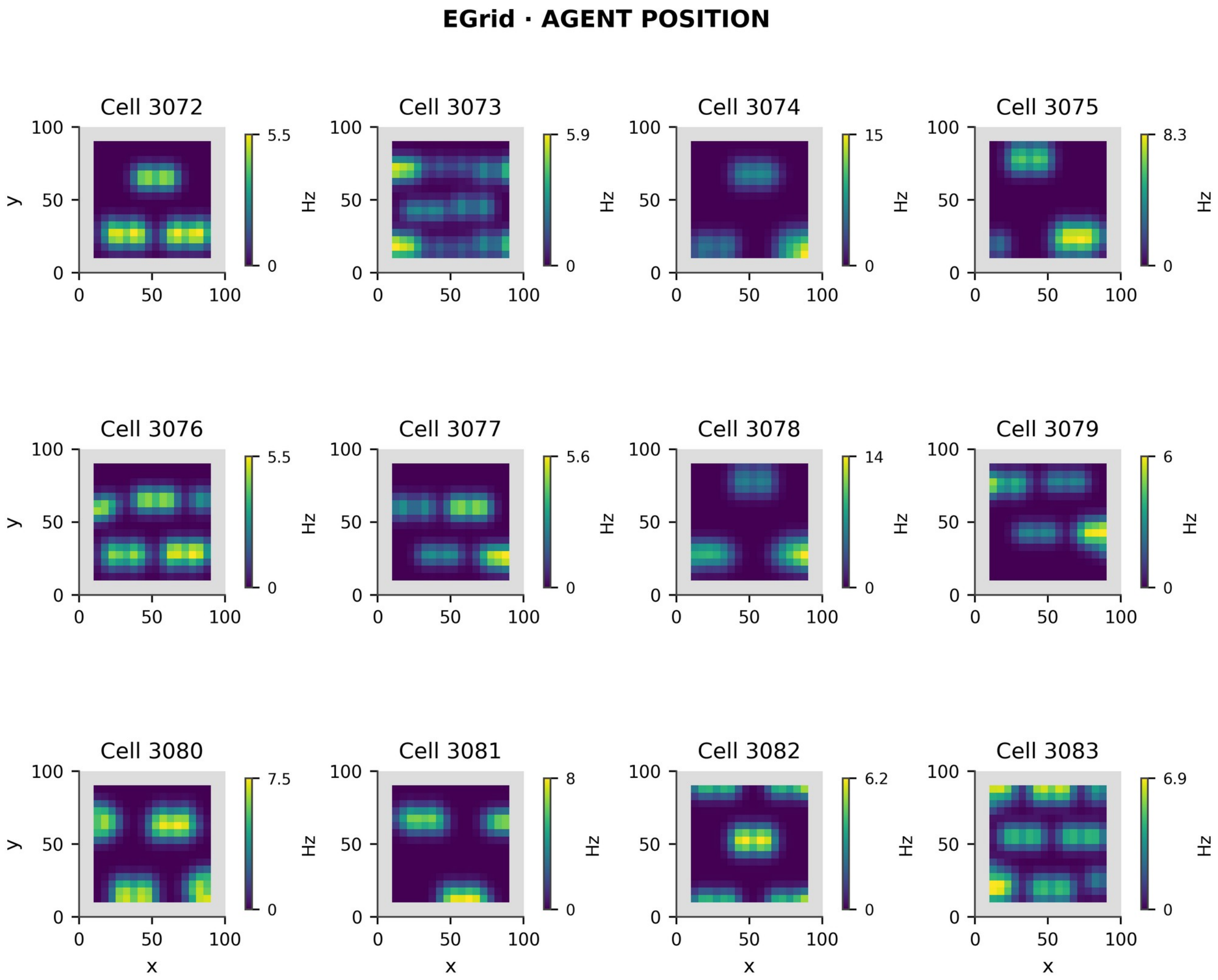


**Figure S5. All 12 preselected EGrid cells, ordered by identifier before viewing map outcomes.** Maps encode agent position. The first 12 cell identifiers were chosen before map inspection, providing a fixed set of examples. Each map has a separate viridis colorbar in Hz, using that cell's own scale. Display smoothing is σ=1 bin; gray is unobserved padding. Silent cells remain present and use a 0-1 Hz display scale. The main Fig. 2 shows the first two of these cells, and the population summaries include all 384. All maps were measured before training in the same fixed seed-1 network.

#### S6 continued. EPlace examples

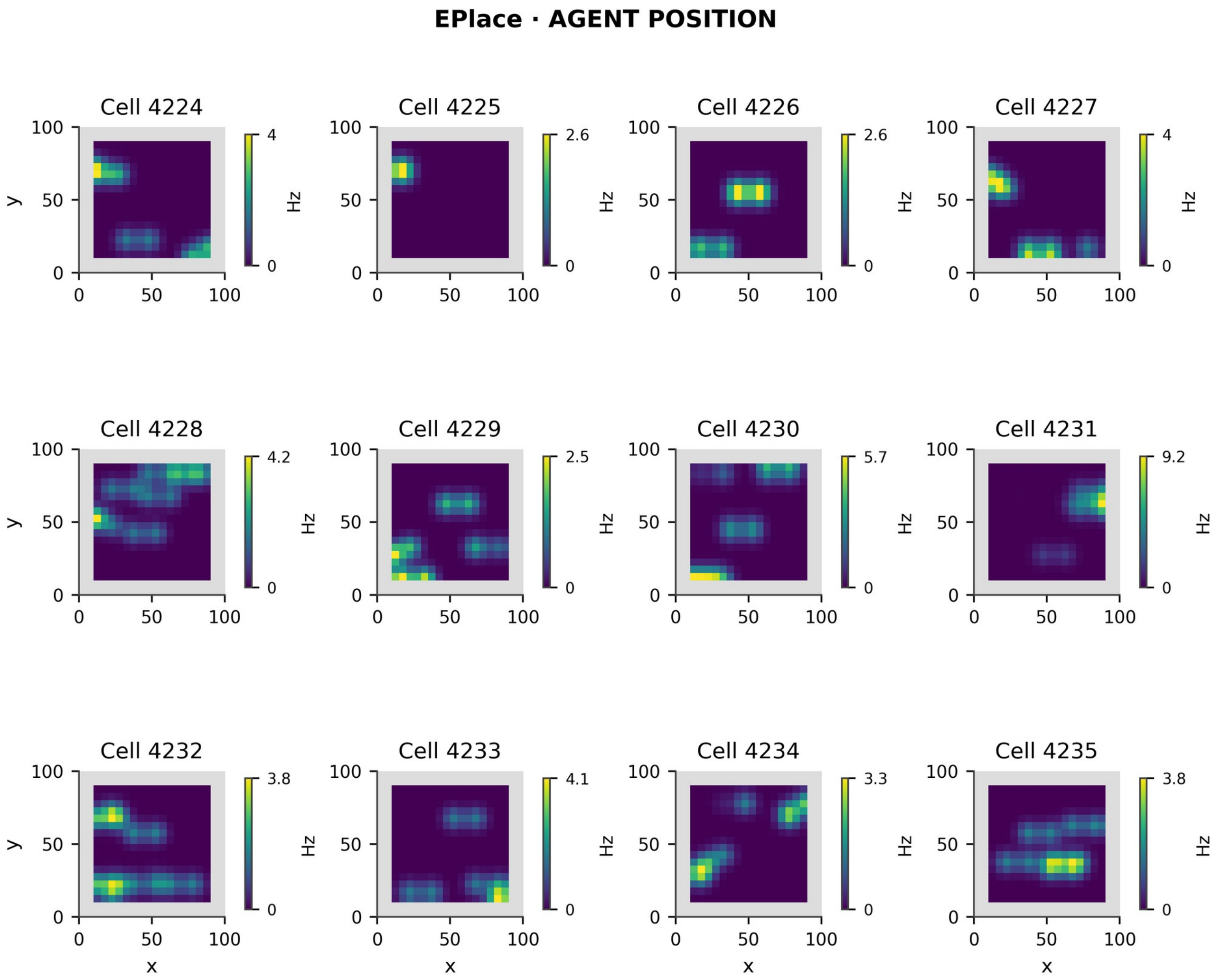


**Figure S6. All 12 preselected EPlace cells, ordered by identifier before viewing map outcomes.** Maps encode agent position. The first 12 cell identifiers were chosen before map inspection, providing a fixed set of examples. Each map has a separate viridis colorbar in Hz, using that cell's own scale. Display smoothing is σ=1 bin; gray is unobserved padding. Silent cells remain present and use a 0-1 Hz display scale. The main Fig. 2 shows the first two of these cells, and the population summaries include all 384. All maps were measured before training in the same fixed seed-1 network.

#### S6 continued. EV examples

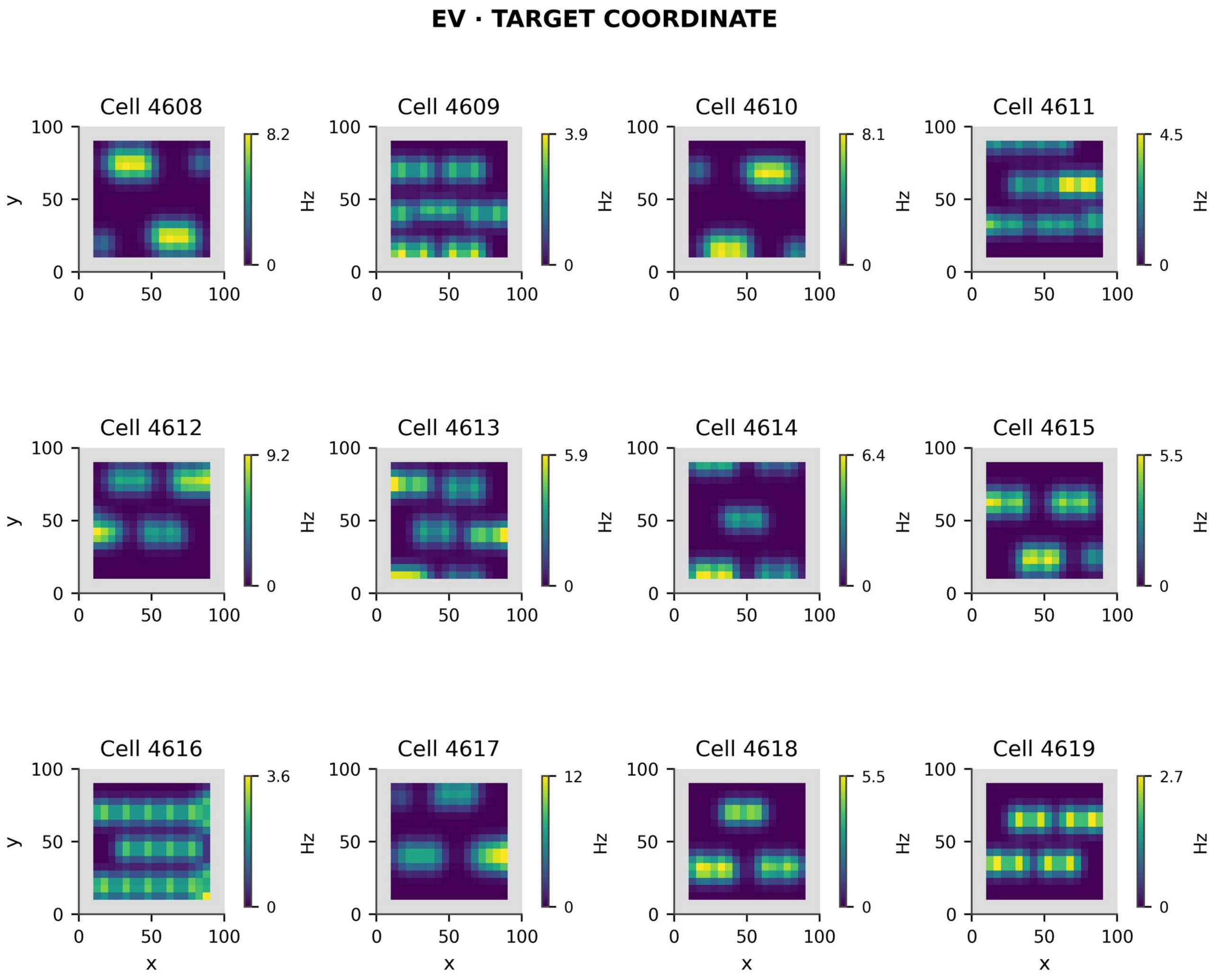


**Figure S7. All 12 preselected EV cells, ordered by identifier before viewing map outcomes.** Maps encode target coordinates. The first 12 cell identifiers were chosen before map inspection, providing a fixed set of examples. Each map has a separate viridis colorbar in Hz, using that cell's own scale. Display smoothing is σ=1 bin; gray is unobserved padding. Silent cells remain present and use a 0-1 Hz display scale. The main Fig. 2 shows the first two of these cells, and the population summaries include all 384. All maps were measured before training in the same fixed seed-1 network.

#### S6 continued. EVPlace examples

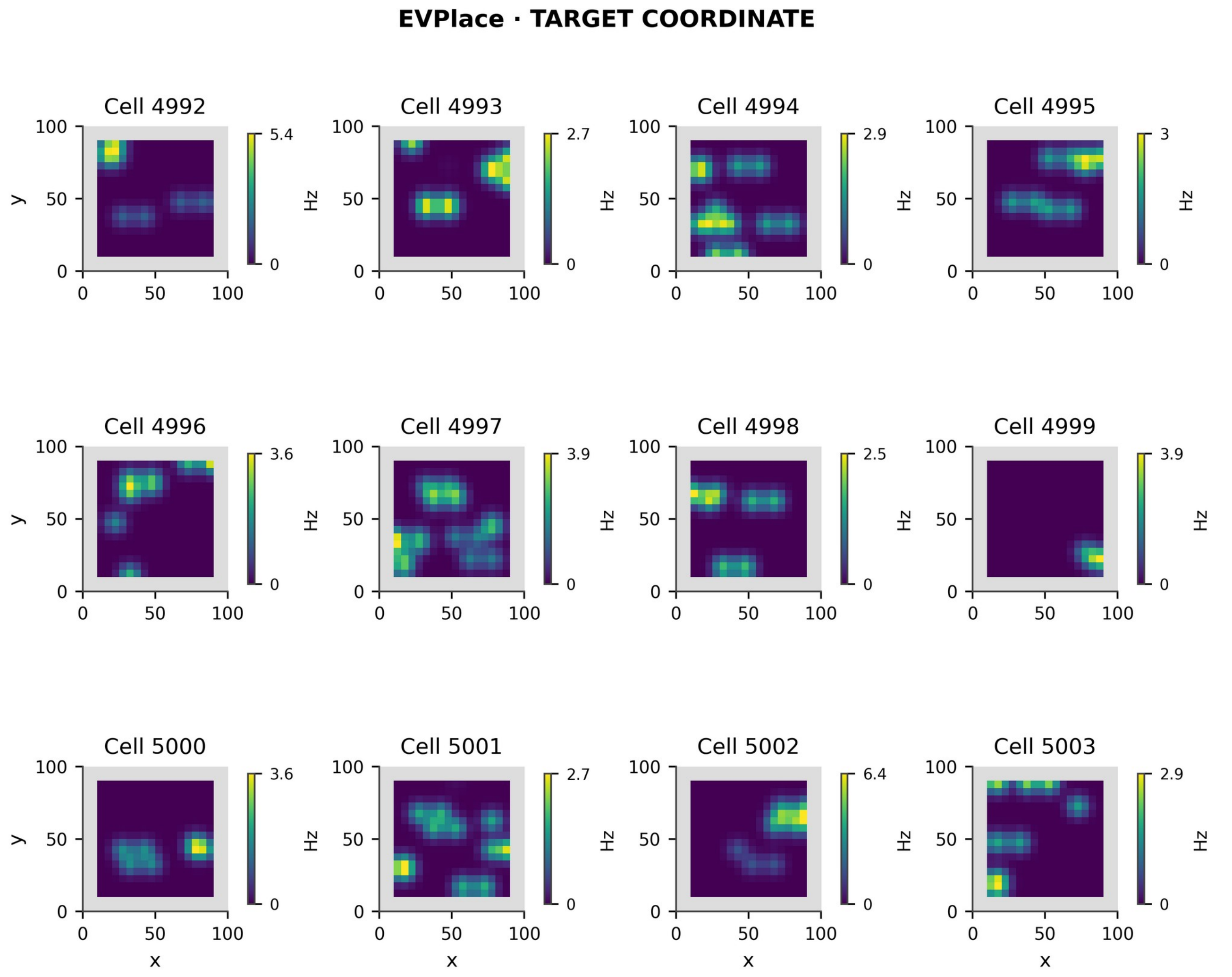


**Figure S8. All 12 preselected EVPlace cells, ordered by identifier before viewing map outcomes.** Maps encode target coordinates. The first 12 cell identifiers were chosen before map inspection, providing a fixed set of examples. Each map has a separate viridis colorbar in Hz, using that cell's own scale. Display smoothing is σ=1 bin; gray is unobserved padding. Silent cells remain present and use a 0-1 Hz display scale. The main Fig. 2 shows the first two of these cells, and the population summaries include all 384. All maps were measured before training in the same fixed seed-1 network.

#### S7. Component learning dynamics and target heterogeneity

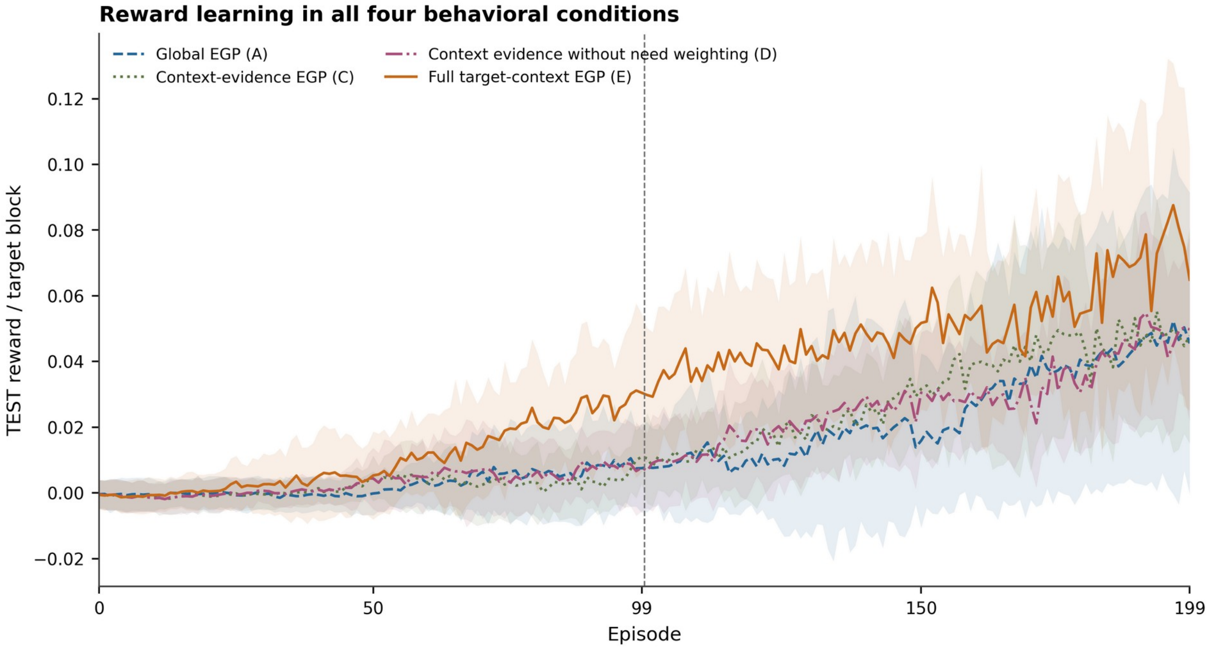


**Figure S9. Enlarged reward-learning curves corresponding to main Figure 12A.** Curves show global EGP (A), context-evidence EGP (C), context evidence without need weighting (D), and full target-context EGP (E) across all 200 episodes and eight network seeds. Curves and bands are unsmoothed mean ± between-seed SD, not confidence intervals. The divider separates episodes 0-99 from 100-199. B is the behaviorally identical segmented shadow inside A, not a fifth stream. The enlarged view resolves overlap among the learning curves also displayed in main Fig. 12A.

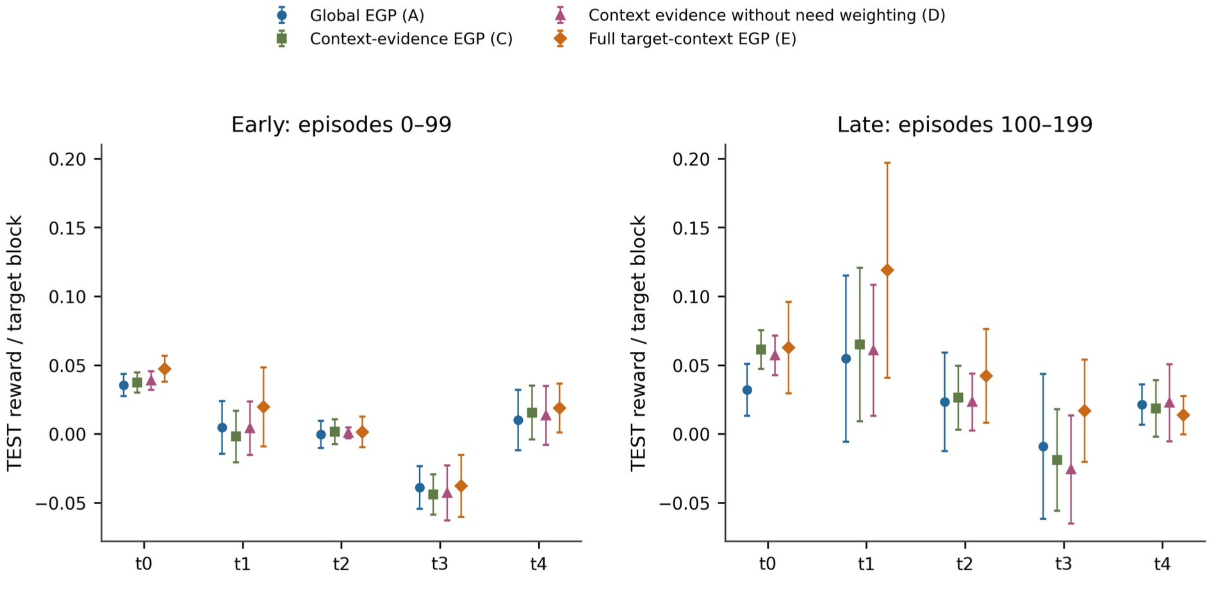


**Figure S10. Target-specific reward in all four behavioral conditions.** Global EGP (A), context-evidence EGP (C), context evidence without need weighting (D), and full target-context EGP (E) are shown for all five targets in early (0-99) and late (100-199) episodes. Points and bars are mean ± between-seed SD, with a shared reward scale. The horizontal categories identify targets rather than time. The panels illustrate early-to-late differences in reward across targets; target-specific and late-window comparisons are descriptive.

#### S7 continued. Reward components and mechanistic tests

In the primary comparison, full target-context EGP exceeded global EGP in early reward by 0.0076 (95% CI 0.0043-0.0114; exact p=0.0078; all eight pairs positive). The three component reward contrasts and 12 mechanistic contrasts are presented in their respective multiplicity families. B equals A behaviorally; its segmented stores use A’s common coefficient. Among the component contrasts, only E−C showed an adjusted reward effect, supporting the added need multiplier on context-specific consolidation. The mechanistic endpoints provide complementary estimates and uncertainty but did not identify a single intermediate process explaining that benefit after Holm-12 correction.

**Table S11. Complete Holm-3 component reward family.**

| Contrast / endpoint | Effect [95% CI] | Exact p | Holm p |
|---|---|---|---|
| C−B / TEST reward | -4.77e-04 [-0.0037, 0.0018] | 0.93 | 0.93 |
| C−D / TEST reward | -0.0011 [-0.003, 1.94e-04] | 0.391 | 0.781 |
| E−C / TEST reward | 0.0081 [0.0039, 0.0126] | 0.00781 | 0.0234 |

**Table S12. Complete Holm-12 mechanistic family.** The 12 mechanistic endpoints comprise one Holm-adjusted family.

| Contrast / endpoint | Effect [95% CI] | Exact p | Holm p |
|---|---|---|---|
| C−B / Durable Δ L1 | -0.3169 [-2.34, 1.96] | 0.797 | 1 |
| C−B / Cancellation L1 | 0.073 [-0.0961, 0.2532] | 0.469 | 1 |
| C−B / Pooled clipping | -9.67e-05 [-4.48e-04, 2.80e-04] | 0.586 | 1 |
| C−B / Context β SD | 0.3144 [0.3055, 0.3267] | 0.00781 | 0.0938 |
| C−D / Durable Δ L1 | -0.1501 [-0.4947, 0.1749] | 0.43 | 1 |
| C−D / Cancellation L1 | -0.0159 [-0.0687, 0.0478] | 0.625 | 1 |
| C−D / Pooled clipping | -4.96e-05 [-1.27e-04, 1.88e-05] | 0.289 | 1 |
| C−D / Context β SD | 0.0067 [8.03e-04, 0.0112] | 0.0625 | 0.438 |
| E−C / Durable Δ L1 | 6.57 [4.92, 8.18] | 0.00781 | 0.0938 |
| E−C / Cancellation L1 | 0.8195 [0.5825, 1.04] | 0.00781 | 0.0938 |
| E−C / Pooled clipping | 1.16e-04 [2.61e-05, 2.03e-04] | 0.0469 | 0.375 |
| E−C / Context β SD | 0.1719 [0.1483, 0.1943] | 0.00781 | 0.0938 |

**Table S13. Early clipping percentages by stage, descriptive only.** Values are means of episode-level fractions within seed, then across eight seeds. These stage-specific values complement the pooled clipping endpoint defined below for inference.

| Condition | Candidate (%) | Commit (%) | Pooled (%) |
|---|---|---|---|
| A | 0.2255 | 0.1956 | 0.2106 |
| C | 0.2128 | 0.1889 | 0.2009 |
| D | 0.2186 | 0.193 | 0.2058 |
| E | 0.2221 | 0.2029 | 0.2125 |

The inferential clipping endpoint is (candidate clipped + commit clipped)/(candidate eligible + commit eligible), calculated per episode and then averaged over episodes 0-99 within seed. Across the 3200 early episodes, each stage has 38400 eligible synapse-events per episode, yielding nonzero pooled denominators throughout. A zero denominator, if encountered, would be recorded as NA rather than zero. Durable Δ L1 and cancellation L1 are in weight units; β dispersion is the SD across context coefficients. The positive E−C cancellation estimate indicates greater cancellation in the full target-context condition.

#### S7 continued. Complete behavioral families and missingness

**Table S14. All five E−A behavioral endpoints (first five rows, Holm-5) and all 15 component-behavior endpoints (remaining rows, Holm-15).** Each family has its own Holm adjustment. Four reached-only comparisons could not be tested because one condition lacked the full required eight-pair set.

| Contrast / endpoint | Effect [95% CI] | Exact p | Holm p | Defined pairs |
|---|---|---|---|---|
| E−A / Contact fraction | -0.0014 [-0.0097, 0.0064] | 0.758 | 1 | 8/8 |
| E−A / Restricted FPT (s) | 0.0117 [-0.0222, 0.0504] | 0.562 | 1 | 8/8 |
| E−A / Reached action ineff. | NA [NA, NA] | NA | NA | 7/8; NA |
| E−A / Reached path ineff. | NA [NA, NA] | NA | NA | 7/8; NA |
| E−A / Conditional wrong dwell | 0.0178 [-0.037, 0.0675] | 0.562 | 1 | 8/8 |
| C−B / Contact fraction | -0.0034 [-0.0079, 6.50e-04] | 0.188 | 1 | 8/8 |
| C−B / Restricted FPT (s) | 0.0155 [-0.0036, 0.0387] | 0.227 | 1 | 8/8 |
| C−B / Reached action ineff. | NA [NA, NA] | NA | NA | 7/8; NA |
| C−B / Reached path ineff. | NA [NA, NA] | NA | NA | 7/8; NA |
| C−B / Conditional wrong dwell | 0.0226 [-0.0241, 0.0638] | 0.367 | 1 | 8/8 |
| C−D / Contact fraction | -0.0037 [-0.0064, -8.00e-04] | 0.0625 | 0.875 | 8/8 |
| C−D / Restricted FPT (s) | 0.0194 [0.0062, 0.0313] | 0.0391 | 0.586 | 8/8 |
| C−D / Reached action ineff. | 1.25 [-1.79, 4.44] | 0.477 | 1 | 8/8 |
| C−D / Reached path ineff. | 1.03 [-1.28, 3.47] | 0.438 | 1 | 8/8 |
| C−D / Conditional wrong dwell | 0.0118 [8.44e-04, 0.0219] | 0.0859 | 1 | 8/8 |
| E−C / Contact fraction | 0.002 [-0.0038, 0.0066] | 0.531 | 1 | 8/8 |
| E−C / Restricted FPT (s) | -0.0038 [-0.0255, 0.0268] | 0.844 | 1 | 8/8 |
| E−C / Reached action ineff. | -1.23 [-3.75, 1.13] | 0.383 | 1 | 8/8 |
| E−C / Reached path ineff. | -0.8527 [-2.83, 1.1] | 0.453 | 1 | 8/8 |
| E−C / Conditional wrong dwell | -0.0048 [-0.0477, 0.0448] | 0.859 | 1 | 8/8 |

Contact success and conditional wrong dwell are fractions; restricted first-passage time is in seconds. Reached action inefficiency is excess actions relative to the Chebyshev optimum, including no-ops; physical inefficiency is excess path relative to the octile optimum. Both require contact and a positive optimum. In seed 2, global EGP (A) and its segmented shadow (B) had no early successful contact blocks; consequently, four comparisons requiring eight defined pairs were unavailable. The early/late records also contained 22 zero-shortest-path blocks across 12 seed/window rows, for which inefficiency ratios remain undefined. The early contact proportions were 1.6%, 1.2%, 1.6% and 1.4% in A,C,D,E under the 7.5-s horizon.

Exact-test resolution is limited in the larger families, and reached-only path measures describe only successful blocks. The E−C contrast therefore supports improved reward acquisition under this protocol, while effects on path efficiency, cancellation, and biological mediation require separate evidence.

#### S8. Long-duration proposal and consolidation dynamics

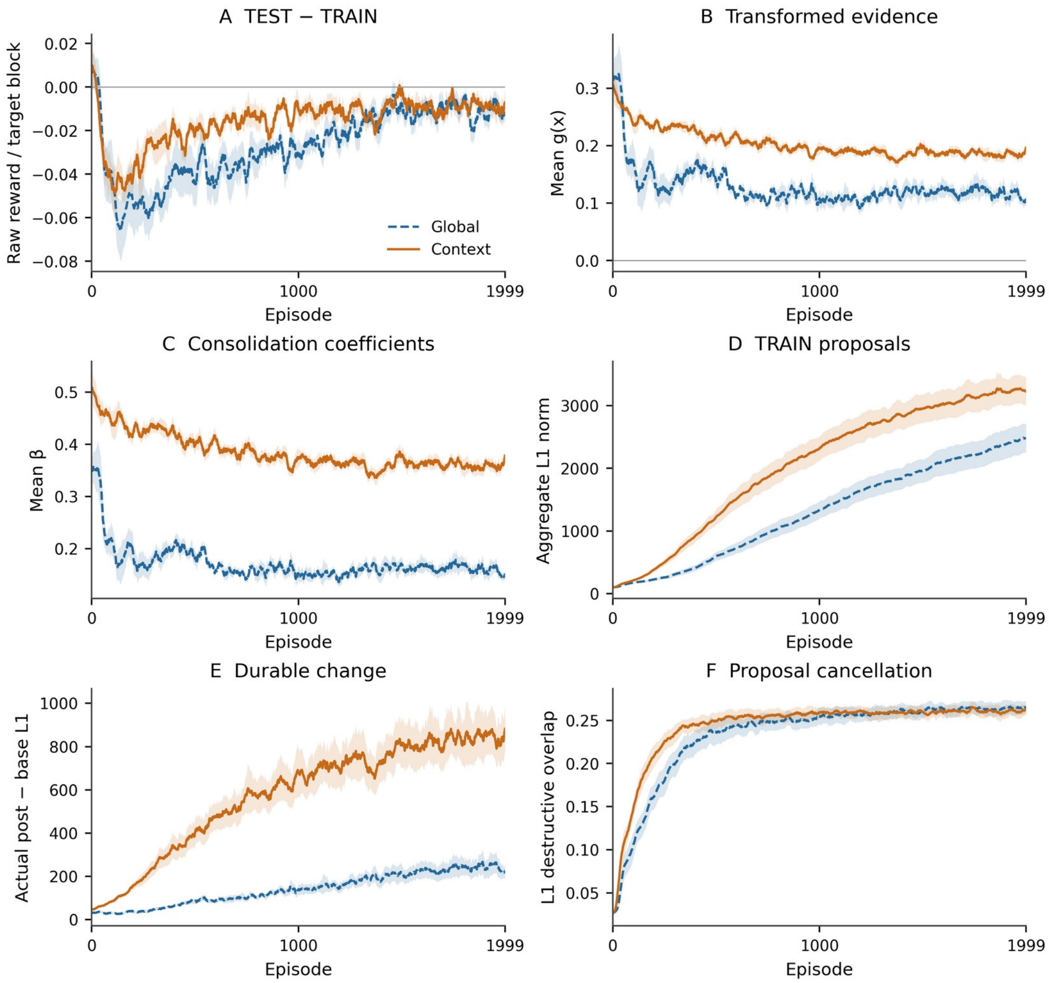


**Figure S11. Raw rewards and internal consolidation quantities during long training.** A) Raw environment TEST−TRAIN reward per target block. The corresponding internal evidence is computed separately by weighting critic samples according to need, calculating their relative TRAIN/TEST change, and applying the nonlinear evidence function. Consequently, a negative raw reward gap can coexist with positive transformed EGP evidence. The raw gap approaches zero while the transformed quantities follow their own dynamics. B) Mean transformed evidence g(x). C) Actual consolidation coefficients β. D) Aggregate TRAIN proposal L1 norm. E) Actual durable-weight change after restoration and clipping. F) L1 destructive cancellation. All 32000 seed/arm/episode rows are retained; curves use the fixed centered 25-episode smoother within seed and shading is between-seed SEM. These curves characterize the internal dynamics across long training; the corresponding component experiment is presented in S7.

Figure S11 follows the internal signals regulating EGP consolidation: raw TEST-minus-TRAIN reward, transformed evidence, consolidation coefficients, accumulated proposals, and durable weight changes. Raw reward is distinct from the need-weighted critic evidence, and proposal magnitude alone does not determine the usefulness of the resulting weight change. The context-specific coefficients summarize five target values, while global EGP has one coefficient. Proposal cancellation and clipping further distinguish the proposed update from the change actually retained in W. Together, these trajectories show how proposal expression, evidence, coefficient scaling, clipping, and consolidation evolve during learning. Cancellation was not reduced in association with the reward advantage; TEST win fractions are summarized descriptively without treating 0.5 as a chance benchmark. Candidate/commit clipping and bound occupancy are shown separately in Fig. S2.

#### S9. Seed-level navigation and computational resources

**Table S15. Success for every paired network.** Internal TEST uses episodes 1900-1999 (2500 target blocks/arm/seed, 30-s horizon). Independent evaluation uses final durable weights (50 fresh-start trials/arm/seed, 187.5-s horizon), with learning and proposals disabled. Values are percentages, with means and ranges describing network-to-network variability. Seed 1, chosen in advance for main Fig. 9, is included with all eight networks.

| Seed | Internal global (%) | Internal context (%) | Independent global (%) | Independent context (%) |
|---|---|---|---|---|
| 1 | 77.20 | 85.40 | 76.00 | 86.00 |
| 2 | 80.96 | 81.64 | 80.00 | 68.00 |
| 3 | 27.64 | 37.00 | 48.00 | 56.00 |
| 4 | 30.60 | 40.04 | 18.00 | 60.00 |
| 5 | 49.24 | 67.20 | 44.00 | 62.00 |
| 6 | 77.72 | 79.52 | 54.00 | 58.00 |
| 7 | 50.84 | 68.48 | 60.00 | 78.00 |
| 8 | 64.56 | 73.76 | 68.00 | 74.00 |
| Mean | 57.35 | 66.63 | 56.00 | 67.75 |
| Range | 27.64-80.96 | 37.00-85.40 | 18.00-80.00 | 56.00-86.00 |

The long-duration simulations were timed using GNU time -v records matched to their completed episode outputs. Each of the 32,000 training episodes simulated 1,500 s using 30 MPI ranks; the measured duration includes process launch, network setup, simulation, and output. Queue delays, controller pauses, SSH setup, and separate post-run verification are excluded. Five unsuccessful or blocked attempts are not included in the completed-episode runtime summaries.

**Table S16. Completed long-duration training episode elapsed wall times in seconds.** Each host contributed 8,000 completed episodes. IQR uses linearly interpolated 25th and 75th percentiles. These values describe observed job wall times on shared systems rather than controlled hardware benchmarks.

| Host / episodes | Median (s) | IQR (s) | Range (s) | Mean (s) |
|---|---|---|---|---|
| Host1 / 8000 | 395.20 | 359.30-468.91 | 325.05-599.17 | 418.58 |
| Host2 / 8000 | 340.43 | 325.25-367.20 | 294.21-716.65 | 367.60 |
| Host3 / 8000 | 328.12 | 311.68-459.79 | 270.57-632.15 | 377.62 |
| Host4 / 8000 | 368.62 | 349.29-396.23 | 308.88-599.84 | 373.25 |
| All hosts / 32000 | 357.15 | 331.81-403.62 | 270.57-716.65 | 384.26 |

Summing the four host capacities, each calculated as 3600/(mean seconds per completed episode), gives approximately 37.6 episodes/hour if every host runs one 30-rank job continuously. Across the 32,000 completed episodes, the summed within-job wall time was approximately 3,416 hours (142.3 host-days), equivalent to about 851 hours (35.5 days) under ideal continuous four-host utilization. The observed campaign duration also reflects shared-host contention, pauses, retries, queue delays, and periods with fewer than four hosts occupied; the idealized time is therefore not a measured calendar duration. SQL queries independently recomputed counts, means, totals, and medians from the controller ledger to verify the reported summaries. These values describe the completed simulations.

Code and supporting analysis materials are available at https://github.com/NathanKlineInstitute/NavNetEGP; the model package is deposited in ModelDB (https://modeldb.science/2041698).

#### S10. Gain controls and matched learning-rule benchmark

To examine the effects of consolidation gain and compare learning rules, we used eight paired network seeds and 200 episodes per condition, with 7,500 actions (375 s) per episode. The conditions were global EGP (A), context-evidence EGP without the extra coefficient multiplier (C), full target-context EGP (E), uniform gain-matched EGP (G), normalized-need EGP (N), shuffled-evidence EGP (S), online STDP/RL (R), and reset-matched STDP/RL (Q). The A/C/E outcomes are those of the component experiment in S7, while G/N/S/R/Q contributed 8,000 additional training episodes. The same seed identities were used across conditions; comparisons using A/C/E are not independent replications of the component study.

##### Coefficient definitions and matching

For each observed target k, $q_k$ is its phase-level exponential mean of raw reward ($\alpha=0.1$). The implemented need equation is $n_k=\mathrm{clip}(1+(\max_j q_j-q_k)/\max(\max_j q_j-\min_j q_j,10^{-9}),0.75,2)$. Its effective range is 1-2; the lower configured bound is inactive. Unseen targets and equal estimates have unit need. When the positive performance range is smaller than $10^{-9}$, the denominator guard limits the increase. All five contexts enter the matching calculation, including contexts with zero proposal.

Context TRAIN and TEST evidence $T_k$ and $E_k$ are mean need-weighted critic samples, distinct from raw environment reward. For $T_k\neq0$, $x_k=(E_k-T_k)/|T_k|$; if $T_k=0$, $x_k=0$ when $E_k=0$ and otherwise $x_k=E_k/\max(|E_k|,10^{-9})$. Let $g(x)=0$ for $x<0$ and $g(x)=2/(1+\exp(-x))-1$ otherwise. With $\eta=1$ and $\varepsilon=0.05$, $b_k=\eta[\varepsilon+(1-\varepsilon)g(x_k)]$ for observed TEST evidence. Absent TEST evidence gives $b_k=0$; the scheduled production data nevertheless require all five target observations. $\varepsilon$ is a consolidation floor in b, not an action-exploration probability.

E: $\beta_k=n_kb_k$;   $c=\Sigma_k n_kb_k/\Sigma_k b_k$;   G: $\beta_k=cb_k$;   N: $\beta_k=n_kb_k/c$;   S: $\beta_k=n_kb\pi(k)$.

If $\Sigma b=0$, c=1 and all G/N coefficients remain zero. No extra floor is applied after N normalization. G preserves the full-rule mean coefficient within G's own state; N preserves the base-coefficient mean within N's own state. These algebraic identities do not imply equal realized coefficient distributions, proposal vectors, cancellation or post-clipping updates across independently trained arms. No arm was calibrated to a chosen reward or durable-L1 outcome.

For S, a fixed SHA-256-indexed schedule selected one of the 44 five-target derangements for each seed, episode and consolidation event, without consuming simulator random draws. It reassigned the base evidence coefficients while keeping target identities, proposal stores and need weights intact. The event-level base multiset was preserved exactly; the final need-weighted β multiset generally was not. Tied values were retained without resampling. All EGP arms expressed the aggregate candidate during TEST, restored durable W, and committed $\mathrm{clip}(W+\Sigma_k\beta_kP_k)$. Target-indexed TEST evidence is therefore not an isolated causal evaluation of $P_k$.

##### Experience, resets, and independent evaluation

The common architecture contained 7168 neurons and 7680 plastic association-to-motor synapses, with paired connectivity/input seeds, initial weights, critic, eligibility and clipping bounds. Each episode started at (65,23). Targets (24,24), (75,75), (24,75), (75,24), and (50,50) repeated in that order in 150-action blocks. EGP used 750-action TRAIN and TEST phases, with five 75-s cycles per episode. TRAIN→TEST restored that TRAIN phase's start; TEST→TRAIN continued from the endpoint. R reset only at episode start. Q also used the EGP within-episode position resets, without proposal/evidence operations. Later reset coordinates can depend on each arm's trajectory, so neither trajectories nor experienced feedback were forced to coincide.

Each episode reconstructed the network using its fixed seeds and imported only the preceding durable weights. Neural state, eligibility and need estimates restarted; this is not continuous-state training across episodes. R and Q committed eligible changes throughout every action interval, including windows corresponding to EGP TEST. The matched budget was 1.5 million environmental interactions per network. Direct durable-update opportunities and synaptic-change magnitude were not matched.

Final episode-199 durable weights were tested with learning and transient proposal expression disabled. A fixed common schedule paired 10 starts for each of five targets within every seed, giving 50 trials per network and 3200 trials overall. Each fresh-start trial lasted 3750 actions (187.5 s). Success required exact active-target contact; failures remained right-censored at the horizon for restricted first-passage means. Conditional time/path endpoints retained their reached-trial denominators. Zero optimal-path or zero occupancy denominators remained NA. This final-only evaluation did not measure a pretraining-to-final improvement or memory decay.

#### Endpoints, inference and missingness

For reward inference, raw action rewards serialized to five decimals were summed within each 150-action target block, averaged across 25 TEST or 50 ALL blocks per episode, then averaged over episodes 0–99 within seed. The network seed, not actions, episodes or trials, was the independent unit. Table S17 retains all ten specified comparisons in the M (Holm-3), L (Holm-4) and C (Holm-3) families. Exact two-sided tests enumerated all 256 sign assignments with absolute tolerance $10^{-15}$. Individual percentile intervals used 100000 paired bootstrap samples from NumPy PCG64 seed 20261030, reset for each contrast. The exact test assumes sign symmetry under the null; its discrete decision and unadjusted percentile interval are not interchangeable. Separate families do not provide experiment-wide family-wise control.

The comparisons and analysis windows were specified before the G/N/S/R/Q simulations. E−C and E−A use the same network realizations as the component experiment and therefore do not constitute additional replications. Late-window, target-specific, mechanistic, and frozen-navigation summaries are descriptive. The 12-test component analysis remains as reported in S7. Undefined endpoints retain their allocated places in the Holm correction; no missing values are replaced by zero. Within-learning reached-only path measures were undefined when the required episode-level ratios were not available and are not plotted. For the frozen-weight comparisons involving R or Q, reached-only timing and path summaries could not be estimated across all eight seeds because one seed in each condition had no successful trials. Exact-target success and all-trial restricted first-passage time remain defined for every seed.

#### Synaptic and activity diagnostics

Actual durable change is the norm of the saved weight-matrix difference across each common 75-s window, averaged over five windows per episode and then over episodes within seed. It is neither the whole-episode net change nor the cumulative sum of individual event increments. L2 is computed from the complete weight vector rather than by summing rank-local norms. Candidate, weighted-proposal, discarded TEST-only, and retained changes remain distinct. Early E/G mean β values were 0.516772/0.514604 and net durable L1 changes were 14.5169/14.5156. Early A/E/R/Q net L1 changes were 8.26627/14.5169/113.889/116.195; the respective L2 changes were 0.892124/2.12545/9.67229/10.0531. These descriptive differences do not establish causal mediation or a plasticity-budget-matched comparison.

L1 target cancellation is $1-||\Sigma P_k||_1/\Sigma||P_k||_1$, with zero denominators undefined; A uses its passive target shadow. Coefficient-weighted and L2 counterparts remain separate numerical diagnostics. Candidate and commit clipping fractions use clipped/eligible synapse-event counts separately; pooled clipping uses their summed numerators divided by summed denominators per episode, followed by the seed-level episode mean. Each complete EGP episode has 38400 eligible events at each stage. R/Q have no EGP candidate/commit endpoint. Their direct-event clipping counts and reward/active-call denominators are separate from EGP clipping. Durable bound occupancy uses the matched saved-weight endpoints. Only EMN, EME, EMS and EMW rates were recorded consistently across arms; other populations are NOT_RECORDED. None of these diagnostics is an additional inferential test.

Table S17. Paired reward comparisons for the consolidation controls and matched STDP/RL benchmark. All endpoints average episodes 0–99 within each of eight paired seeds. M/C use internal TEST; L uses ALL interactions. Differences are first minus second condition, in raw reward per target block. Intervals are unadjusted paired 95% percentile intervals. Exact and adjusted p values are distinct.

| Family | Contrast | Mean difference [95% CI] | Exact p | Holm p |
|---|---|---|---|---|
| M | E−G | 0.002643 [0.000319, 0.005106] | 0.0703125 | 0.140625 |
| M | E−N | 0.007584 [0.004181, 0.010987] | 0.0078125 | 0.0234375 |
| M | E−S | 0.002809 [-0.000826, 0.006693] | 0.21875 | 0.21875 |
| L | E−R | -0.068889 [-0.096785, -0.039539] | 0.0078125 | 0.03125 |
| L | A−R | -0.075183 [-0.105056, -0.043558] | 0.0078125 | 0.03125 |
| L | E−Q | -0.067797 [-0.091636, -0.042992] | 0.0078125 | 0.03125 |
| L | A−Q | -0.074092 [-0.099964, -0.046948] | 0.0078125 | 0.03125 |
| C | E−C | 0.008125 [0.003905, 0.012585] | 0.0078125 | 0.0234375 |
| C | E−A | 0.007648 [0.004313, 0.011456] | 0.0078125 | 0.0234375 |
| C | N−C | 0.000541 [-0.000884, 0.001872] | 0.4765625 | 0.4765625 |

M comprises three mechanism-control contrasts; L four learning-rule contrasts; C three contextual contrasts. E−C and E−A refer to the component-study networks rather than additional independent samples. N−C did not reject (Holm p=0.4765625). The three families retain separate error control. No frozen-navigation or new mechanism p values were added.

Table S18. All 64 seed-level outcomes. Early TEST and ALL reward are means per target block over episodes 0–99. Success is the number of exact-target contacts among 50 final frozen trials, counted once per successful trial. Restricted first-passage time includes all 50 trials, with failures censored at 187.5 s. Durable L1 is the early mean net change per 75-s window. These seed records support the paired analyses and retain weak and zero-success outcomes.

| Arm | Seed | TEST reward | ALL reward | Success | Restricted time (s) | Durable L1 |
|---|---|---|---|---|---|---|
| A | 1 | 0.011015 | 0.001563 | 2/50 | 180.222 | 9.469 |
| A | 2 | -0.001713 | 0.003603 | 10/50 | 163.261 | 2.960 |
| A | 3 | -0.000536 | -0.007488 | 9/50 | 175.170 | 8.120 |
| A | 4 | 0.009846 | -0.001158 | 6/50 | 175.915 | 9.860 |
| A | 5 | 0.005535 | -0.000387 | 12/50 | 162.861 | 13.157 |
| A | 6 | -0.009219 | -0.020398 | 8/50 | 168.201 | 11.645 |
| A | 7 | 0.002710 | -0.006285 | 7/50 | 169.672 | 6.694 |
| A | 8 | 0.000769 | -0.000242 | 8/50 | 178.316 | 4.224 |
| C | 1 | 0.000158 | -0.004779 | 6/50 | 172.192 | 5.720 |
| C | 2 | 0.001500 | 0.003774 | 5/50 | 174.229 | 8.802 |
| C | 3 | -0.000257 | -0.007577 | 7/50 | 169.611 | 5.673 |
| C | 4 | 0.011614 | -0.003392 | 5/50 | 174.111 | 7.973 |
| C | 5 | 0.008007 | 0.000269 | 8/50 | 169.936 | 9.624 |
| C | 6 | -0.008434 | -0.019516 | 5/50 | 175.534 | 11.655 |
| C | 7 | 0.001422 | -0.008285 | 5/50 | 177.433 | 6.864 |
| C | 8 | 0.000580 | -0.002683 | 3/50 | 183.233 | 7.284 |
| E | 1 | 0.016824 | 0.008427 | 7/50 | 173.613 | 12.570 |
| E | 2 | 0.001918 | 0.005467 | 7/50 | 170.665 | 13.037 |
| E | 3 | 0.000848 | -0.007913 | 10/50 | 162.785 | 9.027 |
| E | 4 | 0.015235 | 0.002497 | 3/50 | 178.597 | 11.615 |
| E | 5 | 0.021904 | 0.012787 | 3/50 | 179.856 | 18.248 |
| E | 6 | -0.003353 | -0.015406 | 2/50 | 180.658 | 19.462 |
| E | 7 | 0.008912 | 0.000165 | 7/50 | 168.866 | 17.114 |
| E | 8 | 0.017301 | 0.013541 | 4/50 | 179.473 | 15.062 |
| G | 1 | 0.019324 | 0.011821 | 1/50 | 183.942 | 13.233 |
| G | 2 | 0.001123 | 0.003051 | 7/50 | 165.943 | 14.472 |
| G | 3 | 0.000410 | -0.006702 | 13/50 | 156.748 | 8.355 |
| G | 4 | 0.008478 | -0.004760 | 7/50 | 170.082 | 11.766 |
| G | 5 | 0.019244 | 0.009854 | 5/50 | 174.416 | 19.496 |
| G | 6 | -0.003433 | -0.014649 | 8/50 | 168.367 | 18.229 |
| G | 7 | 0.004653 | -0.003535 | 3/50 | 182.580 | 15.485 |
| G | 8 | 0.008642 | 0.004701 | 5/50 | 177.237 | 15.089 |
| N | 1 | 0.002162 | -0.003954 | 3/50 | 177.123 | 5.757 |
| N | 2 | 0.001661 | 0.004354 | 5/50 | 174.412 | 7.936 |
| N | 3 | -0.001766 | -0.009433 | 9/50 | 164.391 | 5.362 |
| N | 4 | 0.011660 | -0.002435 | 4/50 | 175.276 | 8.042 |
| N | 5 | 0.011025 | 0.004702 | 5/50 | 176.384 | 9.955 |
| N | 6 | -0.011536 | -0.023247 | 3/50 | 180.338 | 11.475 |
| N | 7 | 0.002110 | -0.007088 | 6/50 | 176.333 | 7.665 |
| N | 8 | 0.003601 | 0.000169 | 7/50 | 178.068 | 7.666 |
| S | 1 | 0.004613 | -0.001450 | 2/50 | 185.763 | 11.264 |
| S | 2 | -0.002173 | 0.000040 | 3/50 | 180.920 | 13.577 |
| S | 3 | -0.001682 | -0.008845 | 11/50 | 166.619 | 10.191 |
| S | 4 | 0.014875 | 0.001531 | 7/50 | 174.072 | 13.641 |
| S | 5 | 0.012675 | 0.005918 | 7/50 | 172.595 | 18.308 |
| S | 6 | 0.000578 | -0.007273 | 5/50 | 174.030 | 20.340 |
| S | 7 | 0.006515 | -0.002161 | 4/50 | 179.385 | 15.510 |

| Arm | Seed | TEST reward | ALL reward | Success | Restricted time (s) | Durable L1 |
|---|---|---|---|---|---|---|
| S | 8 | 0.021719 | 0.017875 | 2/50 | 181.860 | 15.027 |
| R | 1 | 0.121670 | 0.108110 | 2/50 | 180.169 | 95.103 |
| R | 2 | 0.060858 | 0.070680 | 2/50 | 180.125 | 156.006 |
| R | 3 | 0.001962 | -0.002139 | 7/50 | 172.963 | 68.573 |
| R | 4 | 0.028723 | 0.026285 | 2/50 | 180.474 | 85.597 |
| R | 5 | 0.131389 | 0.121164 | 2/50 | 180.214 | 159.183 |
| R | 6 | 0.109567 | 0.106879 | 0/50 | 187.500 | 144.640 |
| R | 7 | 0.042001 | 0.029186 | 1/50 | 183.792 | 95.614 |
| R | 8 | 0.109028 | 0.110511 | 2/50 | 180.128 | 106.399 |
| Q | 1 | 0.106748 | 0.091553 | 1/50 | 183.861 | 90.825 |
| Q | 2 | 0.062497 | 0.068158 | 0/50 | 187.500 | 156.190 |
| Q | 3 | 0.013538 | 0.005410 | 6/50 | 171.916 | 87.935 |
| Q | 4 | 0.037282 | 0.029795 | 2/50 | 180.744 | 82.484 |
| Q | 5 | 0.124351 | 0.112315 | 2/50 | 180.528 | 148.350 |
| Q | 6 | 0.104593 | 0.099990 | 2/50 | 180.174 | 154.484 |
| Q | 7 | 0.056311 | 0.040909 | 5/50 | 180.068 | 100.917 |
| Q | 8 | 0.116073 | 0.113816 | 4/50 | 172.889 | 108.379 |

A: global EGP; C: context-evidence EGP; E: full target-context EGP; G: uniform gain matched; N: normalized need; S: shuffled evidence; R: online STDP/RL; Q: reset-matched STDP/RL. For R/Q, TEST is an analysis window only. Each row is one independent network realization per condition. Table S18 does not add tests, and repeated trials within a network are not independent network replicates.

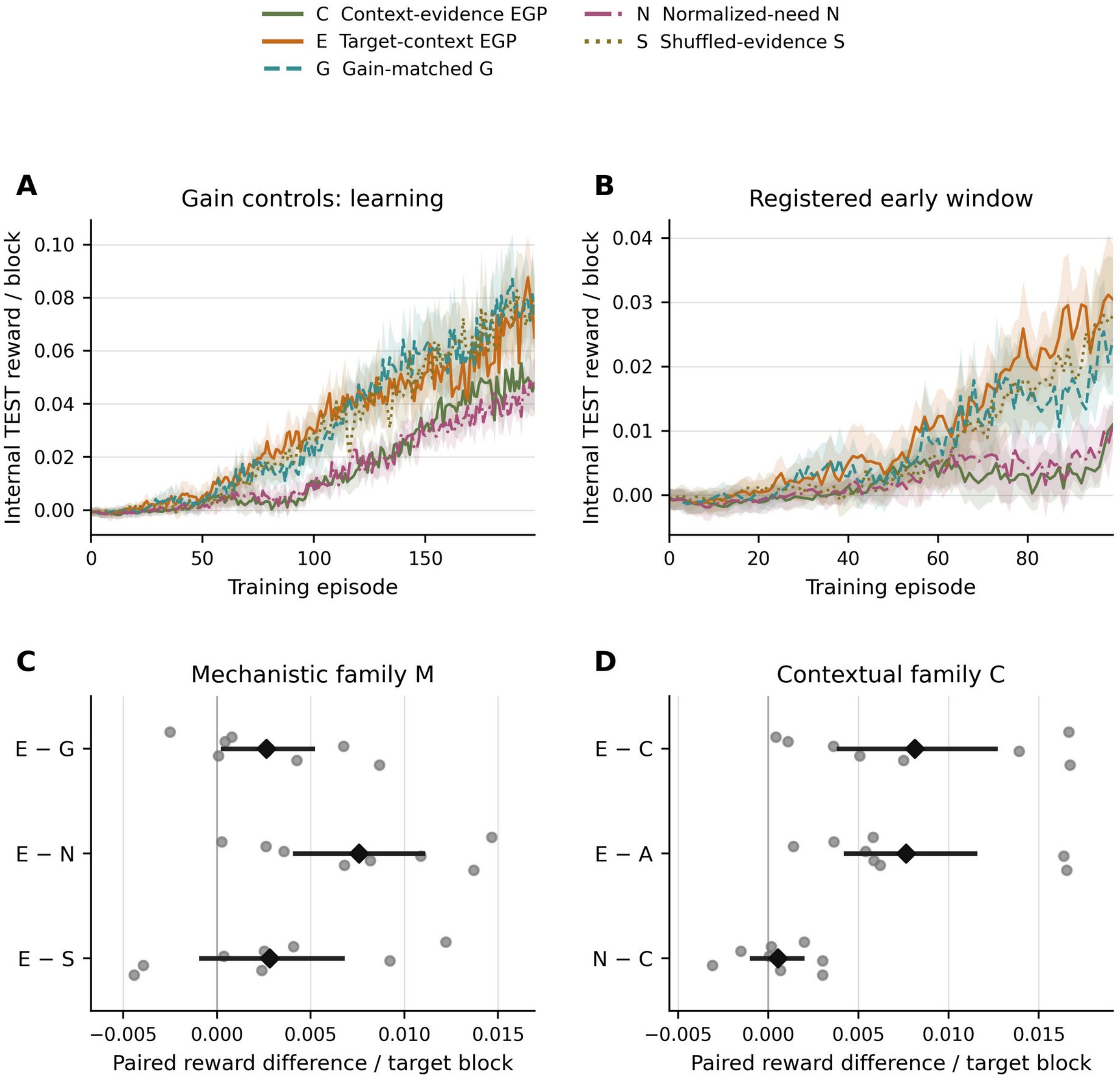


Figure S12. Reward acquisition and gain-control contrasts. A,B) Unsmoothed mean ± between-seed SEM for the complete and early internal-TEST learning curves; TEST temporarily expresses aggregate proposals. C,D) All eight seed differences (gray points), means (diamonds), and unadjusted 95% paired-bootstrap intervals for the complete M and C families. Exact and Holm-adjusted p values are in Table S17; interval exclusion of zero does not override the adjusted test. E−C and E−A use the component-study networks and do not add independent replications.

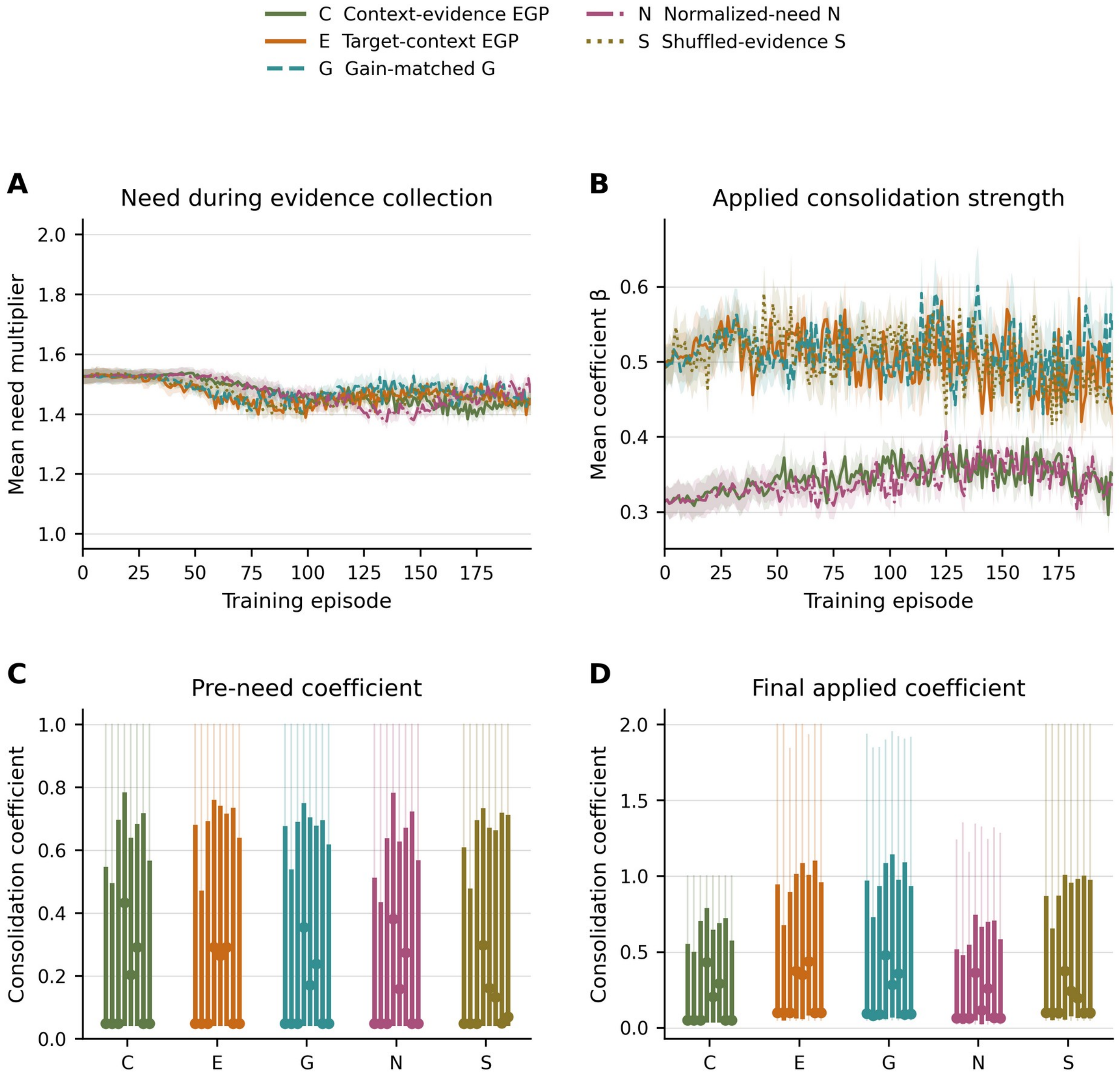


Figure S13. Need scaling and consolidation coefficients. A,B) Unsmoothed mean ± between-seed SEM over 200 episodes. The implemented need formula yields values between 1 and 2; its configured 0.75 lower bound is inactive. C,D) Early within-seed distributions: thin line, full range; thick line, interquartile range; dot, median. Each seed includes 2500 repeated target-events across episodes 0–99. S preserves the event-level multiset of pre-need base coefficients, but multiplying by the target-specific need weights generally changes the final β multiset. G and N match counterfactual means within their own state; independently evolving arms need not match exactly.

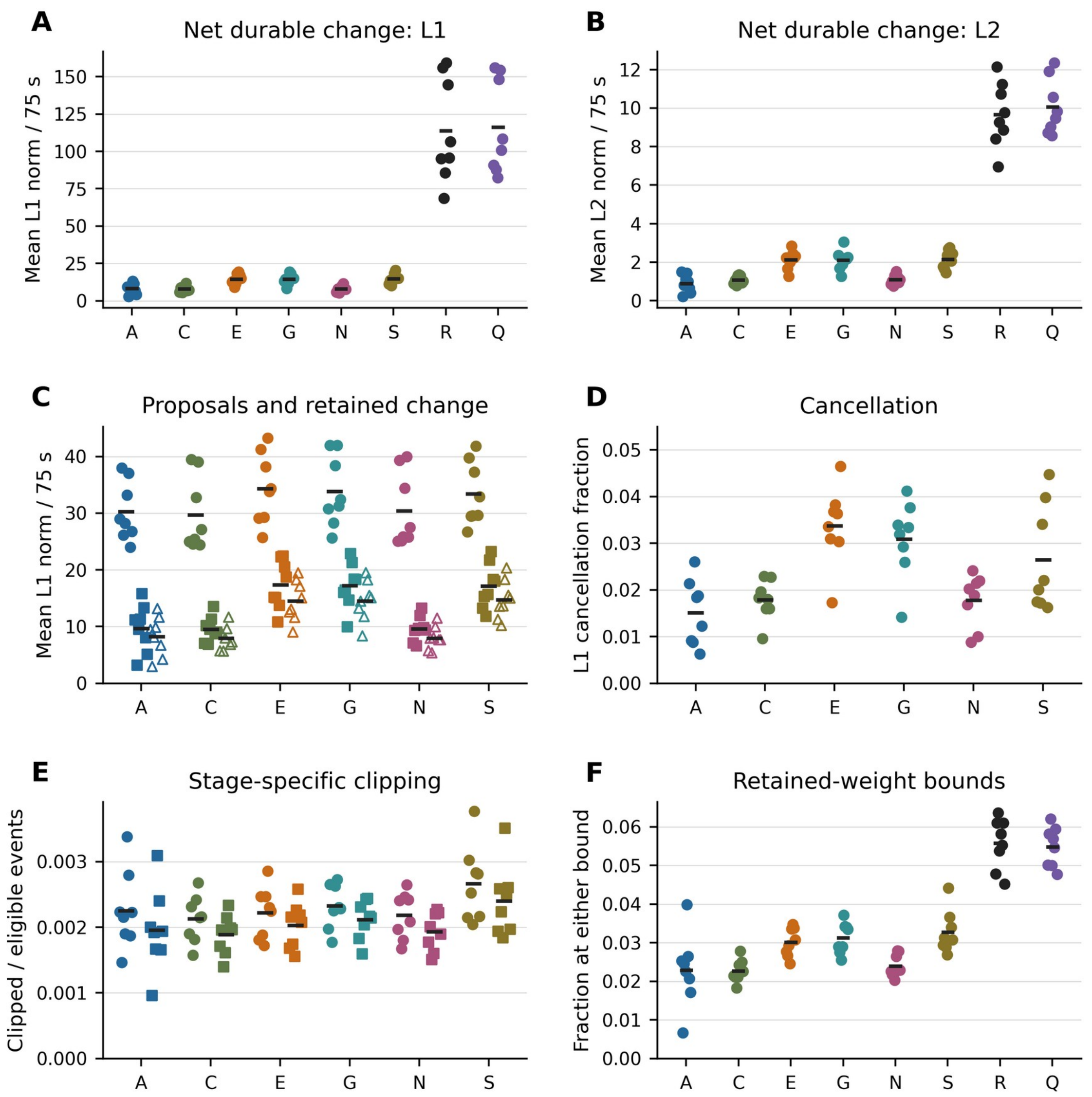


Figure S14. Proposed updates and retained synaptic changes in episodes 0–99. A,B) Net durable weight-matrix L1 and L2 changes across matched 75-s windows; these are not sums of individual event updates. C) Pooled proposal (circles), coefficient-weighted proposal (squares), and retained change after clipping (open triangles), in L1 units. D) Destructive L1 overlap among target proposals; global A uses its passive target shadow. E) Candidate (circles) and commit (squares) clipping fractions, with separate eligible-event denominators. F) Durable-weight occupancy at either instantiated bound. Points are all eight seed endpoints and black marks are strict means. Episode means average five windows and seed endpoints average 100 episodes. Candidate/proposal quantities do not apply to R/Q. All diagnostics are descriptive; fuller numerical records retain L2 cancellation, pooled clipping, and late-window results.

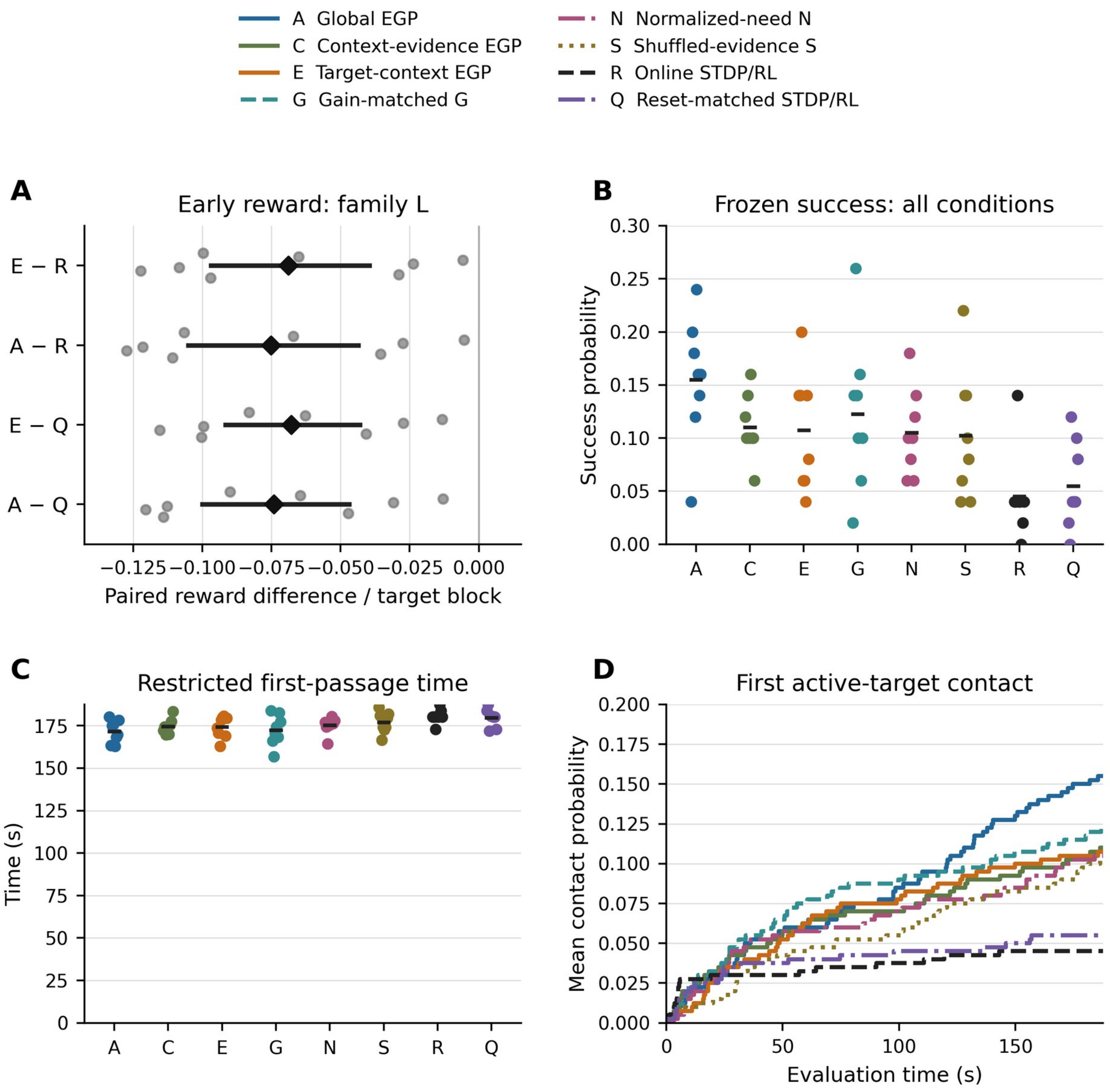


Figure S15. Matched baseline reward and independent navigation across all eight conditions. A) The four specified early all-interaction reward differences, with eight paired seed differences, means, and unadjusted 95% bootstrap intervals; all Holm-4 p values are 0.03125 (Table S17). B,C) Seed success and restricted mean first-passage time after 200 training episodes; black marks are strict means. D) Mean cumulative first-contact curves, with right-continuous steps and failures censored at 187.5 s. Each of 64 networks contributed 50 trials on the common paired schedule (3200 trials). Panels B–D are descriptive. Reached-only quantities retain conditional denominators and are not substituted for the all-trial measures.

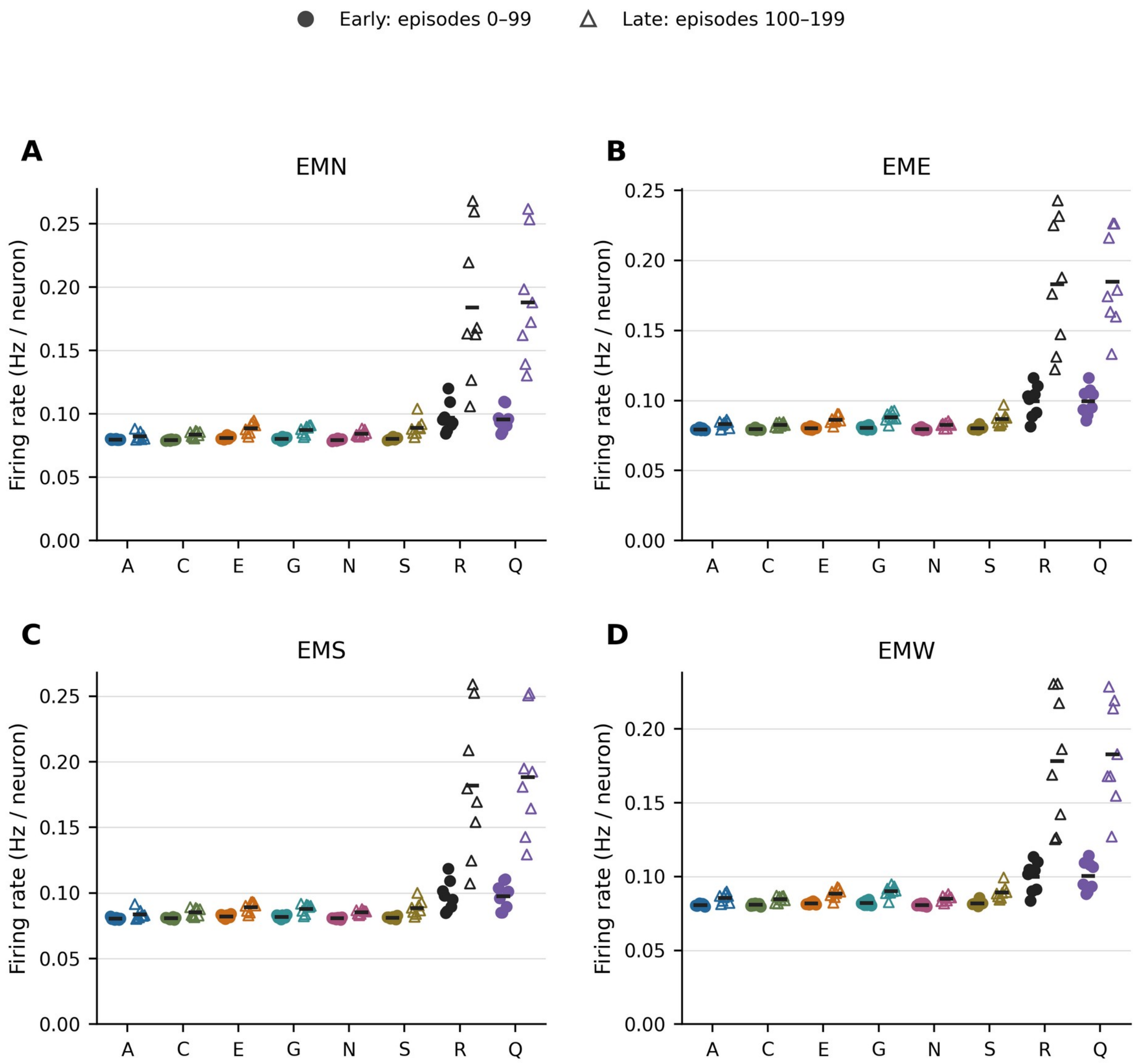


Figure S16. Recorded motor-population activity during learning. Mean all-interaction firing rates per neuron for EMN, EME, EMS, and EMW. Filled circles show early seed means (episodes 0–99), open triangles late seed means (100–199), and black marks strict arm means. Each arm retains all eight network seeds. Only these four motor populations were recorded consistently across conditions; firing rates for other populations were not consistently available and are not interpreted as zero. These descriptive activity summaries are not tests of physiological realism.
